# Auxetic Two-Dimensional Nanostructures from DNA

**DOI:** 10.1101/2020.08.21.262139

**Authors:** Ruixin Li, Haorong Chen, Jong Hyun Choi

## Abstract

Architectured materials exhibit negative Poisson’s ratios and enhanced mechanical properties compared with regular materials. Their auxetic behaviors should emerge from periodic cellular structures regardless of the materials used. The majority of such metamaterials are constructed by top-down approaches and macroscopic with unit cells of microns or larger. On the other extreme, there are molecular-scale auxetics including naturally-occurring crystals which are not designable. There is a gap from few nanometers to microns, which may be filled by bottom-up biomolecular self-assembly. Here we demonstrate two-dimensional auxetic nanostructures using DNA origami. Structural reconfiguration experiments are performed by strand displacement and complemented by mechanical deformation studies using coarse-grained molecular dynamics (MD) simulations. We find that the auxetic properties of DNA nanostructures are mostly defined by geometrical designs, yet materials’ chemistry also plays an important role. From elasticity theory, we introduce a set of design principles for auxetic DNA metamaterials, which should find diverse applications.

## INTRODUCTION

Architectured auxetic materials have received significant attention due to their exotic behaviors and properties. They exhibit a negative Poisson’s ratio (*ν*) and possess significantly improved indentation resistance^1^, greater shear modulus^2^, and enhanced fracture toughness^3^ compared with regular materials. These characteristics are believed to emerge from their periodic cellular structures rather than materials and related chemistries, forming the basis of mechanical metamaterials.^4^ Most of auxetic architectures are manufactured by top-down approaches using metals^5^, polymers^6^, and fibers^7^. Their unit cells range from microns to centimeters, forming macroscopic bulk materials. On the other extreme, there are molecular-scale auxetics, including α-cristobalite and cubic metals, whose unit cell sizes are on the order of few chemical bond lengths.^8–10^ These naturally-occurring crystals are not designable. Synthetically, metal-organic frameworks (MOFs) have been theorized as auxetic building blocks, yet the concept has not been demonstrated clearly through experiments.^11,12^ Thus, there is a wide gap from few nanometers to microns for well-defined auxetic unit structures. As part of bottom-up approaches, biomolecular self-assembly is ideal to create custom-designed auxetic nanoarchitectures, filling the gap. Further, such nanostructures could also serve as a platform to examine any possible effects of materials on auxetic behaviors and mechanical properties at a very small lengthscale, which have been difficult to study to date.

Here we demonstrate auxetic two-dimensional (2D) nanostructures using wireframe DNA origami and elucidate relevant mechanics. DNA origami is a widely used self-assembly method for creating arbitrary nanoarchitectures with excellent programmability and structural predictability.^13–16^ We constructed DNA nanostructures, composed of one or few unit cells, and achieved their auxetic transformations via strand displacement. Atomic force microscopy (AFM) measured the origami conformations, from which Poisson’s ratios were calculated. Coarse-grained molecular dynamics (MD) simulations were performed to observe mechanical deformations upon compressive loading, extracting structural properties. We found that the auxetic behaviors are largely defined by geometrical designs, but materials also play a role, which shall not be ignored at this very small scale. Based on elastic beam theory, we developed a model that accounts for edge rigidity and joint flexibility, yielding a set of guiding principles for auxetic DNA metamaterials.

### Auxetic vs. Regular Structures

To demonstrate DNA auxetics, we chose a well-studied, re-entrant honeycomb structure^17^. When expanding (or shrinking) in one direction, this periodic structure will react the same way in the orthogonal direction, as illustrated by (i)–(iii) in Figure 1a. In contrast, its conjugate, regular honeycomb, will do the opposite ((iii)–(v) in Figure 1a). In our design, each origami includes four intact unit cells (bold lines in Figure 1a), where edges and multi-arm joints at vertices form a wireframe structure.^18–21^ Specifically, two double-stranded DNA (dsDNA) helices form an edge (Figure 1b, black), and single-stranded DNA (ssDNA) segments are used for the joint (Figures S2, S3, also see SI for design details). In addition, we added ‘jack’ edges shown in red in Figure 1b whose length can be changed to modulate the angle (*γ* indicated by the red dots) and have different conformations. For example, the structure is largest at 90° with the jacks fully stretched. As the jack length is shortened to make the angle 60° and then 30°, both the height and width, represented by *y*_1_ and *x*_1_, respectively, become shorter, thus resulting in a shrinkage of the entire structure ((i)–(iii) in Figure 1b). The Poisson’s ratio may be expressed as a function of *γ*,

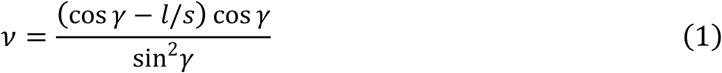

where *l* and *m* are lengths of edges. Equation 1 is plotted as the black curve in Figure 1c. *ν* is 0 at 90° and marches toward negative and positive infinity at *γ* = 0° and 180°, respectively.

**Figure 1.**
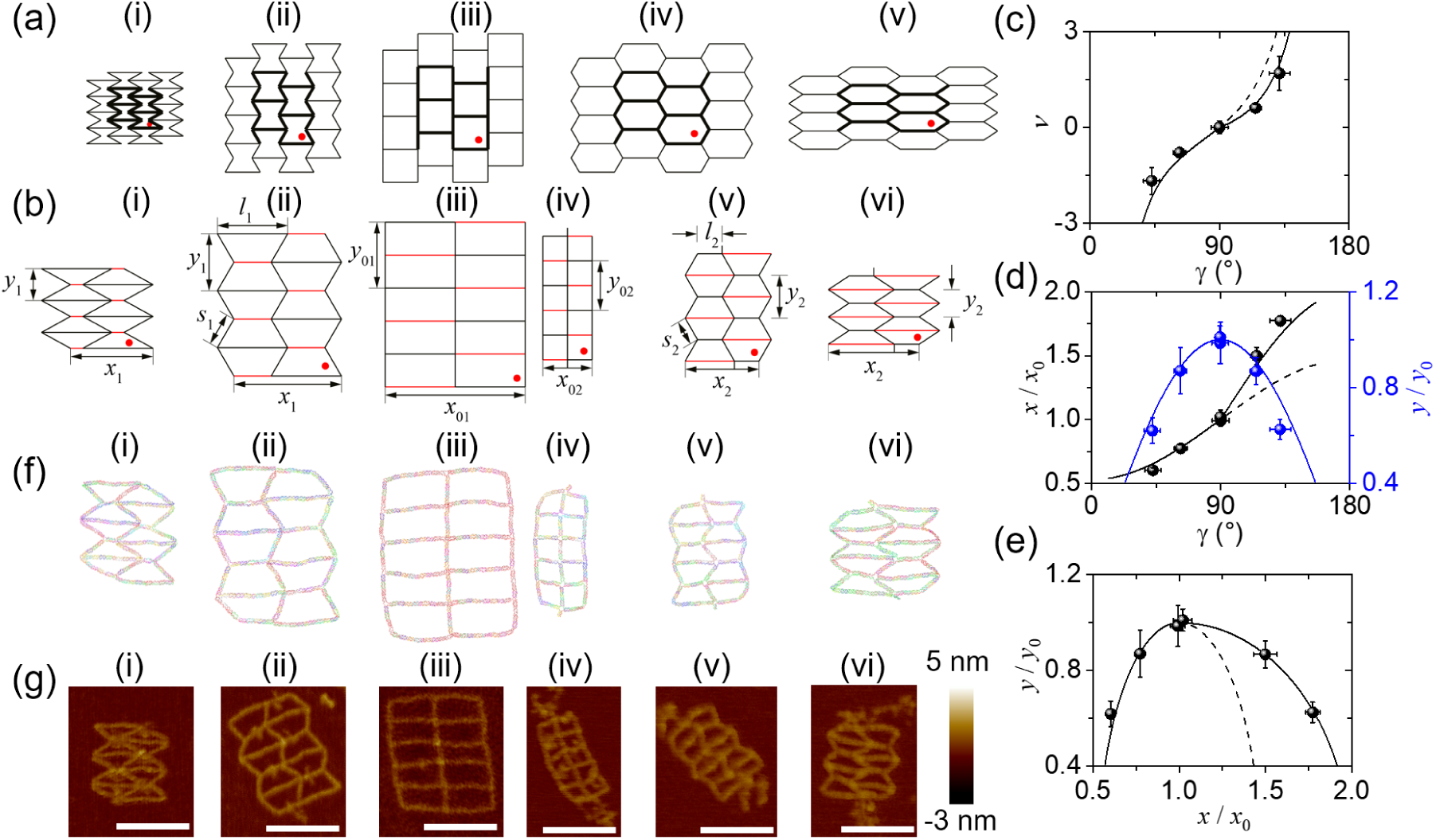
Re-entrant and regular honeycombs. (a) Schematics of the honeycombs in different conformations. The angles (*γ*), noted by red dots, are 30° (i), 60° (ii), 90° (iii), 120° (iv), and 145° (v). (b) Designs of honeycombs including four intact unit cells. The black lines represent the regular edges, while jack edges are shown in red. The coordinates, *x* and *y*, and the edges, *l* and *s*, are also noted. Subscript 1 indicates re-entrant structures ((i)–(iii)), whereas 2 represents regular honeycombs ((iv)–(vi)). Subscript 0 denotes the reference states ((iii) and (iv), *γ* = 90°). (c)–(d) Poisson’s ratio (*ν*), *x*/*x*_0_, and *y*/*y*_0_ as a function of *γ*. (e) *x*/*x*_0_ versus *y*/*y*_0_. Solid lines represent theoretical predictions of re-entrant honeycombs for *γ* ≤ 90° (*x*/*x*_0_ ≤ 1.0) and regular structures for *γ* ≥ 90° (*x*/*x*_0_ ≥ 1.0). The dashed curves represent theoretical predictions of the regular structures that would be imaginary conjugates of the re-entrant honeycombs, if elongated jack edges were used. (f) Equilibrium conformations of DNA origami honeycombs from coarse-grained MD simulations. (g) AFM images of DNA origami structures corresponding to the designs in (b) and the simulations in (f). Unused scaffold and jack segments slightly blur the images. Scale bar: 100 nm. The experimental data are also shown as filled circles with error bars in (c)–(e).

To induce positive *ν*, the jacks must be elongated, forming the regular honeycomb. Due to the size limitation of the scaffold (M13mp18), we designed another, smaller regular honeycomb that can stretch as the jack length increases and the angle transforms from 90° to 120° and then to 145° as shown in (iv)–(vi) of Figure 1b (also see Figure S2, S3). Our design includes joints with different lengths of ssDNA for a range of *γ* (Tables S3.1, S3.2). For example, 6 nucleotides (6-nt) were used at a joint for 90–145°. The ssDNA joint provides a degree of freedom and flexibility for structural transformation. Since the scales of the re-entrant and regular honeycomb structures are different, dimensionless width and height (*x*/*x*_0_ and *y*/*y*_0_) are plotted as a function of *γ* with the reference states (*x*_0_ and *y*_0_) set at *γ* = 90° (Figure 1d). It is evident that the width increases monotonically with *γ*, while the height shows opposite trends for negative and positive *γ*. The height vs. width plot in Figure 1e shows that the height of the re-entrant structure increases with expanding width, whereas it decreases in the regular honeycomb.

MD simulations were performed based on the coarse-grained model using oxDNA^22,23^ to estimate the origami conformations. The structures with different jack edges were created in caDNAno2^24^ and subsequently converted into oxDNA files for initial conformations. Each conformation was then set at 4 °C to reach an equilibrium state in the simulation. Figure 1f presents the equilibrium conformations where all the unit cells show symmetry within themselves and between neighboring units. The simulated structures reasonably follow the corresponding designs in Figure 1b. Origami structures were synthesized and visualized by AFM (see SI for experimental details and additional AFM images). The AFM images look alike the corresponding designs and simulations (Figure 1g, Figure S29–S34). In the structures with short jacks, the unused scaffold segments also show up in the AFM images, somewhat interfering with the geometrical designs. Measurements on the experimental results are plotted as filled circles in Figure 1c–e, along with the theoretical calculations (Figures S8). The computational and experimental results suggest the feasibility of the origami-based approach for nanoscale auxetics.

### Structural Transformation via Chemical Stimuli

To demonstrate auxetic reconfiguration, we used a simpler, re-entrant triangle structure (Figure 2a). The re-entrant triangle may be considered as a reduced structure from the re-entrant honeycomb where two edges are shrunk to a point (see Figure S4). It thus behaves alike the re-entrant honeycomb and can display simultaneous expansion (or compression) in two orthogonal directions. In our design, the re-entrant triangle origami has quadrilateral unit-cells (Figure 2b), where each edge is made of two dsDNA and every joint consists of ssDNA segments (Figure S5). The jack edges, shown in red, are also included to enable structural deformation. Despite the structural simplicity, it is still difficult to apply mechanical loads accurately onto nanostructures.^25^ Hence, we used chemical loads instead to modulate the jack length through toehold-mediated strand displacement,^26–28^ thereby resulting in reconfiguration, either expansion or shrinkage. For example, the re-entrant triangle can switch between 30° and 58° as shown in Figure 2b, by modulating the jack length of approximately 45 or 59 nm (6 or 8 full turns), respectively.

**Figure 2.**
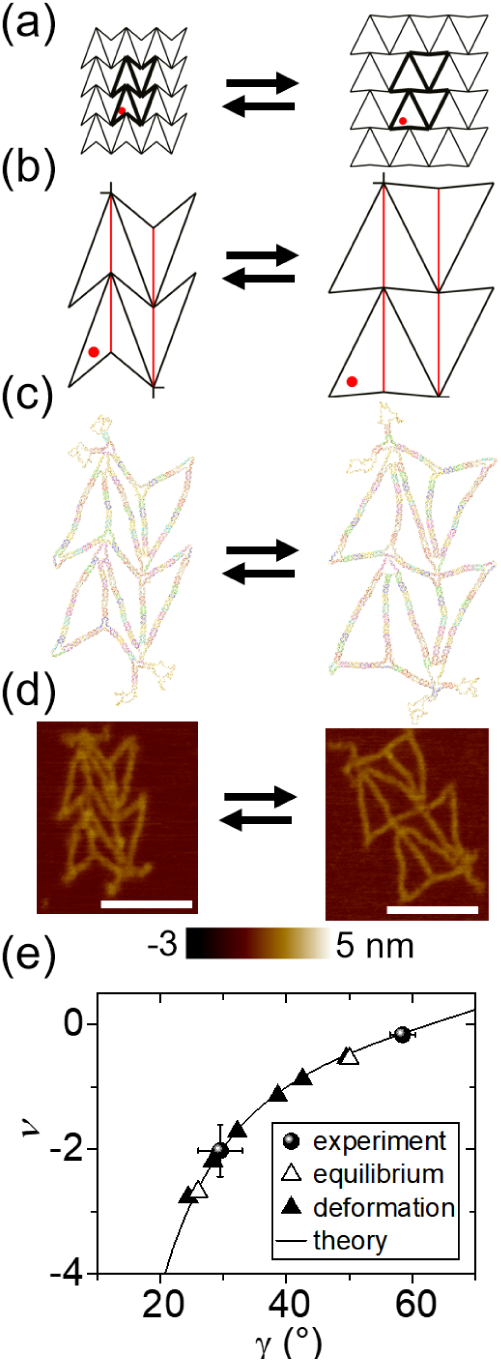
Structural transformation of the re-entrant triangle via chemical stimuli. (a)–(b) Schematics and designs in different conformations. The red dot indicates the angle *γ*. The black lines denote the regular edges, while jack edges are shown in red. (c) Equilibrium conformations of DNA origami from oxDNA simulations. (d) AFM images of DNA origami after reconfiguration. Unused scaffold and jack segments slightly blur the images. Scale bar: 100 nm. (e) Poisson’s ratio versus angle. Solid circles represent the experimental data with error bars. The empty triangles show the MD simulations of equilibrium structures, while the filled triangles denote the simulated deformation data. The line represents the theoretical prediction based on the finite model.

Figure 2c–d shows the re-entrant triangle structures in two distinct states from MD simulations and AFM measurements (Figures S35, S36). Experimentally, we first added invading strands (*i*.*e*., releasers) to remove the staples from the jack edges initially in an equilibrium, followed by addition of another set of staples to form the jacks with a different length. As a result, the triangle can transform from 58° to 30°, and *vice versa*. Figure 2e presents the angle dependence of *ν* based on the measurements from experiments and simulations. Although the computational and experimental results generally follow the designs, curved edges and blunt joints are quite noticeable. These issues are caused by the design model which fundamentally assumes infinitely rigid edges and infinitely flexible joints (termed *infinite* model), as in typical architectured metamaterials studies. They may be more pronounced at the nanoscale where materials properties must be accounted for.

### Mechanical Deformation in MD Simulations

Next, we performed mechanical deformations using MD simulations, where external forces were applied directly on the re-entrant triangle origami. In the equilibrium structure, the jack edges were cut off to free up space which was subsequently subject to loading (Figure 3a). The moving harmonic trap in the oxDNA was used to pull the jack residues in from *γ* = ∼50° until the structure reaches ∼26° (Figure 3b, Figure S26). During the deformation, forces and displacements were recorded and converted into stress-strain, from which Young’s modulus *E* in the loading direction was calculated (Figure 3c, filled triangles). The computational results show the angle dependence, and *E* is on the order of few pN/nm^2^, significantly lower than that of dsDNA (∼100 pN/nm^2^).^29,30^ The small Young’s modulus is common in cellular structures, especially due to ssDNA (∼1 pN/nm^2^) at the joints,^31^ serving as the determinant of *E*. To better understand the mechanics, we developed a model that accounts for finite rigidity in edges and finite flexibility in joints (which we term *finite* model; see the SI for details). The model includes three deformation modes, flexing (*K*_*f*_), stretching (*K*_*s*_), and hinging (*K*_*h*_), which all resemble the Hooke’s law^32^:

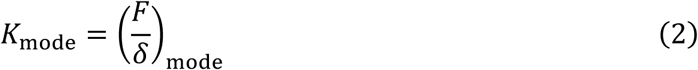

Among the force constants, *K*_*f*_ and *K*_*h*_ are on the same order of magnitude, while *K*_*s*_ is much larger than *K*_*f*_ and *K*_*h*_. The ratio of stretching and flexing force constants (*K*_*s*_/*K*_*f*_) is about 300 and 70 for edge *l* and *m*, respectively. Therefore, it is much more difficult to stretch or compress an edge than to bend or rotate it. The primary mode that accounts for the structural deformation can change as the angle changes.

**Figure 3.**
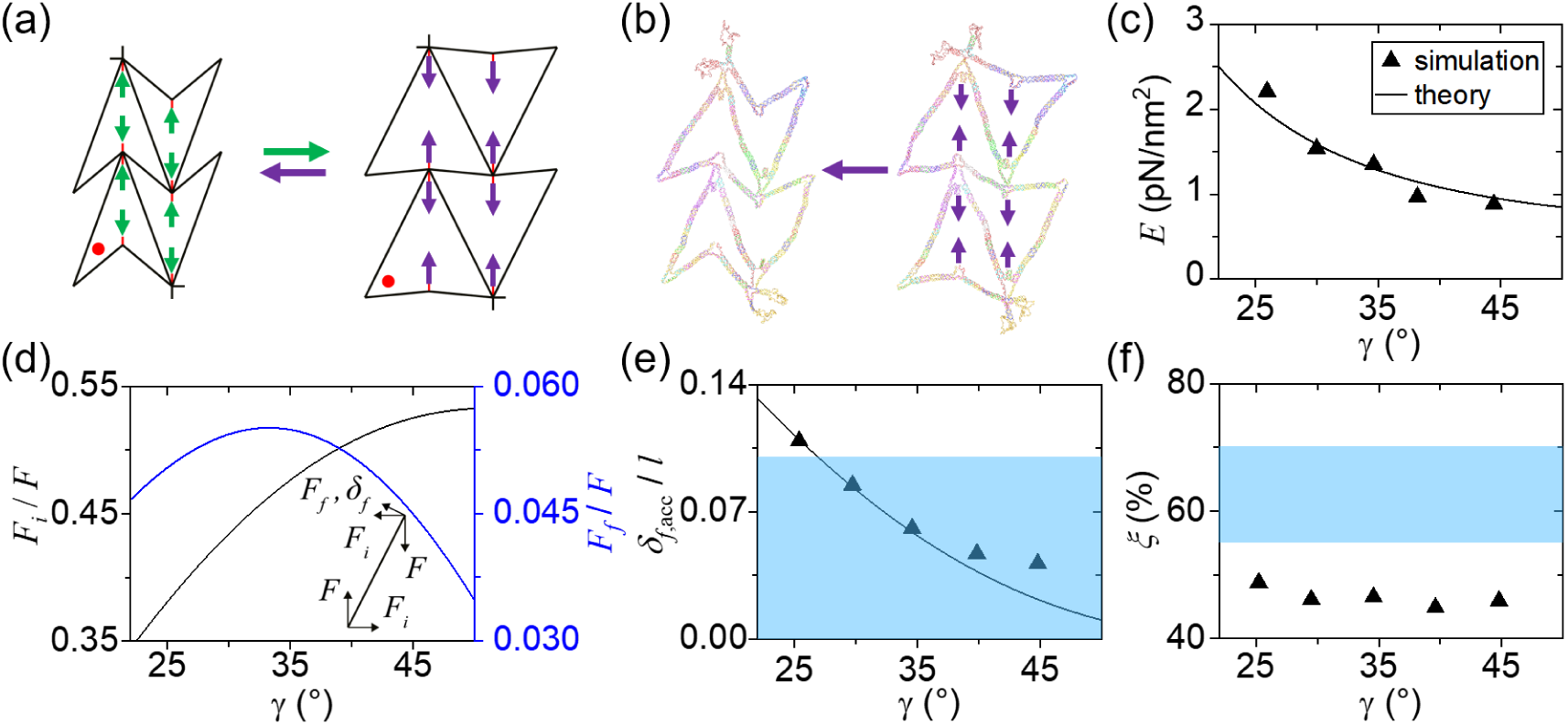
Mechanical deformation of the re-entrant triangle in MD simulations. (a) Schematics of the simulation domain with the jack edges removed. The compressive and tensile forces are shown in purple and green, respectively. (b) Simulation results of the re-entrant triangle under compressive loads, reconfiguring from *γ* = ∼50° to ∼26°. (c) Young’s modulus (*E*) versus angle *γ. E* increases moderately as the triangle folds, indicating that it becomes harder to compress. (d) Dimensionless internal horizontal force (*F*_*i*_/*F*) and the force responsible for flexure (*F*_*f*_/*F*) as a function of *γ*. The inset shows the edge *l* with the forces and flexure. (e)–(f) Dimensionless total flexure (*δ*_*f*,acc_/*l*) and joint stretch (*ξ*). The finite model (black line) reasonably predicts the simulated deformation data (solid triangles). Blue shades indicate the design requirements. The flexure exceeds the recommended range at small angles (< 27°). The stretch level is ∼45%, indicating that the joints are too loose, which results in structural inaccuracy.

The displacement in the loading direction Δ*x* can be expressed as a sum of displacements in different modes (*δ*_mode_) projected along the loading direction, yielding the relationship between the force and displacement at any given angle. The accumulated, total force (*F*_acc_) that transforms the origami from an initial (*γ*_*i*_) to a final angle (*γ*_*f*_) is calculated by integration. With knowledge of *F*_acc_ and Δ*x*, the stress (*σ*_acc_) and the strain (*ε*_1,acc_) are calculated, from which *E* is obtained.

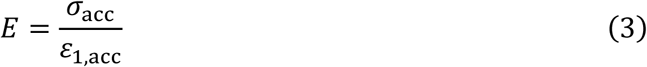

This model predicts the angle dependence of *E* (Figure 3c). As *γ* decreases, for example, *E* increases, implying that it is harder to compress the structure as it deforms. The stretching mode becomes a dominant deformation mode as the triangle narrows. The theoretical prediction is in excellent agreement with the data from MD calculations. It should be noted that the finite model prediction also matches well with the experiment and simulation results in Figure 2e. Moreover, it shows that even though the components are assumed to have linear elasticity, the overall elastic properties can be nonlinear due to the cellular geometry.

### Design Principles

The finite model also allowed us to study curved edges. Upon external loads (*F*), an internal horizontal force (*F*_*i*_) is generated to hold the edges in place against falling apart (inset in Figure 3d). As the compressive loading continues, the ratio of forces *F*_*i*_/*F* decreases, because the horizontal force becomes less prominent as the triangles fold to smaller angles. The force responsible for flexure of the edge *l* (*F*_*f*_) is calculated by projecting *F*_*i*_ and *F* onto the flexure (*δ*_*f*_) direction. *F*_*f*_ increases as *γ* shrinks due to a better alignment of *F*_*i*_ with *δ*_*f*_. As the triangle further narrows, however, *F*_*i*_ decreases significantly so that *F*_*f*_ also drops. *F*_*f*_ is positive at all angles, suggesting that the edge will bend continuously to the *δ*_*f*_ direction. By integrating the flexure with respect to angle, we obtain *δ*_*f*,acc_/*l* as a function of *γ* as shown in Figure 3e, where the dimensionless total flexure increases monotonically as the structure deforms. Notably, Figure 3e shows significant flexure, particularly at small angles (*γ* < 27°), suggesting a compromised structural integrity. To suppress flexure, we propose that *δ*_*f*_ should be less than maximum thickness of the edge (*t*_max_). To simplify, we consider the loads that contribute directly to the flexure and the elastic range that is significantly less than buckling forces (*e*.*g*., ≤10%). Thus, we obtain a design requirement on *t*_max_/*l* (see SI for details).

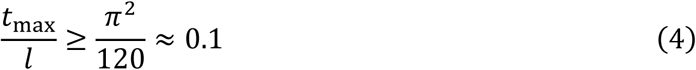

This criterion, shown as the blue shade in Figure 3e, suggests that an edge must have a thickness larger than 10% of its length to avoid any significant curvature.

It is also important to examine the flexibility in multi-edge joints. Auxetic designs assume that a joint at a vertex provides flexibility, allowing the structure to deform in response to external loads. Experimentally, the flexibility is realized by adding a ssDNA segment, and the longer segment will have more flexibility. Such a soft joint, however, creates a distance between edges at a joint, resulting in structural uncertainties. Therefore, a balance between structural accuracy and flexibility must be reached in DNA origami. To quantify the flexibility and integrity of a joint, we define the stretch *ξ* as the ratio of the displacement of ssDNA to the distance. For example, 0% stretch implies that the joint is in a complete relaxation (*i*.*e*., loose joint), while 100% indicates a full stretch (*i*.*e*., not flexible).

In theory, the structural accuracy is better maintained with a shorter ssDNA segment. We hypothesize that the displacement must be less than the minimum thickness of the edge, for example, Δ*d* ≤ 0.8×*t*_min_. This criterion yields *ξ* ≥ 55%. In parallel, the joint shall have enough flexibility such that as neighboring edges are pulled around the joint upon loading, the curvature of the edges can be avoided. To satisfy this condition, we propose that *δ*_*f*_ /*l* must be less than 10%, yielding another criterion of *ξ* ≤ 70% (Figures S21). Combining the two criteria, we obtain a guiding principle for multi-edge joints as indicated as blue shade in Figure 3f.

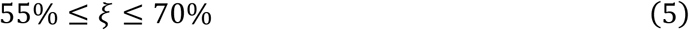

Equations 4 and 5 provide recommended designs to ensure appropriate edge rigidity and joint stretch/flexibility. To ensure rigid edges without significant flexure, the thickness must meet the requirement; as for the stretch of ssDNA at the vertex, the joint should be in a good balance between structural accuracy and flexibility. The re-entrant triangle deviates the edge criterion at *γ* ≤ 27° as shown in Figure 3e. The MD simulations also suggest that the stretch of ssDNA joints is approximately 45% (Figure 3f). Thus, the structure appears to have curved edges with blunt joints.

### Stiffness Reinforced Rotating Square

We examined the validity of the design strategies with another structure, rotating square (Figure 4a). This structure, unlike the re-entrant honeycomb and triangle, is an isotropic, single unit design (Figure 4b). In theory, 2D isotropic materials should have a range of the Poisson’s ratio from −1 to 1, whereas anisotropic structures can have any values of *ν*.^33^ The rotating square in particular is centrosymmetric with *ν* = −1. In comparison, the Poisson’s ratio of the re-entrant triangle is at approximately −2 at *γ* = 30°.

**Figure 4.**
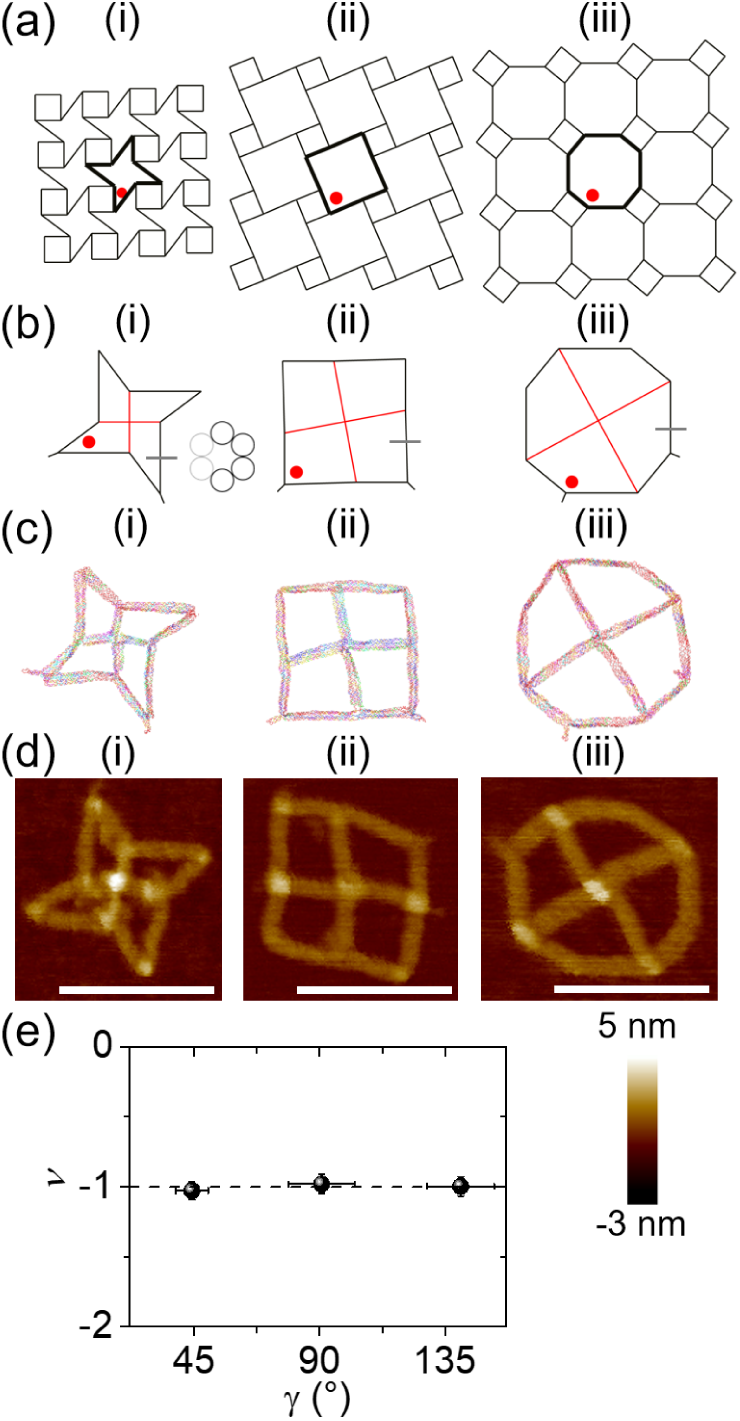
Stiffness reinforced rotating square. (a)–(b) Schematics and designs of the rotating square under different deformations. The angle *γ* = 40° (i), 90° (ii), and 140° (iii). Regular and jack edges are shown in black and red. The inset shows the edge cross-section, where four double helices (black) are used in a hexagonal arrangement. (c) MD simulations and (d) AFM images of the rotating square DNA origami. Scale bar: 100 nm. (e) Poisson’s ratio versus angle. Solid circles show the experimental data with error bars, while the dashed line represents the ideal centrosymmetric rotation with *ν* = −1 regardless of *γ*.

In our origami design, each edge is stiffened with four dsDNA helices (such that *t*_max_*/l* = 0.1) as shown in the inset of Figure 4b (Figures S6, S16), and shorter ssDNA segments are used for the joint (Table S3.3), aiming for ∼60% stretch. Jack edges are also used, shown in red in Figure 4b. Figure 4c–d shows the MD simulation and experimental results at various angles. It is evident that each edge is nearly straight, there are no blunt joints, and the overall structures in equilibrium are well formed. The measured Poisson’s ratios are approximately −1 (Figure 4e).

We performed mechanical deformations using oxDNA with jack edges removed (Figure 5a). We also applied the finite model on the rotating square to further validate the model and design guidelines (see SI for details). As the origami compresses from *γ* = ∼137° to ∼39° by pulling the structure toward the center (Figure 5b), *F*_acc_ increases monotonically as expected, which is in excellent agreement with the theoretical prediction (Figure 5c). Approximately 35 pN is needed to deform the origami fully. Young’s modulus has a strong angular dependence, as shown by the model and simulation (Figure 5d). Similar to the re-entrant triangle, this is largely due to the change of dominant deformation modes. In the rotating square, the stretching mode is predominant at both large and small angles. In the mid-range of angles, however, flexing and hinging become more important. The structure will be less rigid at around 80° given *K*_*f*_ << *K*_*s*_, thus showing a minimum of *E* = ∼0.43 pN/nm^2^. During the simulated deformation, the edge flexure is maintained below the threshold, which reasonably follows the model prediction (Figure 5e). This suggests that the edges are stiff enough to resist flexing deformation. Further, the joint stretch is also within the preferred range throughout the deformation, implying every joint is in modest tension (Figure 5f). The agreement between design, theory, simulation, and experiment confirms the applicability of the finite model and recommended design guidelines. Compared with the re-entrant honeycomb and triangle, the rotating square demonstrates significantly improved structural accuracy. The model and design guidelines should be applicable for various edge cross-sections and joint designs.

**Figure 5.**
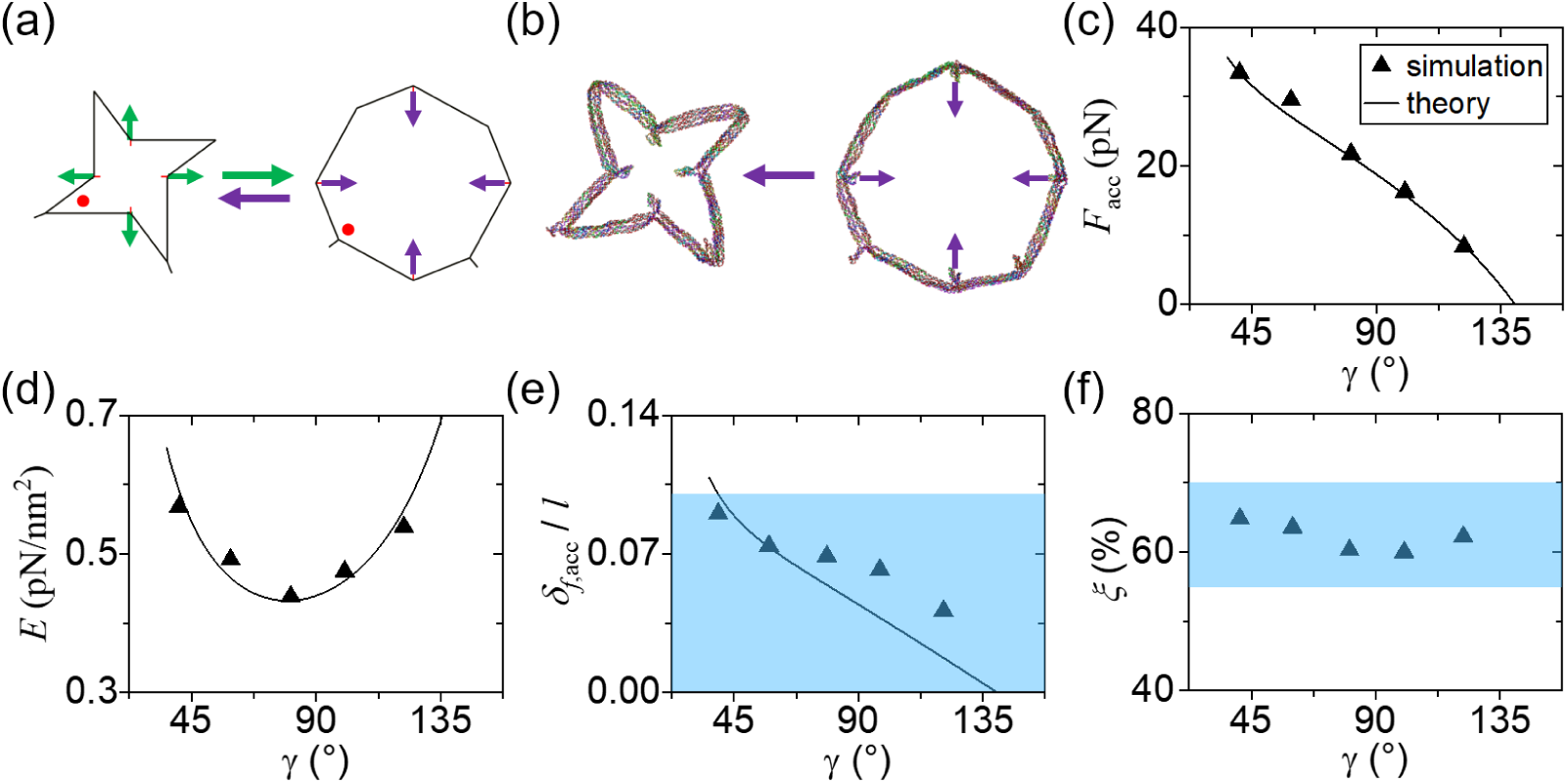
Mechanical deformation of the rotating square in MD simulations. (a) Schematics of the computational domain with the jack edges removed. (b) Simulation results of the re-entrant triangle under compression from *γ* = ∼137° to ∼39°. (c) The accumulated external force (*F*_acc_) monotonically increases as the angle *γ* decreases, reaching ∼35 pN at *γ* = 39°. (d) Young’s modulus has a strong angular dependence, with a minimum value of *E* = ∼0.43 pN/nm^2^. The simulation data and the finite model prediction are in excellent agreement. (e)–(f) Dimensionless flexure (*δ*_*f*,acc_/*l*) and joint stretch (*ξ*) as a function of *γ*. Both edge flexure and joint stretch data are all within the preferred ranges.

## DISCUSSION

### Chemical vs. Mechanical Deformations

In this study, we explored chemical and mechanical loads to enable structural deformations. There are two major steps in chemical deformation: (i) jack-staple release process via toehold-mediated strand displacement and (ii) recombination process via reannealing. A set of staples for jack edges are removed by fully complementary releaser strands, during which no forces are induced to the structure. After the release process, a new set of staples are added for jack edges, where forces towards the target state are introduced. Therefore, the deformation forces may be estimated by considering the free energy change (Δ*G*) and the displacement during the recombination process (see SI for details). The average force per jack edge for reconfiguring the re-entrant triangle is estimated to be ∼55 pN. Since the free energy change is the maximum driving potential for structural transformation, this should be the upper limit of the force. In the computational studies, the displacements are caused directly by the applied forces. During the compression, the force is small at the beginning, and then it increases and reaches the final value of ∼21 pN. The calculated force is the minimum needed for mechanical deformation, thus it is smaller than the force associated with chemical deformation. In the rotating square, the chemical and mechanical forces are approximately 72 and 35 pN, which are the upper and lower limits, respectively. The different forces for the re-entrant triangle and rotating square are due to both the geometry and cross-section design (*e*.*g*., two-dsDNA vs. four-dsDNA edges). Larger forces are needed for deforming the rotating square with stiffer edges, tighter joints, and greater displacements.

### Unidirectional vs. Centrosymmetric Loading

The re-entrant and regular honeycombs, as well as the re-entrant triangle were loaded in one direction by modulating jack edges, and the orthogonal direction followed for overall reconfiguration. In contrast, the external forces were applied in two orthogonal directions given the centrosymmetric structure of the rotating square. Such loading creates additional moment of force (see SI for details). In the design, the edge lengths *l* > *m* causes a clockwise rigid-body rotation. The rotational motion leads to a contraction of the entire structure. For stress calculations, a projected area is normally defined as the area perpendicular to the loading direction. With two orthogonal directions, there is no perpendicular area for stress. We thus defined a hypothetical area that has an angle of 45° with both loading directions as the projected area. As a result, the Young’s modulus of the rotating square is less than that of the re-entrant triangle, even with the stiffness reinforced structure.

### Validity of Design Requirements

The design requirements in equations 4 and 5 are consistent with previous reports. Yan *et al*. explored the edge rigidity in wireframe DNA origami using single and two dsDNA helices.^18,21^ The *t*_max_/*l* used is ≥ 0.1, satisfying equation 4. Seeman and colleagues found that the more helices are used, the longer the persistence length a DNA edge has.^34^ They further noted that the increase of persistence length does not result from the addition of helices, but rather originate from the enhanced thickness associated with a given bend axis. This directional dependence is considered in our finite model and simplified using the maximum thickness in equation 4.

For joint flexibility, Saleh and coworkers studied the elasticity of ssDNA with different sequences and found that when the strands are in 60% or greater of their fully-stretched lengths, their mechanical behaviors are similar regardless of sequences.^35^ Under this condition, base-stacking, a sequence-dependent interaction, is no longer effective such that Young’s modulus of ssDNA becomes nearly uniform. This sequence independence is critical for the design process because it is difficult to specify the sequence of ssDNA joints for a given scaffold. Bathe and colleagues reported multiple autonomously designed wireframe DNA origami structures using 0.42 nm/base for ssDNA segments at the joints.^36^ Assuming the length of fully stretched ssDNA of ∼0.63 nm/base,^37^ the stretch used was approximately 67%, agreeing with equation 5.

We also examined the design requirements for edges and joints using the honeycomb structures. The stretch values are mostly lower than 55% in both regular and re-entrant honeycombs (Figure S23), indicating the joints are under-stretched (*i*.*e*., too loose). Their edges are slightly less rigid than the requirement (Table S4.2). As a result, the honeycombs show blunt joints and minor curved edges as seen in Figure 1f–g. In addition, we tested another control design of re-entrant triangle with weak edges and loose joints. Its designed *t*_max_/*l* is ∼5%, significantly lower than the recommendation (Figure S7, Table S4.2), and the joints are mostly under-stretched (Figure S24). As a result, this control structure does not reasonably follow the re-entrant triangle design and appears significantly worse than the original re-entrant triangle (Figure S40–41). Overall, the edges and joints must satisfy the respective design requirements, together ensuring the integrity of the periodic cellular structures.

### Infinite vs. Finite Models

This study explores both infinite and finite models using wireframe DNA origami for nanoscale auxetics. The infinite model describes ideal design strategies for auxetic metamaterials, while the finite model accounts for the mechanics of DNA. Regardless of the model used in the design, the requirement on edges in equation 4 should be met if possible. Under the infinite model, the overall size of the structure and the most suitable cross-section of edges are determined by the total length of the scaffold. Ideally, a joint is a flexible point connecting edges. However, a wireframe origami uses soft spacers made of ssDNA. The number of nucleotides at a joint may be calculated based on the stretch level (*e*.*g*. ∼60% or ∼0.38 nm/base) to ensure sufficient flexibility while maintaining structural integrity. In most cases, this is a quick and reasonable approach, especially when the structures are static. For dynamic structures, it may be difficult to find a joint design that satisfies the requirement in the whole range of deformation. For example, a ssDNA segment for an angle between 30° and 60° will be different from the spacer designed for the 60°∼150° range. Thus, detailed calculations based on the finite model should be performed to reach a good balance between structural accuracy and flexibility. It is important to note that the effects of structural mechanics are not as critical in macroscopic metamaterials because the edges are typically made of stiff materials (∼10^3^ pN/nm^2^) and the deformations are either often limited (≤10%)^38^ or accommodated using specially designed hinges for a wide range of reconfiguration.^39^

### Biomolecular Auxetics

The sizes of auxetic unit cells made of various materials via different manufacturing methods are summarized in Figure S1. There is a wide gap between molecular auxetics and macroscopic metamaterials. Both DNA and proteins may serve as self-assembling building blocks to fill the gap. With the ability to form rigid components and flexible parts, they could be ideal candidates for periodic auxetic architectures with well-defined unit cells. Proteins, when folded, can be rigid, while unfolded parts are more flexible. Tezcan and coworkers constructed micron-sized, isotropic 2D networks of protein assemblies based on disulfide bonds and metal-coordination.^40^ They demonstrated stimuli-responsive, centrosymmetric transformation of the protein crystals, exhibiting ν = ∼−1.

In comparison, DNA self-assembly uses base-pairing and strand displacement based on hydrogen bonding. With the chemical and structural simplicity, DNA origami may be more advantageous in achieving geometrical complexity and controllability. Complex architectured materials may be constructed through sequence design and arrangement. DNA structures are not limited to any specific shapes, and both isotropic and anisotropic geometries are possible as shown in this work. Auxetic 3D structures should be also achievable. The DNA origami in this work includes one or few unit cells, yet polymerization via sticky ends or increased ionic strength could lead to large-scale structures.^41,42^ Further, other strategies such as DNA bricks^43^ and 3D DNA crystals^44^ may be employed to create macroscopic mechanical metamaterials.

In conclusion, auxetic 2D nanostructures are realized using DNA origami. Their elastic properties rely largely on the designed cellular structures and may be estimated quickly and roughly through the infinite model. However, the finite rigidity of edges and the flexibility of joints will affect the structural behaviors, for which DNA properties must be considered in the design. Mechanics-based guiding principles are developed to prevent curved edges and loose joints. As a bottom-up approach, DNA origami opens the opportunities for constructing well-defined auxetic unit cells and, with proper scale-up strategies, macroscopic metamaterials in 2D and 3D.

## ACKNOWLEDGMENT

This work was financially supported by the U.S. Department of Energy. J.H.C. acknowledges partial support from the U.S. National Science Foundation.

