## Supplementary Materials for "Auxetic Two-Dimensional Nanostructures from DNA"

**Content**

S1. Lengthscale of Auxetic Unit Cells
S2. Materials and Methods
S3. DNA Origami Designs
S4. Theories and Model Systems
S5. MD Simulations with oxDNA
S6. Supporting AFM Images
S7. DNA Sequence Information
S8. References

### S1. Lengthscale of Auxetic Unit Cells

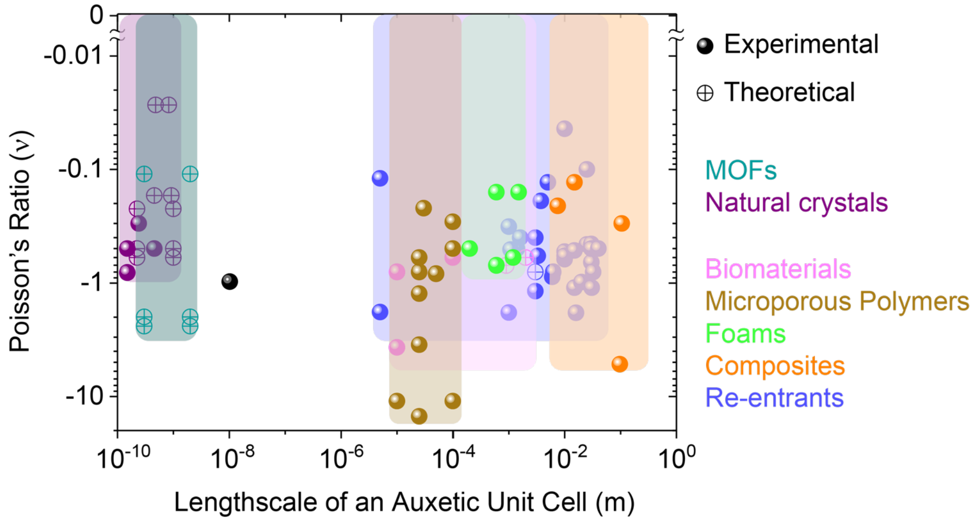


**Figure S1.** Poisson's ratio versus lengthscale of auxetic unit cells from the literature. There are two major groups in terms of unit size: (i) molecular auxetics with sub-nanometer unit cells and (ii) macroscopic metamaterials whose unit cells are microns or larger.

#### Molecular Auxetics

This category includes natural crystals and metal-organic frameworks (MOFs), whose unit cells are in the sub-nanometer range. The natural crystals exhibiting auxetic properties include α-cristobalite[1,2], cubic metals[3], face-centered cubic (FCC) crystals[4], graphene[5], black phosphorus[6,7] and tinselenidene[8]. These crystals are non-designable and typically anisotropic, showing auxetic behaviors only in certain directions. In theory, MOFs, compounds with metal ions and organic groups, can exhibit negative Poisson's ratios by arranging the chemical bonds in auxetic designs.[9-11] However, they have not been demonstrated experimentally.

#### Macroscopic Metamaterials

This group includes biomaterials, microporous polymers, foams, composites, and re-entrants. The major criteria on classifying these materials are the lengthscale and the structure of the unit cells. Biomaterials are different from others in that they are based on the origin of materials. They include bones[12], teat tissues[13], and tendons[14], whose unit cells range from microns to millimeters. Their auxetic properties may come from the arrangement of biological cells and/or protein fibers such as collagen and elastin. Microporous polymers are made of various polymers including polyethylene[15-20], polypropylene[21], and polytetrafluoroethylene (PTFE)[22]. Their unit sizes are tens to hundreds of microns. It is worth noting that PTFE often exhibits a Poisson's ratio below -10 due to significantly anisotropic units. Similar to microporous polymers, foams often have re-entrant features that enable auxetic behaviors. However, their unit sizes are significantly larger, up to few millimeters, and are made of metals as well as polymers.[23-26] The composites are unique in that they are made of two or more materials, for example, a fiber and an elastomer.[27-30] Typically, the fiber is stiffer than the elastomer and is used to reinforce the elastomer for improved mechanical strength. By having different arrangements and orientations of fibers, the composites can possess distinct Young's modulus and Poisson's ratio. They are often highly anisotropic and auxetic in one or two directions. Since the fibers are in the range of millimeters, the unit cells of the composites typically range from centimeters to decimeters. Re-entrants are categorized based on the structure of the unit cells, which belongs to a class of well-studied, periodical structures, among which the re-entrant honeycombs are the most well-known. The main difference between the re-entrants and the re-entrant structures of microporous polymers and foams is manufacturing. For example, the re-entrants are constructed by precisely fabricating edges and joints using either additive manufacturing (*e.g.* direct laser writing and 3D printing) or subtractive methods (*e.g.* laser ablation). In contrast, the re-entrant structures of microporous polymers and foams are produced by modulating overall physical and chemical conditions, such as temperature, pressure, and chemical concentrations. Using advanced manufacturing technologies, re-entrants can be made from many kinds of materials, such as polymers, metals, and ceramics. Moreover, the unit cells can be as small as few microns and can go as large as few centimeters.[31-52]

Besides those listed above, a previous review[53] reports that sintered ceramics may have the unit size of 0.1 nm to 10 μm. However, their unit cells are not well defined, and the overall auxetic properties likely result from the porous structures in the ceramics.[54] Therefore, sintered ceramics are not included in Figure S1.

**The Gap at the Nanoscale**

Figure S1 includes a single black filled circle which indicates isotropic 2D networks of protein assemblies with unit cells less than ten nanometers and the Poisson's ratio of -1 (in a recent publication by Tezcan and colleagues).[55] Even with this report on auxetic protein assemblies, it is clear that there is a wide gap between few nanometers and microns. The gap may be filled by biomolecular self-assembly. As discussed in the main text, there are similarities and differences between protein networks and DNA origami. This work exploits DNA self-assembly, realizing nanoscale auxetics.

### S2. Materials and Methods

#### Materials

All DNA oligomers, including staples and releasers, were obtained from Integrated DNA Technologies (sequence information presented in S7). The M13mp18 scaffold was supplied by Bayou Biolabs. All DNA strands were used directly from plates or tubes without further purification. All other chemicals were purchased from Sigma Aldrich.

#### DNA origami synthesis

All the origami structures were synthesized by mixing 10 nM scaffold strands with 4× DNA staple strands in 1× TAE buffer (an aqueous solution of 40 mM trisaminomethane, 1 mM ethylenediaminetetraacetic acid (EDTA) disodium salt, and 20 mM acetic acid at pH ~8) that also contains 12.5 mM magnesium acetate (termed TAEM buffer). The mixture was then thermally annealed from 90 °C to 4 °C in a BIO-RAD S1000 Thermal Cycler. The annealing process was adopted from Yan *et al.*'s work.[56] First, the mixture was annealed from 90 °C to 85 °C at -4 °C per 5 min; then from 85 °C to 70 °C at -1 °C per 5 min; after that, from 70 °C to 40 °C at -1 °C per 15 min; later on, from 40 °C to 25 °C at -1 °C per 10 min; finally, it was held at 4 °C. Before AFM imaging, no further purification was performed.

#### Chemical deformation of DNA origami

There were two major steps in chemical deformation, strand displacement and reannealing.[57,58] The target structures from synthesis were mixed with 30× DNA releasers. The mixture was then incubated at 44 °C for 12 hours for toehold-mediated strand displacement. After that, purifications were performed 3 times by using the centrifugal filter (100 kDa) from Amicon. Then, 20× new DNA staples designed for a different length of jack edges were mixed with the purified structures and incubated at 40 °C for 18 hours. Then, the temperature was reduced from 40 °C to 20 °C at a rate of 1 °C per 1 min. After reannealing, the mixture was stored at 4 °C. The incubation and reannealing were performed with a BIO-RAD S1000 Thermal Cycler. No further purification was performed before AFM imaging.

#### AFM imaging

Before AFM imaging, all DNA origami samples, buffers and mica surfaces were kept at 4 °C to keep the samples fresh and suppress thermal fluctuation. For deposition, the target sample was diluted to 0.5 to 1.0 nM with a buffer (termed 1× MES buffer), which contains 50 mM 2-(N-morpholino)ethanesulfonic acid (MES) sodium salt, 5 mM magnesium chloride, and 50 mM NaCl (pH ~6.5). An aliquot of 10 µL diluted sample was pipetted onto mica surface and incubated for 5 min at 4 °C. Then, 20 µL 1× MES, also containing 3 mM nickel chloride, was added onto the mica to increase the adhesion of DNA to mica. After 2 min of second incubation at 4 °C, the mica was blown dry with compressed air and rinsed with 80 µL deionized (DI) water for about 3 sec and then blown dry again with compressed air.

AFM imaging was performed in air in the Peak-Force tapping mode with a Bruker Dimension Icon AFM and SCANASYST-AIR probes. For each mica surface, at least three different places were imaged. On each spot, large-scale and zoom-in scans were performed. For large-scale scans, either 2 × 2 µm or 1.5 × 1.5 µm areas were characterized. The zoom-in scans measured 500 × 500 nm.

### S3. DNA Origami Designs

#### Overall design strategy

The focus of origami design is the construction of periodic cellular structures, which consist of edges and joints. Each edge is made of 2 or 4-dsDNA bundles depending on the size and shape of the origami. Each dsDNA bundle is composed of a scaffold segment and staples, forming crossovers to bring them together. A joint consists of unpaired scaffold segments and/or ssDNA domains (poly Thymidine or poly T)[56]. The edges signify the overall shape, while the joints link the edges together and combine all the scaffold segments into a loop. In each design, one circular (closed loop) scaffold is used (M13mp18).

There are two strategies used in this work: *routing* and *lay-out* methods. The routing method focuses on the joints. Since linking closest neighboring scaffold segments may lead to excessive loops (11 loops as shown in Figure S2c), some links are arranged to connect the opposite or diagonal neighboring segments (Figure S2d). By carefully arranging the linkages, one closed scaffold loop is formed. This method may be better suited for designs with a limited number of joints. The lay-out approach requires *bridges* in certain edges, which are pairs of crossovers of the scaffold. Typically, one bridge will merge two loops of scaffold together. In such a way, joints link closest neighboring scaffold segments together, while loops created by joints are brought together by bridges until a single loop is reached (Figure S3d).[56] This method may work well for designs with a limited number of edges.

#### Re-entrant honeycomb

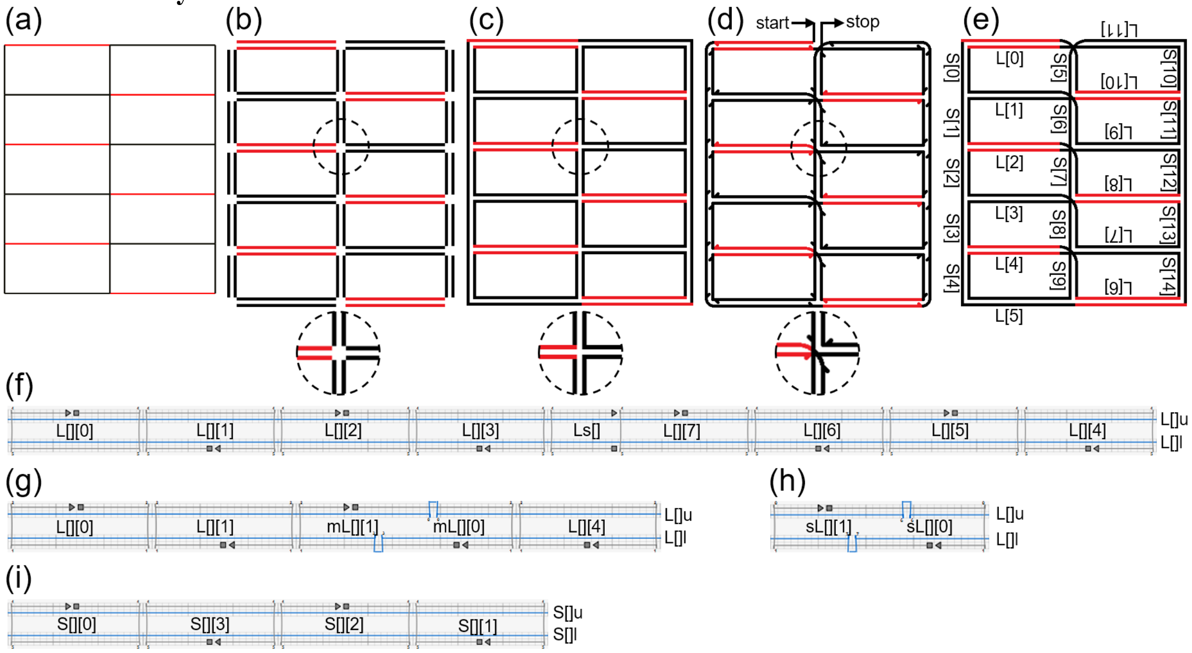


**Figure S2.** Schematics of the re-entrant honeycomb design. (a) Re-entrant honeycomb at $\gamma = 90°$. The black lines represent regular edges, while jack edges are shown in red. (b) Each edge is represented by two lines, which are two antiparallel segments of the scaffold. The inset shows the center joint when segments of scaffold are not linked yet. (c) The arrangement of scaffold if neighboring segments of the scaffold are connected. In such case, there are 11 closed loops. The inset shows the detailed center joint where segments are linked with the closest neighbor. (d) The arrangement of scaffold after applying routing method. The arrows denote the direction of the scaffold (from 5' to 3' end). The inset shows the detailed center joint where segments are linked with the closest, opposite, or diagonal neighbor. (e) The final routing of the scaffold with the start and stop points linked together, forming one closed loop. All the edges are named. The horizontal edges, including jack edges, are Ls (from 0 to 11), and the vertical edges are Ss (from 0 to 14). The orientations of the texts indicate the directions of the edges. (f)–(i) The detailed edge designs in caDNAno2.[59] Blue represents the scaffold segments and gray for staples. The squares and triangles mark the 5' and 3' ends of the staples, respectively. The staples are named with empty square brackets '[]'. The numbers to be filled in are the numbers of the corresponding L or S. For example, the left end staple of edge L is L[][0]. For the edge L[0], this staple is called L[0][0]. (f) The arrangement of scaffold segments and staples in both regular Ls and jack Ls for the conformation of $\gamma$ = 90°. The upper and lower scaffold segments are called L[]u and L[]l, respectively, which are 179 nt each. The staples with the length of 42 nt are named from L[][0] to L[][7], whereas the one with 22 nt, Ls[]. (g) Jack Ls for the conformation of $\gamma$ = 60°. Since the length of jack Ls at $\gamma$ = 60° is in between those at $\gamma$ = 90° and $\gamma$ = 29°, it is denoted as medium jacks. The associated staples are thus named mL[][0] and mL[][1] which are 32 nt and marked at the 5' ends. The scaffold segments are the same as for $\gamma$ = 90°. Since all the regular edges are kept the same, to present a different conformation of the whole structure, the base-paired region of jack edges has to be altered, whose length is 95 nt for each scaffold segment at $\gamma$ = 60°. Thus, there is a free loop of 84 nt marked as a bump in blue for each segment. (h) Jack Ls for the conformation of $\gamma$ = 29°. The scaffold segments are the same as for 90° and the base-paired region is 32 nt each, leaving a free loop of 147 nt each. The Ls at $\gamma$ = 29° are denoted as short jacks due to their shortest length among all three conformations. The associated staples are thus named sL[][0] and sL[][1], which are 32 nt each. (i) The arrangements of vertical edges S. The upper and lower scaffold segments are called S[]u and S[]l, which are 84 nt each. The staples, S[][0] to S[][3], are 42 nt each.

The re-entrant honeycomb is first designed for the conformation of $\gamma$ = 90°. Then, the jack edges were varied for other conformations. As shown in Figure S2, horizontal edges are named L and vertical edges are named S with numbers to distinguish them. Half of the Ls are jack edges (red) and the other half of Ls, together with all Ss, are regular edges (black). The designed lengths are calculated assuming 0.332 nm/bp for B-DNA. At $\gamma$ = 90°, all Ls are designed to be approximately 59 nm, all Ss, ~28 nm; at $\gamma$ = 60°, the jack Ls, ~32 nm; at $\gamma$ = 29°, and the jack Ls, ~11 nm.

Besides the jack edges, the joints between the edges are also important. The joints must be flexible enough to accommodate different angles, while they should not be too loose to link the edges together. To simplify the design, all the joints are made of single-stranded scaffold segments, whose lengths are adjusted based on the interior angle (of all three conformations) of the joints. Due to the nature of the routing method, the opposite and diagonal neighboring scaffold segments are linked to each other, besides the closest segments. As the angle becomes smaller, two jointed segments may sperate further apart. Thus, the length of a joint is generally inversely proportional to the minimum possible interior angle. In the following chart of the length of the segments, the joints are referred by the ending edges (from 5' to 3' end) – for example, the joint from L[0]u to S[0]l are noted as S[0]l. Further, since the scaffold is a closed ssDNA loop, it will return to the origin after a full route, which means that there is a joint noted as L[0]u (*i.e.* the joint from S[5]u to L[0]u).

**Table S3.1.** Minimum angle and the length of a joint of the re-entrant honeycomb.

| name | minimum angle (°) | length (nt) | name | minimum angle (°) | length (nt) |
|---|---|---|---|---|---|
| L[0]u | 90 | 8 | S[3]u | 90 | 7 |
| S[0]l | 90 | 7 | L[3]l | 29 | 10 |
| S[1]l | 58 | 9 | L[8]u | 180 | 3 |
| S[2]l | 58 | 9 | S[13]l | 90 | 7 |
| S[3]l | 58 | 9 | L[7]l | 29 | 10 |
| S[4]l | 58 | 9 | S[8]u | 29 | 10 |
| L[5]l | 29 | 10 | L[3]u | 29 | 10 |
| L[6]u | 180 | 3 | S[2]u | 29 | 10 |

| | | | | | |
|---|---|---|---|---|---|
| S[14]u | 90 | 7 | L[2]l | 90 | 7 |
| S[13]u | 58 | 9 | L[9]u | 180 | 3 |
| S[12]u | 58 | 9 | S[12]l | 29 | 10 |
| S[11]u | 58 | 9 | L[8]l | 90 | 7 |
| S[10]u | 58 | 9 | S[7]u | 90 | 7 |
| L[11]l | 29 | 10 | L[2]u | 90 | 7 |
| S[5]l | 29 | 10 | S[1]u | 90 | 7 |
| S[6]l | 58 | 9 | L[1]l | 29 | 10 |
| S[7]l | 58 | 9 | L[10]u | 180 | 3 |
| S[8]l | 58 | 9 | S[11]l | 90 | 7 |
| S[9]l | 58 | 9 | L[9]l | 29 | 10 |
| L[5]u | 29 | 10 | S[6]u | 29 | 10 |
| S[4]u | 29 | 10 | L[1]u | 29 | 10 |
| L[4]l | 90 | 7 | S[0]u | 29 | 10 |
| L[7]u | 180 | 3 | L[0]l | 90 | 7 |
| S[14]l | 29 | 10 | L[11]u | 180 | 3 |
| L[6]l | 90 | 7 | S[10]l | 29 | 10 |
| S[9]u | 90 | 7 | L[10]l | 90 | 7 |
| L[4]u | 90 | 7 | S[5]u | 90 | 7 |

**Regular honeycomb**

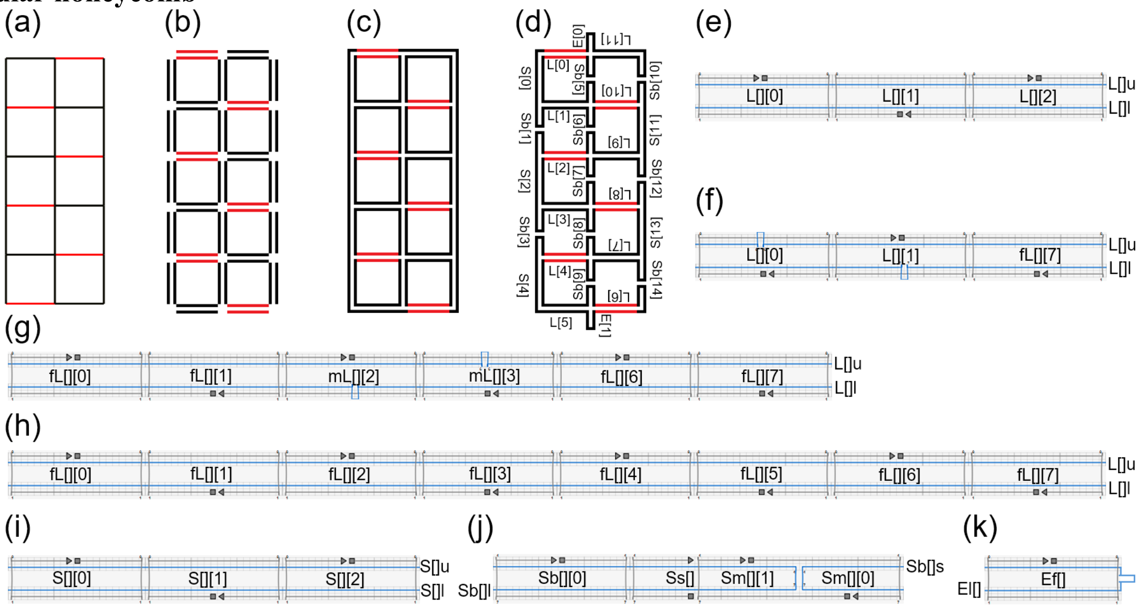


**Figure S3.** Schematics of the regular honeycomb. The color scheme and legends are the same as in Figure S2. (a) Honeycomb at $\gamma = 90°$. (b) Each edge is represented by two lines, which are two antiparallel segments of the scaffold. (c) The intermediate step during applying lay-out method. (d) The final lay-out of the scaffold with the bridges added in the structure and extra scaffold segments left at the top and bottom. All the jack edges, shown in red, are Ls. (e)–(k) The detailed edge designs in caDNAno2. (e) The arrangement of scaffold segments and staples for regular Ls. The upper and lower scaffold segments are called L[]u and L[]l, respectively, which are 63 nt each. The staples, L[][0] to L[][2], are 42 nt each. (f) Jack Ls for the conformation of $\gamma = 90°$. The scaffold segments and staples are similar to those in (e). In order to present different conformations while keeping all regular edges unchanged, the base-paired region of each segment is 63 nt, leaving a free loop of 105 nt each (marked as a bump in blue in each scaffold segment). The staples, L[][0], L[][1], and fL[][7], are 42 nt each. (g) Jack Ls for the conformation of $\gamma = 120°$. The scaffold segments are the same as for $\gamma = 90°$ and the base-paired region is 126 nt each, with a free loop of 42 nt each (shown as a bump in blue). All the staples (fL[][0], fL[][1], mL[][2], mL[][3], fL[][6], and fL[][7]) are 42 nt each. Note that fL[][7] is identical with fL[][7] in (f). (h) Jack Ls for the conformation of $\gamma = 146°$. The scaffold segments are the same as for $\gamma = 90°$ and are fully base-paired (168 nt). All the staples, fL[][0], fL[][1], fL[][2], fL[][3], fL[][4], fL[][5], fL[][6], and fL[][7], are 42 nt each. Note that fL[][0], fL[][1], fL[][6], and fL[][7] are identical with those in (g). (i) Ss without any bridge. The upper scaffold segments are called S[]u, and the lower, S[]l, which are 63 nt each. The staples, S[][0] to S[][2], are 42 nt each. (j) Ss with a bridge (Sbs). The left scaffold segments are called Sb[]l (94 nt), and the right, Sb[]s (32 nt), which are labelled at the 5' ends, respectively. The staples, Ss[], Sm[][0], Sm[][1], and Sb[][0], have different lengths, varying from 22 to 42 nt. (k) The arrangement of Es which are placed at the top and bottom of the honeycomb. There is only one long scaffold segment in each Es, named El[]. The base-paired region is 42 nt each. The size of the free loop (marked as a bump in blue) in El[0] is 1122 nt, and that in El[1] is 1121 nt.

The structure is designed for $\gamma = 90°$ with jack edges variable for other conformations. As shown in Figure S3, horizontal edges are named L and vertical edges are named S (or Sb, meaning S with a bridge) and E with numbers to distinguish them. To guarantee a single scaffold loop, bridges are placed in some of the S edges (*i.e.* Sb edges). The Es are added just for holding unused scaffold segments (top and bottom together 2243 nt).

In the design, half of the Ls are jack edges (red) and the rest are regular edges (black). The designed lengths are also calculated based on 0.332 nm/bp. At $\gamma$ = 90°, all Ls are designed to be ~21 nm, all Ss (including Sbs), ~21 nm, and all Es, ~7 nm; at $\gamma$ = 120°, the jack Ls, ~42 nm; at $\gamma$ = 146°, the jack Ls, ~56 nm. As for the joints, the balance between flexibility and structural accuracy needs to be reached. Due to the nature of the lay-out method, only the closest neighboring scaffold segments are linked to each other. As the angle becomes larger, two jointed segments can sperate further apart. Besides, the separation can be different even at the same angle depending on the actual conformation of the joint. In the case of outside corners, longer joints are required. Overall, the larger the angle is, the longer the joint typically is. In the chart of the length of the segments, the joints are referred by the ending edges, following the same manner as in the re-entrant honeycomb.

**Table S3.2.** Angle range and the length of a joint of the regular honeycomb.

| name | angle range (°) | length (nt) | name | angle range (°) | length (nt) |
|---|---|---|---|---|---|
| L[0]u | 34~90 | 2 | L[5]u | 90~146 | 6 |
| S[0]l | 34~90 | 10 | S[4]u | 90~146 | 6 |
| Sb[1]l | 68~180 | 4 | L[4]l | 34~90 | 2 |
| L[1]l | 90~146 | 6 | Sb[9]s | 34~90 | 2 |
| Sb[6]s | 90~146 | 6 | L[7]u | 90~146 | 6 |
| L[10]u | 34~90 | 2 | Sb[14]l | 90~146 | 6 |
| S[11]u | 34~90 | 2 | S[13]l | 68~180 | 4 |
| L[9]l | 90~146 | 6 | Sb[12]s | 68~180 | 7 |
| Sb[6]l | 90~146 | 6 | L[8]l | 34~90 | 2 |
| L[2]u | 34~90 | 2 | Sb[7]l | 34~90 | 2 |
| Sb[1]s | 34~90 | 2 | L[3]u | 90~146 | 6 |
| S[2]l | 68~180 | 7 | S[2]u | 90~146 | 6 |
| Sb[3]l | 68~180 | 4 | L[2]l | 34~90 | 2 |
| L[3]l | 90~146 | 6 | Sb[7]s | 34~90 | 2 |
| Sb[8]s | 90~146 | 6 | L[9]u | 90~146 | 6 |
| L[8]u | 34~90 | 2 | Sb[12]l | 90~146 | 6 |
| S[13]u | 34~90 | 2 | S[11]l | 68~180 | 4 |
| L[7]l | 90~146 | 6 | Sb[10]l | 68~180 | 7 |
| Sb[8]l | 90~146 | 6 | L[10]l | 34~90 | 2 |
| L[4]u | 34~90 | 2 | Sb[5]s | 34~90 | 2 |
| Sb[3]s | 34~90 | 2 | L[1]u | 90~146 | 6 |
| S[4]l | 68~180 | 7 | S[0]u | 90~146 | 6 |
| L[5]l | 34~90 | 10 | L[0]l | 34~90 | 2 |
| El[1] | 90~146 | 6 | Sb[5]l | 34~90 | 2 |
| L[6]u | 34~90 | 2 | L[11]u | 90~146 | 6 |
| Sb[14]s | 34~90 | 10 | Sb[10]s | 90~146 | 6 |
| L[6]l | 34~90 | 2 | L[11]l | 34~90 | 10 |
| Sb[9]l | 34~90 | 2 | El[0] | 90~146 | 6 |

**Re-entrant triangle**

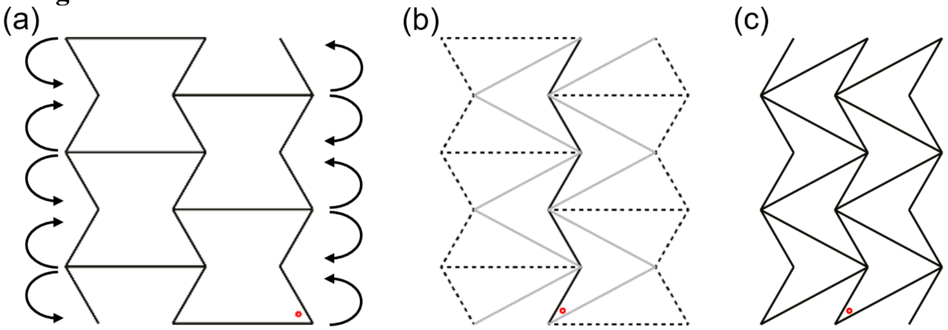


**Figure S4.** Schematics illustrating the evolution of a re-entrant triangle from a re-entrant honeycomb. The solid black lines represent regular edges. Jack edges are not shown. The internal angle ($\gamma$) is noted by a red dot. (a) The starting point: re-entrant honeycomb. Arrows indicate the directions of splitting and shrinking and merging. (b) The half-way: dashed lines depict the edges to be split and shrunk, while gray lines denote the ones to be merged into. (c) The result: re-entrant triangle. Overall, the re-entrant triangle may be seen as an evolved or reduced structure from a re-entrant honeycomb, where two short edges are shrunk to a point and one long edge is split into two, in one unit.

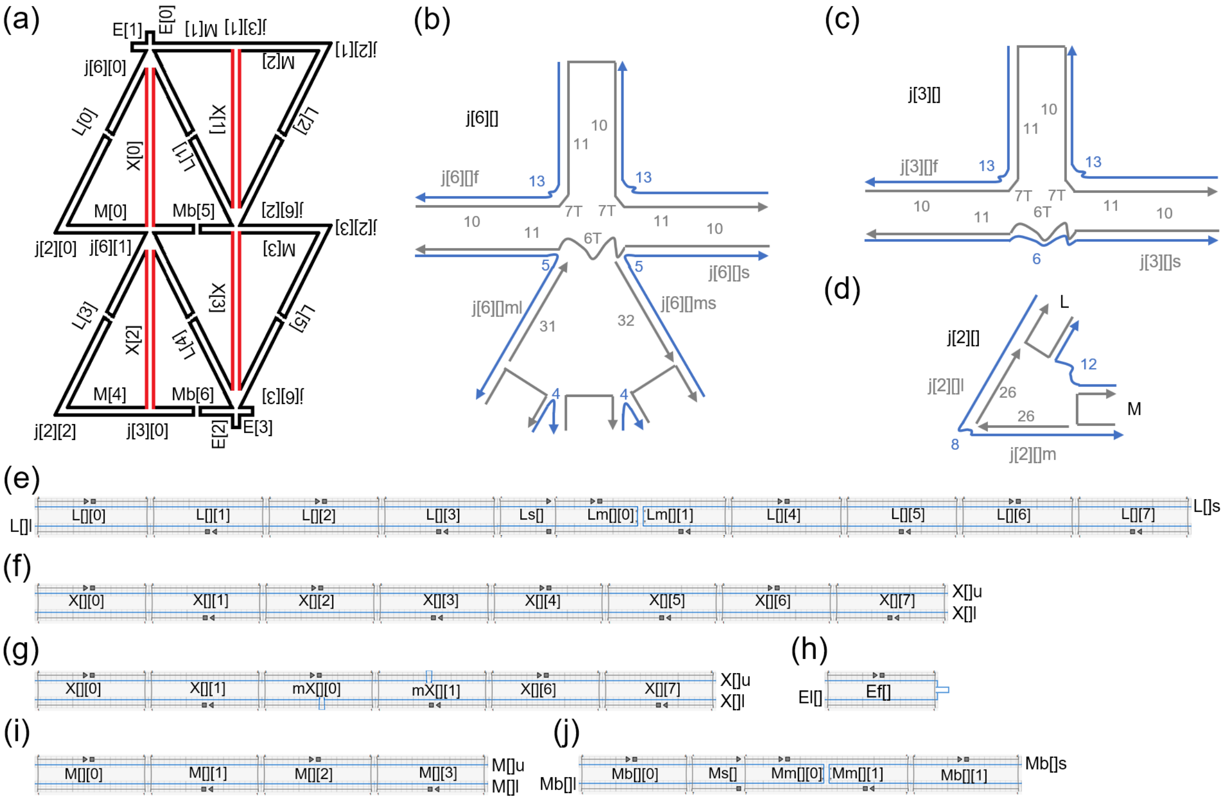


**Figure S5.** Schematics of the re-entrant triangle. The color scheme and legends are the same as in Figure S2. (a) Re-entrant triangle at $\gamma = 58°$ following the lay-out method. All the joints and edges are named. The names of the joints started with 'j', and the first numbers are the counts of edges involved. The horizontal edges are Ms (or Mbs, from 0 to 6) and two Es (total four, from 0 to 3), while the vertical edges are Xs (from 0 to 3) and the other two Es. The tilted edges are Ls (from 0 to 5), and the jack edges are Xs. (b)–(d) The detailed joint designs. The numbers to be filled in the square brackets are the numbers of the corresponding joints. Blue numbers indicate the free scaffold loops at the joints, and gray numbers with T denote the numbers of poly T for the staples. The gray numbers without T show the base-paired lengths of the staples. The names of the staples at the joints are in gray color. Their lengths range from 26 nt to 56 nt.

(e)–(j) The detailed edge designs in caDNAno2. The numbers to be filled in the square brackets are the numbers of the corresponding L, X, E or M. (e) The arrangement of Ls. The left scaffold segments are called L[]l (220 nt) and the right segments are named L[]s (200 nt), both of which are labelled at the 5' ends. The staples, L[][0] through L[][7], are 42 nt each, while Ls[] is 22 nt. Both Lm[][0] and Lm[][1] are 32 nt. (f) Jack Xs for the conformation of $\gamma$ = 58°. The upper and lower scaffold segments are 168 nt each and called X[]u and X[]l, respectively. The staples, X[][0] through X[][7], are 42 nt each. (g) Jack Xs for the conformation of $\gamma$ = 30°. The scaffold segments are the same as for $\gamma$ = 58° and the base-paired region is 126 nt each, with a free loop of 42 nt each (shown as a bump in blue). Staples, mX[][0] and mX[][1], are marked at the 5' ends and 42 nt each. (h) The arrangement of Es at the upper-left and lower-right corners of the triangle. There is only one long scaffold segment in each Es, named El[]. The base-paired region is 42 nt each. The size of the free loop (marked as a blue bump) in El[0] is 220 nt, while those in El[1] through El[3] are 221 nt each. (i) Ms without any bridge. The upper and scaffold segments are called M[]u and M[]l, respectively. Both are 84 nt each. The staples, M[][0] through M[][3], are 42 nt each. (j) Ms with a bridge (Mbs). The left and right scaffold segments are named Mb[]l (94 nt) and Mb[]s (74 nt), respectively. They are labelled at the 5' ends. The staples, Ms[], Mm[][0], Mm[][1], Mb[][0], and Mb[][1], have different lengths, varying from 22 to 42 nt.

The design is completed for $\gamma$ = 58° with jack edges variable for $\gamma$ = 30°. For some of the staples in jack edges, toeholds of 8 nt are placed at the 3' end so that the reconfiguration can be realized. As shown in Figure S5, edges are named L, X, E and M (or Mb, meaning M with bridge) with numbers to distinguish them. Only Xs are used for jack edges (red) and the rest are regular edges (black). The designed lengths (including the double stranded region of the corresponding joints) are also calculated based on 0.332 nm/bp. At $\gamma$ = 58°, all Ls are designed to be ~79 nm, all Xs, ~59 nm, all Es, ~7 nm, and Ms (including Mbs), ~36 nm; at $\gamma$ = 30°, all Xs, ~45nm.

### Rotating square

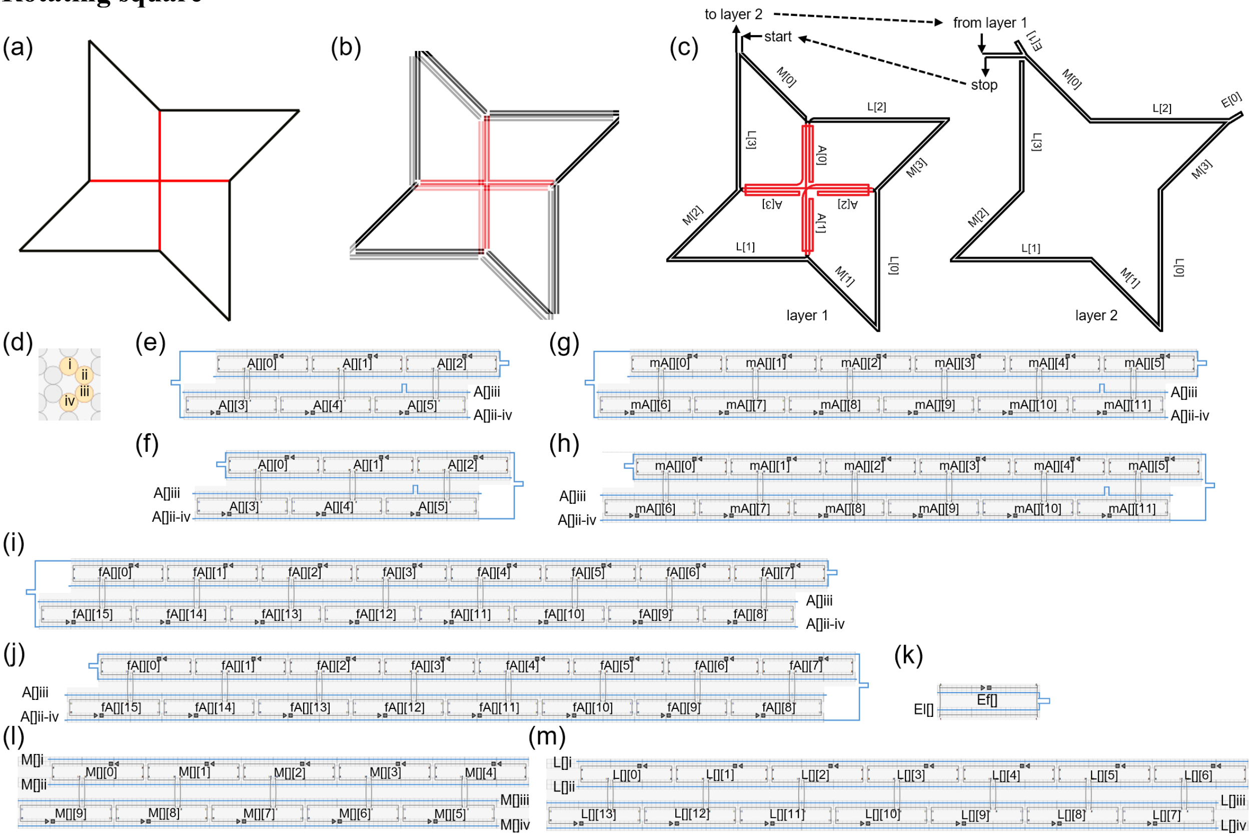


**Figure S6.** Schematics of the rotating square. The color scheme and legends are the same as in Figure S2. (a) Rotating square at $\gamma = 37°$. (b) Each edge is represented by four lines, indicating four scaffold segments. Each segment is antiparallel with the neighboring segment(s). (c) The final routing of the scaffold in two layers (which are for better illustration in the schematics) with the start and stop linked together and extra scaffold segments at the top-left and upper-right corners in layer 2. At the center, the jack edges are As (from 0 to 3), shown in red. The edges on the outside loop are Ms (from 0 to 3), Ls (from 0 to 3), and Es (from 0 to 1). (d)–(m) The detailed edge designs in caDNAno2. The numbers to be filled in the square brackets are the number of the corresponding A, M, L or E. Every staple is 42 nt. (d) The cross section of edge A, M and L with the segments named from i to iv. (e) Jack A[0] and A[2] for the conformation of $\gamma =$ 37°. The scaffold segments are called A[]iii (168 nt) and A[]ii-iv (524 nt), respectively. The base-paired region in A[]iii is 63 nt, while that in A[]ii-iv is 189 nt. The unpaired free loops are shown as bumps in blue. The free loop in A[]iii is 105 nt. The free loop on the left side in A[]ii-iv is 325 nt, while that on the right is 10 nt. The staples are named, from A[][0] to A[][5]. (f) Jack A[1] and A[3] for the conformation of $\gamma =$ 37°. The scaffold segments, A[]iii and A[]ii-iv, have the same length and base-paired region as in (e). The unpaired free loop in A[]iii is the same as in (e) as well. The free loop on the left side is 10 nt, while that on the right is 325 nt. The staples are named in the same ways as in (e). (g) Jack A[0] and A[2] for the conformation of $\gamma = 88°$. The scaffold segments are the same as in (e). The base-paired region in A[]iii is 126 nt, while that in A[]ii-iv is 378 nt. The free loop in A[]iii is 42 nt. The unpaired loop on the left side in A[]ii-iv is 136 nt, while that on the right is 10 nt. Since the jack Ls at $\gamma = 88°$ are recognized as medium jacks due to their length (similar to Figure S2g), the associated staples are named from mA[][0] to mA[][11]. (h) Jack A[1] and A[3] for the conformation of $\gamma = 88°$. Similar to (e) and (f), the differences between (g) and (h) are the free loops in A[]ii-iv. The loop on the left side in A[]ii-iv is 10 nt, while that on the right is 136 nt. (i) Jack A[0] and A[2] for the conformation of $\gamma = 140°$. The scaffold segments are the same as in (e) and (g). A[]iii is fully base-paired (168 nt), while the two free loops in A[]ii-iv are both 10 nt. With the fully base-paired A[]iii (*i.e.*, full length with no free loops), the staples are named from fA[][0] to fA[][15].

(j) Jack A[1] and A[3] for the conformation of $\gamma = 140°$. The free loops in A[]ii-iv are both 10 nt. Similar to (i), A[]iii is fully base-paired (168 nt), and staples are named fA[][0] through fA[][15]. (k) The arrangement of Es. There is only one scaffold segment in each E, named El[], and each base-paired region is 42 nt. The size of the free loop in El[0] is 39 nt, while that in El[1] is 40 nt. (i) The arrangement of Ms. The segments are 105 nt each and named M[]i to M[]iv. The staples are named from M[][0] to M[][9]. (j) The arrangement of Ls. The segments are 147 nt each and named L[]i to L[]iv. The staples are named from L[][0] to L[][13].

In this design, 4 dsDNA helices are used to mechanically strengthen the structure. The design is completed for $\gamma = 37°$ with jack edges variable for other conformations. As shown in Figure S6, the edges are named A, E, M and L with numbers to distinguish them. Only As are used for jack edges (shown in red) and they have special routing. For example, instead of going directly through, segment A[]ii-iv goes back and forth, serving as 'three scaffold segments' with two free-loop linkages (see Figure S6e–j). These linkages are used to modulate the jack length.

Except for As, all Es, Ms and Ls are regular edges (shown in black). The designed lengths are also calculated based on 0.332 nm/bp. At $\gamma = 37°$, As are designed to be ~21 nm, Es, ~7 nm, Ms, ~35 nm, and Ls, ~49 nm; at $\gamma = 88°$, As, ~42nm, at $\gamma = 140°$, As, ~56 nm. The joints are similar to those in the re-entrant honeycomb (Figure S2). The smaller the angle is, the longer the joint is. In the chart below, the joints are referred by the ending edges, in the same manner for the re-entrant honeycomb (Table S3.1).

**Table S3.3.** Minimum angle and the length of a joint of the rotating square.

| name | minimum angle (°) | length (nt) | name | minimum angle (°) | length (nt) |
|---|---|---|---|---|---|
| M[0]iii | 0 | 10 | L[1]ii | 127 | 5 |
| A[0]iii | 68 | 7 | M[2]iii | 37 | 9 |
| A[1]iii | 180 | 3 | L[3]ii | 127 | 5 |
| M[1]iii | 68 | 7 | El[1] | 162 | 5 |
| L[0]ii | 37 | 9 | M[0]i | 162 | 5 |
| A[2]iii | 61 | 8 | L[2]i | 127 | 5 |
| A[3]iii | 180 | 3 | El[0] | 162 | 5 |
| M[2]ii | 68 | 7 | M[3]iv | 162 | 5 |
| L[1]iii | 37 | 9 | L[0]i | 127 | 5 |
| A[1]ii-iv | 61 | 8 | M[1]iv | 37 | 9 |
| A[2]ii-iv | 90 | 6 | L[1]iv | 127 | 5 |
| M[3]iii | 68 | 7 | M[2]i | 37 | 9 |
| L[2]ii | 37 | 9 | L[3]iv | 127 | 5 |
| M[0]ii | 127 | 5 | L[3]i | 0 | 10 |
| L[3]iii | 37 | 9 | M[2]iv | 127 | 5 |
| A[3]ii-iv | 61 | 8 | L[1]i | 37 | 9 |
| A[0]ii-iv | 90 | 6 | M[1]i | 127 | 5 |
| L[2]iii | 61 | 8 | L[0]iv | 37 | 9 |
| M[3]ii | 37 | 9 | M[3]i | 127 | 5 |
| L[0]iii | 127 | 5 | L[2]iv | 37 | 9 |

| | | | | | |
|---|---|---|---|---|---|
| M[1]ii | 37 | 9 | M[0]iv | 127 | 5 |

**Re-entrant triangle, control**

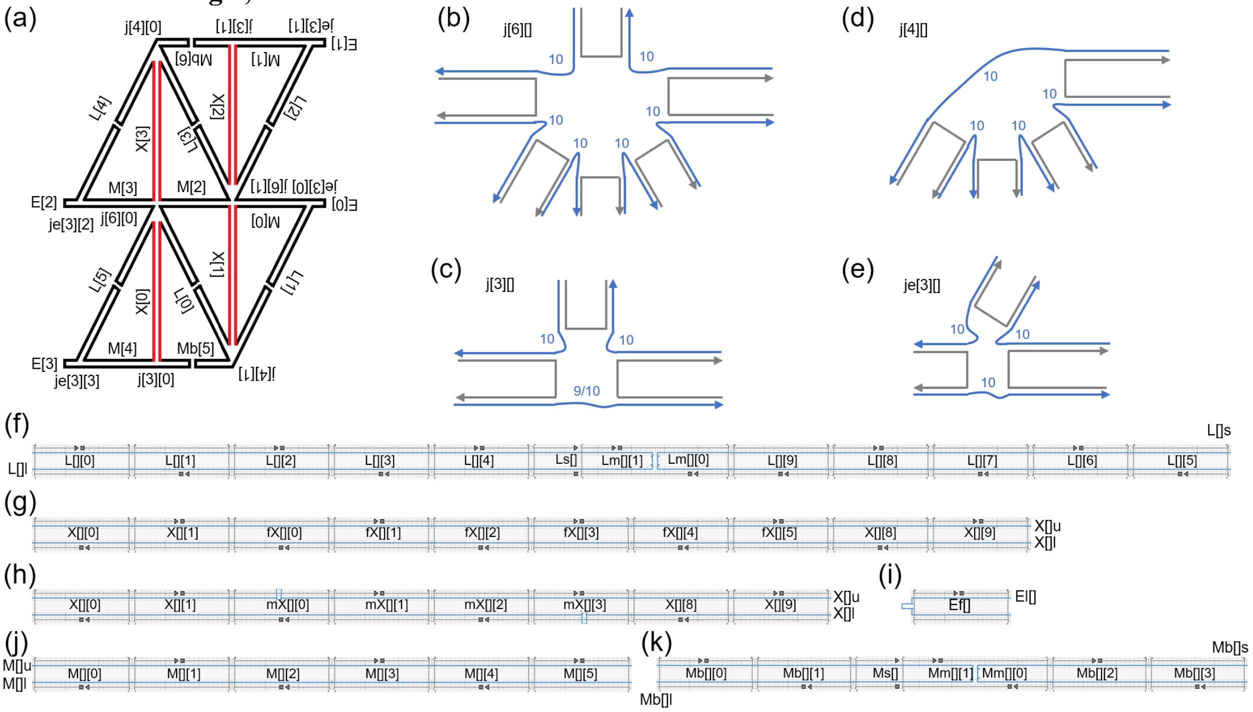

**Figure S7.** Schematics of the design of the re-entrant triangle control. The color scheme and legends are the same as in Figure S2. (a) Re-entrant triangle at $\gamma = 56°$ with the lay-out method applied. All the edges and joints are named. The horizontal edges are Ms (or Mbs, from 0 to 6) and Es (total from 0 to 3). The tilted edges are Ls from 0 to 5. The vertical edges are Xs from 0 to 3, which are jack edges. The names of the joints start with 'j', and the first numbers are the counts of edges involved. (b)–(e) The detailed joint designs. The numbers to be filled in the second square brackets are the numbers of the corresponding joints. Blue numbers denote the length (in nucleotides) of the free, unpaired scaffold segments. Note that '9/10' in (c) indicates that the length of the free scaffold is 9 nt in j[3][0], the linkage between M[4]l and Mb[5]l; it is 10 nt in j[3][1] between M[1]l and Mb[6]l. (f)–(k) The detailed edge designs in caDNAno2. The numbers to be filled in the square brackets are the numbers of the corresponding L, X, E or M. (f) The arrangement of Ls. The left and right scaffold segments are called L[]l (262 nt) and L[]s (242 nt), respectively. Both are labelled at the 5' ends. The 42 nt staples are named from L[][0] to L[][9], while Ls[] is 22 nt. Both Lm[][0] and Lm[][1] are 32 nt. (g) Jack Xs for the conformation of $\gamma = 56°$. The upper and lower scaffold segments are named X[]u and X[]l, respectively, which are 210 nt each. All the staples in Xs are 42 nt each. With jack Xs fully base-paired (*i.e.*, no free loop), the staples designed for $\gamma = 56°$ are named fX[][0] through fX[][5]. (h) Jack Xs for the conformation of $\gamma = 36°$. The scaffold segments are the same as for $\gamma = 56°$. The base-paired region is 168 nt each, with a free loop of 42 nt each. All the staples in Xs are 42 nt each. Staples X[][0], X[][1], X[][8], and X[][9] in (g) and (h) are identical. (i) The arrangement of Es at the left and right ends of the triangle. There is only one scaffold segment in each E, named El[], and the base-paired region is 42 nt. The free, unpaired loops of El[0] and El[2] are 58 nt each, while those of El[1] and El[3] are 59 nt each. (j) Ms with no bridge. The upper and lower scaffold segments are called M[]u and M[]l, respectively. They are 126 nt each. The staples, M[][0] to M[][5], are 42 nt each. (k) Ms with a bridge (Mbs). The left and right scaffold segments are named Mb[]l (136 nt) and Mb[]s (116 nt), respectively. They are all labelled at the 5' ends. The staples, Ms[], Mm[][0], Mm[][1], and Mb[][0] through Mb[][3], have different lengths, varying from 22 to 42 nt.

The major difference between this control and the original re-entrant triangle is the design of the joints. In the control, the design is simplified such that all the free scaffold loops are 10 nt except for one linkage between M[4]l and Mb[5]l, and there are no free staples present. As shown in Figure S7, the edges are named L, X, E, and M (or Mb, denoting M with a bridge) with numbers to distinguish them. Xs are for jack edges (shown in red). The designed lengths are also calculated based on 0.332 nm/bp. At $\gamma = 56°$, all Ls are designed to be ~87 nm, all Xs, ~70 nm, all Es, ~7 nm, and Ms (including Mbs), ~42 nm; at $\gamma = 36°$, all Xs, ~56 nm.

### S4. Theories and Model Systems

#### General analysis strategy

The theories are developed to predict the mechanical behavior of the structures. Among all the properties, Poisson's ratio and Young's modulus are the focus. To find the Poisson's ratio, (virtual) loading directions of corresponding structures are identified. Then, the displacements on loading directions and transverse directions are expressed as functions of the angles. With displacements, strains can be expressed, from which Poisson's ratios are determined. Below we show two distinct models, *infinite* and *finite*, which have different assumptions on edges and joints. Young's moduli are derived from the finite model. Stresses are calculated first by identifying force constants. With strains from displacements, Young's moduli are then obtained.

### S4.1 Infinite model

In this model, all the edges are assumed to be infinitely rigid, and all the joints are infinitely flexible. Thus, the edges will always be straight with their original lengths, and the joints will allow the edges to move freely about them.

#### Honeycombs

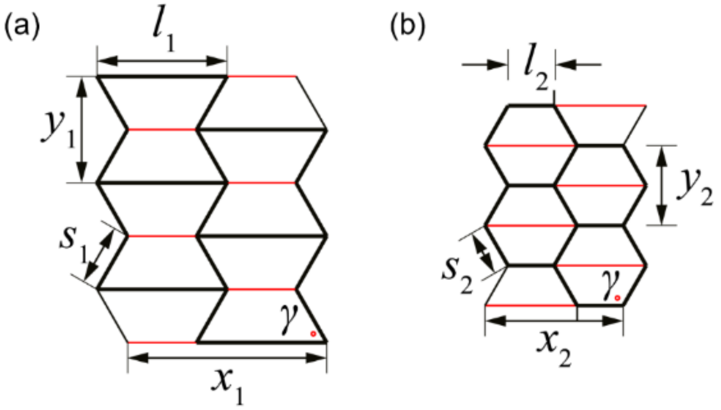


**Figure S8.** Honeycombs with all the lengths (*l*, *s*, *x*, *y*) and angles ($\gamma$) labelled in the figure. The *x* ($x_1$ or $x_2$) is the virtual loading direction. (a) Re-entrant honeycomb at 60°. (b) Regular honeycomb at 120°.

Since the two honeycombs share similar properties, the models are expressed together. For example, *l* and *m* mean either $l_1$, $m_1$ or $l_2$, $m_2$, considering the re-entrant or regular honeycomb, respectively. Similar notations are applied for *x* and *y*. From geometry,

$$x = 2l - 2s\cos\gamma \quad (4.1.1)$$

The displacement in *x* direction with respect to angle is

$$\mathrm{d}x = 2s\sin\gamma\,\mathrm{d}\gamma \quad (4.1.2)$$

The mechanical strain in *x* direction is

$$\varepsilon_1 = \frac{\mathrm{d}x}{x} = \frac{s\sin\gamma}{l - s\cos\gamma}\mathrm{d}\gamma \quad (4.1.3)$$

Similarly, $y = 2s\sin\gamma$, $\mathrm{d}y = 2s\cos\gamma\,\mathrm{d}\gamma$. The mechanical strain in *y* direction is then

$$\varepsilon_2 = \frac{\mathrm{d}y}{y} = \frac{\cos\gamma}{\sin\gamma}\mathrm{d}\gamma \quad (4.1.4)$$

Finally, Poisson's ratio is derived.

$$\nu = -\frac{\varepsilon_2}{\varepsilon_1} = \frac{(\cos\gamma - l/s)\cos\gamma}{\sin^2\gamma} \quad (4.1.5)$$

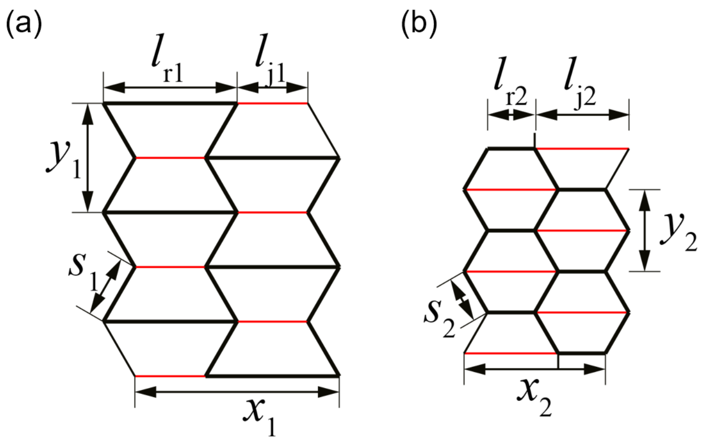

**Figure S9.** Illustration on the measurements of honeycombs. In addition to *s*, *x*, and *y* defined in Figure S8, $l_\mathrm{r}$ and $l_\mathrm{j}$ are added for the length of regular horizontal edges and jack edges. (a) Re-entrant honeycomb. (b) Regular honeycomb.

To calculate the Poisson's ratio experimentally, the edge lengths are measured every structure in AFM images and averaged. $l_\mathrm{r}$ and $l_\mathrm{j}$ mark the regular horizontal and jack edges, respectively. The expression for the Poisson's ratio is the same as equation (4.1.5) with $l$ replaced by $l_\mathrm{r}$. The angle is

$$\gamma = \cos^{-1}\left(\frac{l_\mathrm{r} - l_\mathrm{j}}{2s}\right) \tag{4.1.6}$$

**Re-entrant triangle**

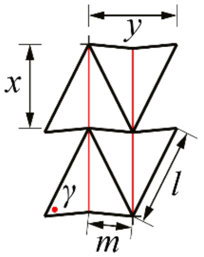


**Figure S10.** Re-entrant triangle at 58° with all the lengths ($l$, $m$, $x$, $y$) and angle ($\gamma$) labelled. The $x$ direction is the loading direction.

Similar to the analysis for honeycombs, by geometry,

$$x = \sqrt{l^2 + m^2 - 2lm\cos\gamma} \tag{4.1.7}$$

Then, the displacement and strain in $x$ direction are

$$\mathrm{d}x = \frac{lm\sin\gamma}{x}\mathrm{d}\gamma \tag{4.1.8}$$

$$\varepsilon_1 = \frac{\mathrm{d}x}{x} = \frac{lm\sin\gamma}{l^2 + m^2 - 2lm\cos\gamma}\,\mathrm{d}\gamma \tag{4.1.9}$$

Similarly, the length, displacement, and strain in $y$ direction are

$$y = 2lm\sin\gamma / x \tag{4.1.10}$$

$$\mathrm{d}y = \frac{2lm\cos\gamma}{x}\mathrm{d}\gamma - \frac{2lm\sin\gamma}{x^2}\frac{lm\sin\gamma}{x}\mathrm{d}\gamma \tag{4.1.11}$$

$$\varepsilon_2 = \frac{\mathrm{d}y}{y} = \frac{\cos\gamma}{\sin\gamma}\mathrm{d}\gamma - \frac{lm\sin\gamma}{x^2}\mathrm{d}\gamma \tag{4.1.12}$$

Finally, Poisson's ratio is determined.

$$\nu = -\frac{\varepsilon_2}{\varepsilon_1} = \frac{1 + (\cos\gamma - l/m - m/l)\cos\gamma}{\sin^2\gamma} \tag{4.1.13}$$

For measurement, the averaged $x$, $y$, $m$, and $l$ are determined in each conformation. The angle and the Poisson's ratio can then be calculated.

$$\gamma = \cos^{-1}\left(\frac{l^2 + m^2 - x^2}{2lm}\right) \tag{4.1.14}$$

$$\nu = -\frac{x}{y}\cdot\left(2\frac{\cos\gamma}{\sin\gamma} - \frac{y}{x}\right) \tag{4.1.15}$$

**Rotating square**

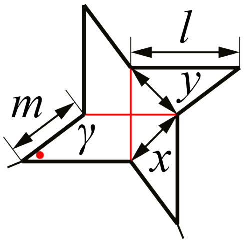


**Figure S11.** Rotating square at 37° with all the lengths ($m$, $l$, $x$, $y$) and angle ($\gamma$) labelled. Given the centrosymmetric structure and deformation, the directions of $x$ and $y$ may be defined in any perpendicular pair. Here, $x$ is defined as the distance between the end points of two neighboring jacks as shown in the schematic; $y$ is defined as the distance in the transverse direction that is perpendicular to $x$.

By centrosymmetric geometry,

$$x = y = \sqrt{m^2 + l^2 - 2ml\cos\gamma} \tag{4.1.16}$$

$$\mathrm{d}x = \mathrm{d}y \tag{4.1.17}$$

Thus,

$$\varepsilon_1 = \varepsilon_2 \tag{4.1.18}$$

$$\nu = -1 \tag{4.1.19}$$

The Poisson's ratio will thus remain the same regardless of the angle, which is valid for centrosymmetric structures.

For each conformation, the averaged $m$, $l$, $x$, and $y$ are obtained from the AFM images. The angle and the Poisson's ratio can then be calculated.

$$\gamma = \cos^{-1}\left(\frac{m^2 + l^2 - (x+y)^2/4}{2ml}\right) \tag{4.1.20}$$

$$\nu = -\frac{x}{y} \tag{4.1.21}$$

#### S4.2 Finite model

The infinite model discussed in S4.1 is based on ideal assumptions, that is, edges are assumed to have finite rigidity and joints are infinitely flexible. Below we develop the finite model that accounts for finite rigidity of the edges and finite flexibility of the joints.

##### S4.2.1 Force constants and displacements on edges of the re-entrant triangle

To describe the edges and joints, three distinct modes of force constants and related displacements are introduced. The force constants, similar to the stiffness in Hooke's law, relate the displacements to the forces which caused them.

$$K_{mode} = \left(\frac{F_{load}}{\delta_{load}}\right)_{mode} \tag{4.2.1}$$

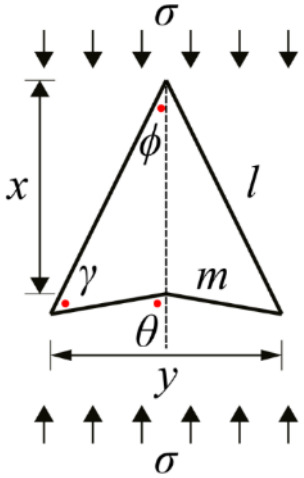


**Figure S12.** A single unit in the re-entrant triangle. The $x$ direction is the loading direction. All the lengths ($l$, $m$, $x$, $y$) labelled and angles ($\gamma$, $\theta$, $\phi$) are marked by red dots.

Three deformation modes are considered in the finite model: flexure, stretching, and hinging. Flexure and stretching describe the transverse and the axial deformation of the edges, respectively. Flexure force constant $K_f$ and stretching force constant $K_s$ are developed for an edge with length $l$ and $m$.[60,61]

$$K_f = \frac{12E_sI}{l^3} \tag{4.2.2}$$

$$K_s = \frac{E_sA}{l} \tag{4.2.3}$$

where $E_s$ is the Young's modulus of an edge, $A$ is the cross-section area of the edge, and $I$ is the second moment of the cross-section area. The hinging depicts the transverse deformation caused by angular deformation at the hinges. In the re-entrant triangle made of DNA, the hinging mode is enabled by ssDNA segments at the joints/hinges. When one of the ssDNA segments has flexure, the edges connected to it will have angular movements, and thus, hinging deformation.

$$K_{f,ssDNA} = \frac{F_{load}}{\delta_{load,ssDNA}} = 12\left(\frac{E_sI}{l^3}\right)_{ssDNA} \tag{4.2.4}$$

$$2\left(\frac{\delta_{load}}{l}\right)_{ssDNA} \approx \left(\frac{\delta_{load}}{l}\right)_{l} \tag{4.2.5}$$

Hence, hinging force constant $K_h$ may be estimated.

$$K_h = \frac{F_{load}}{\delta_{load,l}} = \frac{6}{l}\left(\frac{E_sI}{l^2}\right)_{ssDNA} \tag{4.2.6}$$

Similarly, the force constants may be expressed for the edge with length $m$.

$$K_{f,m} = \frac{12E_s I}{m^3} \tag{4.2.7}$$

$$K_{s,m} = \frac{E_s A}{m} \tag{4.2.8}$$

$$K_{h,m} = \frac{6}{m}\left(\frac{E_s I}{l^2}\right)_{ssDNA} \tag{4.2.9}$$

From equation (4.2.2), (4.2.3), and (4.2.6) through (4.2.9), the relationship between force constant of edge $l$ and edge $m$ is obtained.

$$K_{f,m} = K_f \frac{l^3}{m^3} \tag{4.2.10}$$

$$K_{h,m} = K_h \frac{l}{m} \tag{4.2.11}$$

$$K_{s,m} = K_s \frac{l}{m} \tag{4.2.12}$$

From now on, the forces and displacements under analysis is considered to be small. The equations describe the instantaneous relationships for forces and displacements. The loading force $F$ on each edge causes stress $\sigma$ in Figure S12. Assume that the thickness (into the page) is $b$, then

$$\sigma = \frac{F}{bl \sin \phi} \tag{4.2.13}$$

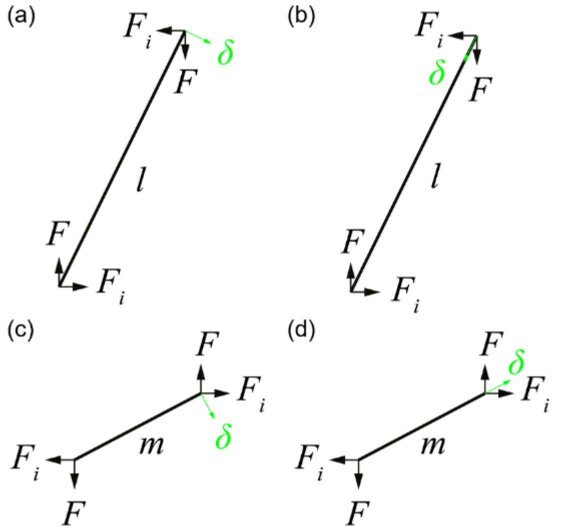


**Figure S13.** Schematics of forces ($F$ and $F_i$) and corresponding displacements ($\delta$) on the edges of a re-entrant triangle. $F_i$ is the internal force caused by different $K$s and orientations of $l$ and $m$. (a) Flexure and hinging displacements of $l$. (b) Stretching displacement of $l$. (c) Flexure and hinging displacements of $m$. (d) Stretching displacement of $m$.

Due to different $K$s and orientations of $l$ and $m$, an internal force $F_i$ keeps the structure from breaking at the joint when subjected to an external load $F$. From Figure S13, the displacements for $l$ may be listed as shown below. Flexure displacement:

$$\delta_f = \frac{F \sin \phi - F_i \cos \phi}{K_f} \tag{4.2.14}$$

Hinging displacement:

$$\delta_h = \frac{F \sin \phi - F_i \cos \phi}{K_h} \tag{4.2.15}$$

Stretching displacement:

$$\delta_s = \frac{F\cos\phi + F_i \sin\phi}{K_s} \tag{4.2.16}$$

Similarly, displacements for *m* can also be obtained.

$$\delta_{f,m} = \frac{-F\sin\theta + F_i \cos\theta}{K_{f,m}} \tag{4.2.17}$$

$$\delta_{h,m} = \frac{-F\sin\theta + F_i \cos\theta}{K_{h,m}} \tag{4.2.18}$$

$$\delta_{s,m} = \frac{F\cos\theta + F_i \sin\theta}{K_{s,m}} \tag{4.2.19}$$

**S4.2.2 Internal force and structural displacement of the re-entrant triangle**

Since the structure shall not break under the external load (*i.e.*, compatibility condition), the displacement along the *y* direction must be the same for *l* and *m*.

$$\Delta y_l = \Delta y_m \tag{4.2.20}$$

where

$$\Delta y_l = -\left(\delta_f + \delta_h\right)\cos\phi + \delta_s \sin\phi \tag{4.2.21}$$

$$\Delta y_m = -\left(\delta_{f,m} + \delta_{h,m}\right)\cos\theta - \delta_{s,m}\sin\theta \tag{4.2.22}$$

Using equations (4.2.10) through (4.2.12) and (4.2.14) to (4.2.22), $F_i/F$ is solved.

$$\frac{F_i}{F} = -\frac{\left(\frac{1}{K_s} - \frac{1}{K_f} - \frac{1}{K_h}\right)\sin\phi\cos\phi - \frac{m}{l}\left(\frac{m^2}{l^2}\frac{1}{K_f} + \frac{1}{K_h} - \frac{1}{K_s}\right)\sin\theta\cos\theta}{\left(\frac{1}{K_f} + \frac{1}{K_h}\right)\cos^2\phi + \frac{1}{K_s}\left(\sin^2\phi + \frac{m}{l}\sin^2\theta\right) + \frac{m}{l}\left(\frac{m^2}{l^2}\frac{1}{K_f} + \frac{1}{K_h}\right)\cos^2\theta} \tag{4.2.23}$$

By geometry, *x* and the relationship between the angles ($\gamma$, $\theta$, $\phi$) can be expressed.

$$x = \sqrt{l^2 - 2lm\cos\gamma + m^2} \tag{4.2.24}$$

$$\sin\theta = \frac{l\sin\gamma}{x} \tag{4.2.25}$$

$$\cos\theta = \frac{l\cos\gamma - m}{x} \tag{4.2.26}$$

$$\sin\phi = \frac{m\sin\gamma}{x} \tag{4.2.27}$$

$$\cos\phi = \frac{l - m\cos\gamma}{x} \tag{4.2.28}$$

The displacement in the *x* direction is expressed.

$$\Delta x = \Delta x_l + \Delta x_m \tag{4.2.29}$$

where

$$\Delta x_l = \left(\delta_f + \delta_h\right)\sin\phi + \delta_s\cos\phi \tag{4.2.30}$$

$$\Delta x_m = -\left(\delta_{f,m} + \delta_{h,m}\right)\sin\theta + \delta_{s,m}\cos\theta \tag{4.2.31}$$

Note that $\theta$ and $\phi$ are functions of $\gamma$ only, thus $F_i/F$ is also a function of $\gamma$ only. As shown by equations (4.2.14) to (4.2.19), all the displacements divided by loading force, such as $\delta_f/F$, are all functions of $\gamma$ only. Since the forces and displacements under analysis are considered to be small, the structural displacements divided by the loading force may be noted as $\Delta y/\Delta F$ and $\Delta x/\Delta F$, as shown below in equations (4.2.32). Considering equations (4.2.20) to (4.2.22) and (4.2.29) to (4.2.31), both $\Delta y/\Delta F$ and $\Delta x/\Delta F$ are also functions of $\gamma$ only.

**S4.2.3 Calculation of accumulated forces $F_{\mathrm{acc}}$ and $F_{i,\mathrm{acc}}$ of the re-entrant triangle**

Since we consider structural deformation from an initial angle ($\gamma_i$) to a final angle ($\gamma_f$), the accumulated, total forces are calculated. Since $\Delta x$ and $\Delta F$ are small,

$$\frac{\mathrm{d}x}{\mathrm{d}F} \approx \frac{\Delta x}{\Delta F} \tag{4.2.32}$$

From equation (4.2.24), the rate of structural displacement in $x$ direction with respect to $\gamma$ is calculated.

$$\frac{\mathrm{d}x}{\mathrm{d}\gamma}=\frac{lm}{x}\sin\gamma \tag{4.2.33}$$

With the chain rule,

$$\frac{\mathrm{d}F}{\mathrm{d}\gamma}=\frac{\mathrm{d}x}{\mathrm{d}\gamma}\cdot\left(\frac{\mathrm{d}x}{\mathrm{d}F}\right)^{-1} \tag{4.2.34}$$

By assuming quasi equilibrium deformation, $F_{\mathrm{acc}}$ is then calculated by integration.

$$F_{\mathrm{acc}}=\int_{\gamma_i}^{\gamma_f}\frac{\mathrm{d}F}{\mathrm{d}\gamma}\cdot\mathrm{d}\boldsymbol{\gamma} \tag{4.2.35}$$

where

$$\mathrm{d}\boldsymbol{\gamma}=\begin{cases}\mathrm{d}\gamma & \text{if } \gamma_f\geq\gamma_i\\ -\mathrm{d}\gamma & \text{if } \gamma_f<\gamma_i\end{cases} \tag{4.2.36}$$

Similarly,

$$\frac{\mathrm{d}F_i}{\mathrm{d}\gamma}=\frac{\mathrm{d}x}{\mathrm{d}\gamma}\cdot\left(\frac{\mathrm{d}x}{\mathrm{d}F_i}\right)^{-1}=\frac{\mathrm{d}x}{\mathrm{d}\gamma}\cdot\frac{F_i}{F}\cdot\left(\frac{\mathrm{d}x}{\mathrm{d}F}\right)^{-1} \tag{4.2.37}$$

$$F_{i,\mathrm{acc}}=\int_{\gamma_i}^{\gamma_f}\frac{\mathrm{d}F_i}{\mathrm{d}\gamma}\cdot\mathrm{d}\boldsymbol{\gamma} \tag{4.2.38}$$

#### S4.2.4 Calculation of structural Young's modulus of the re-entrant triangle

For Young's modulus, accumulated strain and stress are needed. The total displacement and strain in $x$ direction from $\gamma_i$ to $\gamma_f$ are derived from equation (4.2.24).

$$\Delta x_{\mathrm{acc}}=\left|x(\gamma_i)-x(\gamma_f)\right| \tag{4.2.39}$$

$$\varepsilon_{1,\mathrm{acc}}=\frac{\Delta x_{\mathrm{acc}}}{x(\gamma_i)} \tag{4.2.40}$$

Equation (4.2.13) is used for the calculation of the stress.

$$\sigma_{\mathrm{acc}}=\frac{F_{\mathrm{acc}}}{bl\sin(\phi_f)} \tag{4.2.41}$$

where $\sin(\phi_f)$ is calculated from equation (4.2.27) using $\gamma_f$. Structural Young's modulus is then

$$E=\frac{\sigma_{\mathrm{acc}}}{\varepsilon_{1,\mathrm{acc}}}=\frac{F_{\mathrm{acc}}}{bl\sin(\phi_f)}\cdot\frac{x(\gamma_i)}{\Delta x_{\mathrm{acc}}} \tag{4.2.42}$$

#### S4.2.5 Calculation of accumulated flexure of the re-entrant triangle

Equation (4.2.14) indicates the relationship between flexure ($\delta_f$) and forces ($F$ and $F_i$). By taking the derivative with respect to angle $\gamma$ on both sides and considering $F$ and $F_i$ are small, we obtain

$$\frac{\mathrm{d}\delta_f}{\mathrm{d}\gamma}\approx\frac{1}{K_f}\left(\frac{\mathrm{d}F}{\mathrm{d}\gamma}\sin\phi-\frac{\mathrm{d}F_i}{\mathrm{d}\gamma}\cos\phi\right) \tag{4.2.43}$$

The accumulated, total flexure $\delta_{f,\mathrm{acc}}$ can then be calculated by integration.

$$\delta_{f,\mathrm{acc}}=\int_{\gamma_i}^{\gamma_f}\frac{\mathrm{d}\delta_f}{\mathrm{d}\gamma}\cdot\mathrm{d}\boldsymbol{\gamma} \tag{4.2.44}$$

Note that $\delta_{f,\mathrm{acc}}$ is negative after calculation, indicating that its actual direction is the opposite to Figure S13a. In Figure 3, the actual direction and positive value of flexure are plotted.

#### S4.2.6 Calculation of Poisson's ratio of the re-entrant triangle

Poisson's ratio is defined as a negative ratio between strain in $y$ and $x$ directions.

$$\nu(\gamma)=-\frac{\varepsilon_2}{\varepsilon_1}=-\frac{\Delta y/y}{\Delta x/x}=-\frac{x}{y}\cdot\frac{\Delta y}{\Delta x} \tag{4.2.45}$$

where

$$\Delta y = 2\Delta y_l \tag{4.2.46}$$
$$y = 2l\sin\phi \tag{4.2.47}$$
Since $\Delta y/\Delta F$ and $\Delta x/\Delta F$ are functions of $\gamma$ only, Poisson's ratio is also a function of $\gamma$ only.

#### S4.2.7 Deformation behavior of the rotating square

The rotating square is different from the re-entrant triangle due to its centrosymmetric structure. The forces on each edge and the deformation behavior are considered.

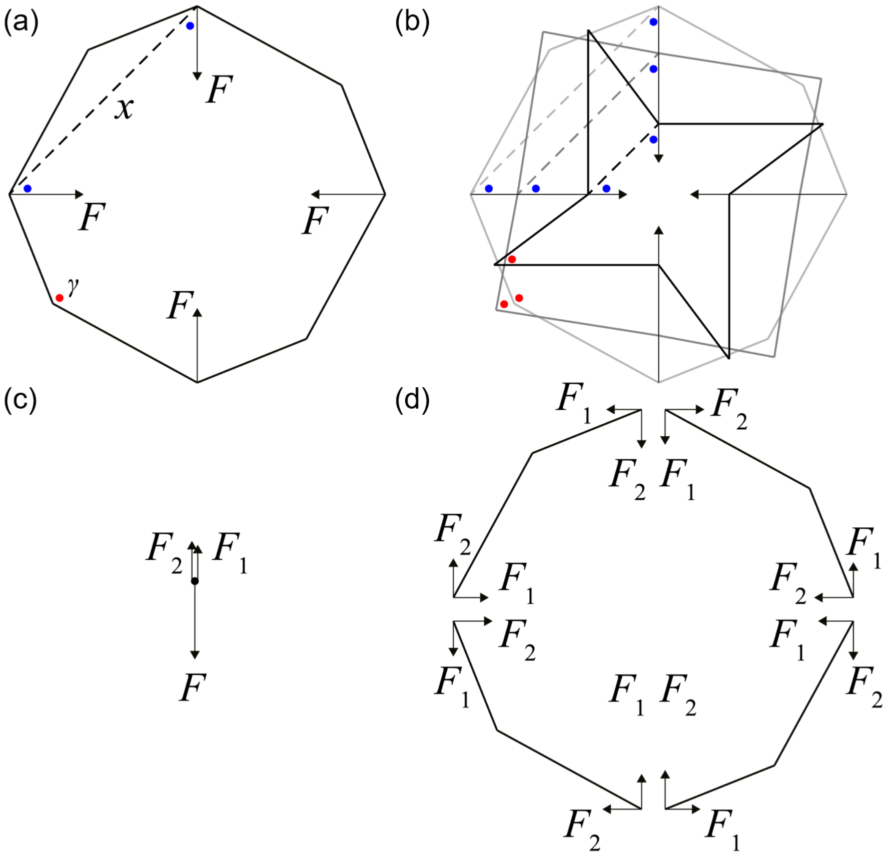


**Figure S14.** A single unit of the rotating square. The $x$ is marked as a dashed line. The angle $\gamma$ is marked by red dots and the angles noted with blue dots are all π/4 or 45°. (a) The single unit under loading of four $F$s. (b) The deformation of the rotating square from $\gamma$ = 140° (light gray) to 88° (gray) to 37° (black). The arrows indicate the moving direction of the hinging points between four quarter subunits. Note that during deformation, the orientation of $x$ remains the same and the angles noted with blue dots also maintain at 45°. (c) Force balance in vertical direction at the upper hinging point between the upper left and the upper right quarter subunits. $F_2$ is the force from the upper left quarter subunit, while $F_1$ is from the upper right quarter subunit. $F$ is the applied loading. As a result, $F = F_1 + F_2$. (d) Force balance of four quarter subunits separated at the hinging points. All four subunits are centrosymmetric not only in geometry, but also in mechanical loading. For example, $F_1$ is the horizontal force and $F_2$ is the vertical force in the upper left and the lower right quarter subunits, while $F_2$ is horizontal and $F_1$ is vertical in the upper right and the lower left quarter subunits.

The angles between the loading direction and the $x$ direction (*i.e.*, dashed lines) are all π/4 or 45°. During the deformation, these angles as well as the $x$ directions remain the same (blue dots in Figure S14). The distribution of the load $F$ on each edge may not be equal, and thus, noted as $F_1$ and $F_2$ in Figure S14 with the condition $F = F_1 + F_2$. As the force balance of each quarter subunit is analyzed, the assumption on $F_1$ and $F_2$ is compatible with the centrosymmetric condition. Note that: (i) under the centrosymmetric condition,

Poisson's ratio is always -1, and (ii) if $F_1 < F_2$, there will be a clockwise moment (*i.e*. torque) on the rotating square, causing a rigid-body rotation as the structure deforms (*vice versa*).

**S4.2.8 Forces and displacements of the rotating square**

Force constants in the rotating square are defined in the same way as in the re-entrant triangle, thus equations (4.2.2), (4.2.3) and (4.2.6) through (4.2.12) are applicable on the rotating square.

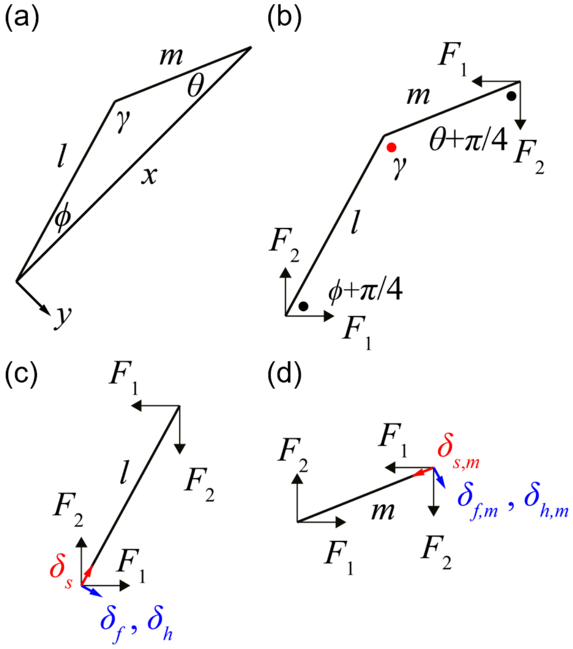


**Figure S15.** Schematics of one quarter subunit of the rotating square. Forces ($F_1$ and $F_2$) and corresponding displacements ($\delta_f$, $\delta_h$, $\delta_s$, $\delta_{f,m}$, $\delta_{h,m}$, and $\delta_{s,m}$) are shown on the two edges $l$ and $m$. $F_1$ is the horizontal force while $F_2$ is the vertical force. (a) One quarter subunit of the rotating square with edges ($l$ and $m$) and angles ($\gamma$, $\theta$, $\phi$) marked. Note that the orientations of $x$ and $y$ are also indicated. (b) Forces ($F_1$ and $F_2$) and related angles on the quarter subunit. The angle $\gamma$ is noted as a red dot, while the angles, $\theta+\pi/4$ and $\phi+\pi/4$, are indicated as black dots. (c) – (d) Flexure, hinging, and stretching displacements under $F_1$ and $F_2$ on edge $l$ and $m$. The flexing and hinging displacements are marked in blue. The stretching displacements are shown in red.

Similar to the re-entrant triangle, the displacements from Figure S15 for $l$ and $m$ are listed as shown below.

$$\delta_f = \frac{F_1 \sin(\phi + \pi/4) - F_2 \cos(\phi + \pi/4)}{K_f} \tag{4.2.48}$$

$$\delta_h = \frac{F_1 \sin(\phi + \pi/4) - F_2 \cos(\phi + \pi/4)}{K_h} \tag{4.2.49}$$

$$\delta_s = \frac{F_1 \cos(\phi + \pi/4) + F_2 \sin(\phi + \pi/4)}{K_s} \tag{4.2.50}$$

$$\delta_{f,m} = \frac{-F_1 \cos(\theta + \pi/4) + F_2 \sin(\theta + \pi/4)}{K_{f,m}} \tag{4.2.51}$$

$$\delta_{h,m} = \frac{-F_1 \cos(\theta + \pi/4) + F_2 \sin(\theta + \pi/4)}{K_{h,m}} \tag{4.2.52}$$

$$\delta_{s,m} = \frac{F_1 \sin(\theta + \pi/4) + F_2 \cos(\theta + \pi/4)}{K_{s,m}} \tag{4.2.53}$$

From the compatibility condition, the displacement along the $y$ direction must be the same for $l$ and $m$. Thus equation (4.2.20) is applicable on the rotating square. Note that $\Delta y_l$ and $\Delta y_m$ are different from the re-entrant triangle.

$$\Delta y_l = (\delta_f + \delta_h)\cos\phi - \delta_s \sin\phi \tag{4.2.54}$$

$$\Delta y_m = (\delta_{f,m} + \delta_{h,m})\cos\theta - \delta_{s,m}\sin\theta \tag{4.2.55}$$

Now $F_1/F_2$ can be solved.

$$\frac{F_1}{F_2} = \frac{\left(\frac{1}{K_f} + \frac{1}{K_h}\right) A\cos\phi + \frac{1}{K_s}\left(B\sin\phi - \frac{m}{l} C\sin\theta\right) + \frac{m}{l}\left(\frac{m^2}{l^2}\frac{1}{K_f} + \frac{1}{K_h}\right) D\cos\theta}{\left(\frac{1}{K_f} + \frac{1}{K_h}\right) B\cos\phi + \frac{1}{K_s}\left(-A\sin\phi + \frac{m}{l} D\sin\theta\right) + \frac{m}{l}\left(\frac{m^2}{l^2}\frac{1}{K_f} + \frac{1}{K_h}\right) C\cos\theta} \tag{4.2.56}$$

where

$$A = \cos\phi - \sin\phi \tag{4.2.57}$$

$$B = \cos\phi + \sin\phi \tag{4.2.58}$$

$$C = \cos\theta - \sin\theta \tag{4.2.59}$$

$$D = \cos\theta + \sin\theta \tag{4.2.60}$$

By geometry, $x$ and the relationship between the angles ($\gamma$, $\theta$, $\phi$) can be expressed. Equations (4.2.24), (4.2.25), and (4.2.27) through (4.2.29) are applicable on the rotating square. The differences are $\cos\theta$, $\Delta x_l$ and $\Delta x_m$.

$$\cos\theta = \frac{m - l\cos\gamma}{x} \tag{4.2.61}$$

$$\Delta x_l = (\delta_f + \delta_h)\sin\phi + \delta_s\cos\phi \tag{4.2.62}$$

$$\Delta x_m = (\delta_{f,m} + \delta_{h,m})\sin\theta + \delta_{s,m}\cos\theta \tag{4.2.63}$$

$\theta$, $\phi$, $F_1/F_2$, and all the displacements divided by $F_2$ (such as $\delta_f/F_2$) are all functions of $\gamma$ only. Since the forces and displacements under analysis are considered to be small, the structural displacements divided by $F_2$ may be noted as $\Delta y/\Delta F_2$ and $\Delta x/\Delta F_2$ which are also functions of $\gamma$ only.

#### S4.2.9 Calculation of accumulated forces ($F_{1,\mathrm{acc}}$ and $F_{2,\mathrm{acc}}$) and flexure of the rotating square

Since the mechanical properties for structural reconfiguration from an initial angle ($\gamma_i$) to a final angle ($\gamma_f$), the accumulated, total forces are of interest to our analysis. Since $\Delta x$ and $\Delta F$ are small,

$$\frac{\mathrm{d}x}{\mathrm{d}F_2} \approx \frac{\Delta x}{\Delta F_2} \tag{4.2.64}$$

With the chain rule,

$$\frac{\mathrm{d}F_2}{\mathrm{d}\gamma} = \frac{\mathrm{d}x}{\mathrm{d}\gamma}\cdot\left(\frac{\mathrm{d}x}{\mathrm{d}F_2}\right)^{-1} \tag{4.2.65}$$

Thus, $F_{\mathrm{acc}}$ is calculated by integration.

$$F_{2,\mathrm{acc}} = \int_{\gamma_i}^{\gamma_f} \frac{\mathrm{d}F_2}{\mathrm{d}\gamma}\cdot \mathrm{d}\boldsymbol{\gamma} \tag{4.2.66}$$

Similarly,

$$\frac{\mathrm{d}F_1}{\mathrm{d}\gamma} = \frac{\mathrm{d}x}{\mathrm{d}\gamma}\cdot\left(\frac{\mathrm{d}x}{\mathrm{d}F_1}\right)^{-1} = \frac{\mathrm{d}x}{\mathrm{d}\gamma}\cdot\frac{F_1}{F_2}\cdot\left(\frac{\mathrm{d}x}{\mathrm{d}F_2}\right)^{-1} \tag{4.2.67}$$

$$F_{1,\mathrm{acc}} = \int_{\gamma_i}^{\gamma_f} \frac{\mathrm{d}F_1}{\mathrm{d}\gamma}\cdot \mathrm{d}\boldsymbol{\gamma} \tag{4.2.68}$$

Since $F = F_1 + F_2$ is always valid,

$$\frac{\mathrm{d}F}{\mathrm{d}\gamma} = \frac{\mathrm{d}F_1}{\mathrm{d}\gamma} + \frac{\mathrm{d}F_2}{\mathrm{d}\gamma} \tag{4.2.69}$$

$$F_{\mathrm{acc}} = \int_{\gamma_i}^{\gamma_f}\left(\frac{\mathrm{d}F_1}{\mathrm{d}\gamma} + \frac{\mathrm{d}F_2}{\mathrm{d}\gamma}\right)\cdot \mathrm{d}\boldsymbol{\gamma} = F_{1,\mathrm{acc}} + F_{2,\mathrm{acc}} \tag{4.2.70}$$

Equation (4.2.48) indicate the relationships between flexure ($\delta_f$) and forces ($F_1$ and $F_2$). By taking the derivative with respect to angle $\gamma$ on both sides and considering $F_1$ and $F_2$ are small,

$$\frac{\mathrm{d}\delta_f}{\mathrm{d}\gamma} \approx \frac{1}{\sqrt{2}K_f}\left[\frac{\mathrm{d}F_1}{\mathrm{d}\gamma}(\cos\phi + \sin\phi) - \frac{\mathrm{d}F_2}{\mathrm{d}\gamma}(\cos\phi - \sin\phi)\right] \quad (4.2.71)$$

$\delta_{f,\mathrm{acc}}$ can then be calculated by integration.

$$\delta_{f,\mathrm{acc}} = \int_{\gamma_i}^{\gamma_f} \frac{\mathrm{d}\delta_f}{\mathrm{d}\gamma} \cdot \mathrm{d}\boldsymbol{\gamma} \quad (4.2.72)$$

**S4.2.10 Calculation of structural Young's modulus of the rotating square**

Accumulated, total strain and stress for entire deformation are needed to calculate Young's modulus. The accumulated displacement and strain from $\gamma_i$ to $\gamma_f$ are derived from equation (4.2.24).

$$\Delta x_{\mathrm{acc}} = \left|x(\gamma_i) - x(\gamma_f)\right| \quad (4.2.73)$$

$$\varepsilon_{\mathrm{acc}} = \frac{\Delta x_{\mathrm{acc}}}{x(\gamma_i)} \quad (4.2.74)$$

As the structure is loaded in two orthogonal directions, the stress is defined differently from the unidirectional loading. Here, we use the sum of two $F$s projected onto the perpendicular direction of $x$.

$$\sigma_{\mathrm{acc}} = \frac{2F_{\mathrm{acc}}}{bx(\gamma_f)}\sin\left(\frac{\pi}{4}\right) = \frac{\sqrt{2}F_{\mathrm{acc}}}{bx(\gamma_f)} \quad (4.2.75)$$

Structural Young's modulus is then

$$E = \frac{\sigma_{\mathrm{acc}}}{\varepsilon_{\mathrm{acc}}} = \frac{\sqrt{2}F_{\mathrm{acc}}}{bx(\gamma_f)} \cdot \frac{x(\gamma_i)}{\Delta x_{\mathrm{acc}}} \quad (4.2.76)$$

**S4.2.11 Moment of force and rigid-body rotation of the rotating square**

If the clockwise direction is the positive direction in the quarter subunit of Figure S15, the moment of force is

$$M_{\mathrm{q}} = F_2 \cdot x/\sqrt{2} - F_1 \cdot x/\sqrt{2} = x(F_2 - F_1)/\sqrt{2} \quad (4.2.77)$$

In the whole unit of Figure S14, the moment of force is

$$M_{\mathrm{all}} = 4 \cdot (F_2 - F_1) \cdot x/\sqrt{2} = 2\sqrt{2}x(F_2 - F_1) \quad (4.2.78)$$

Comparing equations (4.2.77) and (4.2.78), $M_{\mathrm{all}} = 4\, M_{\mathrm{q}}$. This suggests that the clockwise moment has been taken into consideration in each quarter subunit, and thus, the total moment should not cause deformations other than those described in equations (4.2.48) to (4.2.53). The moment can only cause the rigid-body rotation of the entire structure.

#### S4.3 Criteria on the edge rigidity

Ideally, edges shall be straight. Due to the finite rigidity, however, perfectly straight edges may not be possible under loads. The edges should rather be kept near straight.

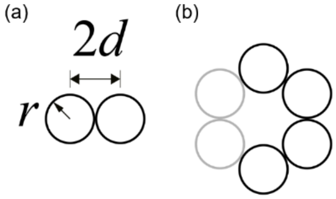


**Figure S16.** Schematics of cross-section of edges: $r$ is the radius of one dsDNA; $d$ is the half distance between the centers of two neighboring dsDNA. $r = 1.1$ nm and $2d = 2.3$ nm are assumed to be invariable regardless of cross-section design. (a) 2-dsDNA edge cross-section design. (b) 4-dsDNA cross-section. The two gray circles in (b) are imaginary bundles in a hexagonal arrangement.

##### S4.3.1 Criterion from buckling

Buckling is an unstable state of an edge, showing severe curvature. Hence, it needs to be avoided. Critical force load for buckling is define as[62]

$$F_{cr} = \frac{\pi^2 E_s I_{\min}}{l^2} = \frac{\pi^2 E_s A}{\left(l/r_g\right)^2} \quad (4.3.1)$$

where $A$ is the cross-section area of the edge and radius of gyration is

$$r_g = \sqrt{I_{\min}/A} \quad (4.3.2)$$

Since the edge can be bent in different ways, there will be a series of second moment of area $I$. $I_{\min}$ is the minimum value among them, indicating the easiest way for the edge to bend. To avoid buckling, $r_g/l$ shall not be too small. Normally, $r_g/l \geq 0.01$.[45,63]

$$\sqrt{I_{\min}/A}/l \geq 0.01 \quad (4.3.3)$$

The geometrical parameters are listed in table S4.1.

**Table S4.1.** Geometrical calculation of designs. Green indicates good agreement with the criterion. Orange implies marginal agreement. Red means that the value does not agree with the criterion.

| | honeycomb | | re-entrant triangle | | rotating square | full hexagon | |
|---|---|---|---|---|---|---|---|
| $A$ (nm$^2$) | 7.60 | | | | 15.2 | 22.8 | |
| $I_{\min}$ (nm$^4$) | 2.30 | | | | 19.7 | 66.0 | |
| $r_g$ (nm) | 0.55 | | | | 1.14 | 1.70 | |
| $l$ (nm) | 59 | 21 | 79 | 87 | 49 | 79 | 59 |
| $r_g/l$ | 0.009 | 0.026 | 0.007 | 0.006 | 0.023 | 0.022 | 0.029 |

##### S4.3.2 Criterion from flexing

Even when an edge is far from buckling, it can still have a strong flexure displacement. To guarantee a straight edge, flexing shall be small. Since the edge is far from buckling, the load is assumed to be small, for example, 10% of the critical loads for buckling in the flexing direction. We also consider the loads that contribute directly to the flexure.

$$F_f = \frac{\pi^2 E_s I}{10 \cdot l^2} \quad (4.3.4)$$

By using equations (4.2.1) and (4.2.2),

$$\delta_f = \frac{F_f}{K_f} = \frac{\pi^2 E_s I}{10 \cdot l^2} \cdot \frac{l^3}{12 E_s I} = \frac{\pi^2 l}{120} \quad (4.3.5)$$

where $\delta_f$ is the flexure, which shall be less than the thickness ($t$) of the edge in order to keep the edge straight.

$$\frac{\pi^2 l}{120} = \delta_f \leq t \tag{4.3.6}$$

Hence,

$$\frac{t}{l} \geq \frac{\pi^2}{120} \approx 0.1 \tag{4.3.7}$$

The geometrical information can be extracted from the cross-section and length designs, which is listed in table S4.2. Note that $t_{max}$ is used to simplify the directional dependence.

**Table S4.2.** Geometrical calculation of designs. Green indicates good agreement with the criterion; orange implies marginal agreement; red means poor agreement with the criterion.

| | honeycomb | | re-entrant triangle | | rotating square | full hexagon | |
|---|---|---|---|---|---|---|---|
| $t_{min}$ (nm) | 2.2 | | | | 4.2 | 6.2 | |
| $t_{max}$ (nm) | 4.5 | | | | 6.8 | 6.8 | |
| $l$ (nm) | 59 | 21 | 79 | 87 | 49 | 79 | 59 |
| $t_{min}/l$ | 0.04 | 0.1 | 0.03 | 0.02 | 0.08 | 0.08 | 0.1 |
| $t_{max}/l$ | 0.08 | 0.2 | 0.06 | 0.05 | 0.1 | 0.09 | 0.1 |

#### S4.3.3 Evaluation of flexure

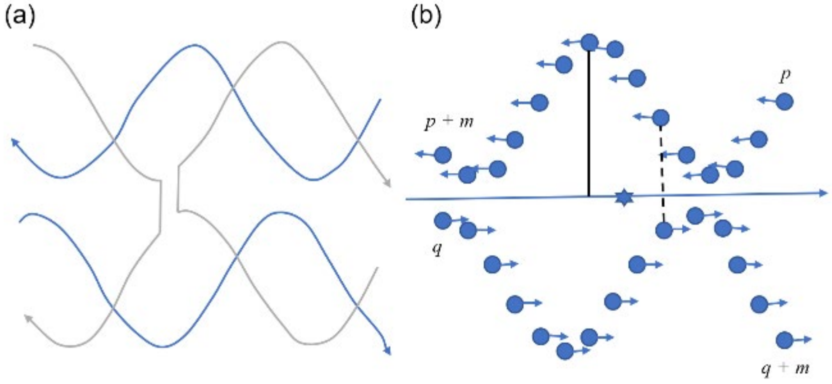


**Figure S17.** Detailed calculation of flexure on an edge. (a) A fraction of an edge. Blue represents the scaffold segment and gray denotes staples. The arrow indicates the 3' ends. (b) Phosphate residues on the backbone of the scaffold segments in (a). The blue dot represents the phosphate residue, and the blue star shows the center of all phosphate residues. The short blue arrow represents the normal vector of the base. The long blue arrow represents the averaged normal vector of all the bases. The black solid line indicates the distance from a phosphate residue to the center line. The black dashed line indicates the distance from a phosphate residue on one side of the scaffold to the corresponding phosphate residue on the other side of the scaffold. There are ($m$ + 1) phosphate residue on either side of the scaffold. The upper side starts at number $p$ and ends at number ($p + m$), while the lower counts from $q$ to ($q + m$).

As shown in Figure S17a, each edge is composed of scaffold segments and staples. Since they are bonded to each other in a consistent way, only the scaffold is considered in the calculation of flexure (Figure S17b). Two key parameters needed in the calculation are: the averaged normal vector of all the bases ($\mathbf{n}_{\text{ave}}$) and the center of all phosphate residues ($\mathbf{r}_{\text{ave}}$), which can be determined as shown below. Since the upper and lower scaffold segments point at opposite directions, a subtraction is used in the calculation of average normal vector.

$$\boldsymbol{n}_{\text{ave}} = \frac{1}{2 \cdot (m+1)} \sum_{i=0}^{m} \left(\boldsymbol{n}_{p+i} - \boldsymbol{n}_{q+i}\right) \tag{4.3.8}$$

$$\boldsymbol{r}_{\text{ave}} = \frac{1}{2 \cdot (m+1)} \sum_{i=0}^{m} \left(\boldsymbol{r}_{p+i} + \boldsymbol{r}_{q+i}\right) \tag{4.3.9}$$

With knowledge of the averaged point ($\boldsymbol{r}_{\text{ave}}$) and direction ($\boldsymbol{n}_{\text{ave}}$), a center line can be created. Thus, distances from all phosphate residues to the center line ($d_{\text{c},i}$) are calculated.

To obtain the accurate flexure of the edge, its thickness must be taken into consideration. As the number of nucleotides on either chain of the scaffold is the same for any given edge, corresponding pairs of phosphate residues from either chain can be found. The thickness of the edge ($t_{\text{edge}}$) is the maximum value of all the distances between corresponding phosphate residues.

$$t_{\text{edge}} = \max_{i=0 \text{ to } i=m} \left| \boldsymbol{r}_{p+i} - \boldsymbol{r}_{q+m-i} \right| \tag{4.3.10}$$

Finally, the calculated flexure is given by

$$\delta_{f,\text{cal}} = 2 \cdot \max_{i=0 \text{ to } i=m} \left( d_{\text{c},i} \right) - t_{\text{edge}} \tag{4.3.11}$$

For the evaluation of an edge,

$$\frac{\delta_{f,\text{cal}}}{l} \leq 0.1 \tag{4.3.12}$$

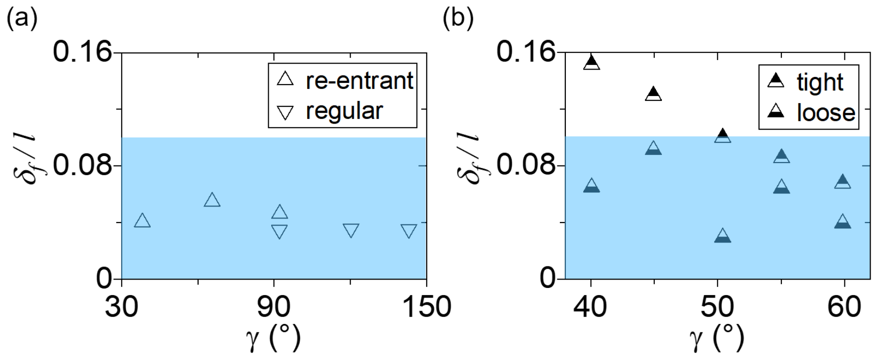


**Figure S18.** Dimensionless flexure of different structures from MD simulations. (a) Honeycombs. With short $l$, $\delta_f/l$ is within the recommended range (blue shade). (b) Control design of re-entrant triangle. The design includes two types of joints: tight and loose. Accordingly, edges are classified as tight and loose to indicate the joints that they are connected to. With long $l$ and tight joints, half of the data points in (b) are outside the recommended range.

Figure S18 shows that the control re-entrant triangle is mostly out of the range, which is due to long, thin edges. In contrast, the honeycombs are all within the requirement, thanks to short $l$.

### S4.4 Criteria on the joint stretch

A joint must be well defined and have considerable flexibility. Neither resting too loose nor holding too tight is appropriate. Thus, a degree of stretch is needed.

#### S4.4.1 Definition of deviational distance

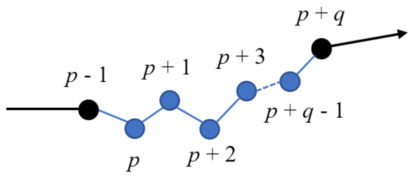


**Figure S19.** Schematic of phosphate residues on the backbone of a scaffold segment, including a joint. The part before number $p$ and after number ($p + q - 1$) phosphate residues is a double-stranded region (black), while the part in the middle is a single-stranded domain (blue), serving as a joint. The arrow indicates the 3' end.

For a joint connected by ssDNA as shown in Figure S19, the deviational distance ($\Delta d$) is defined by the magnitude of displacement.

$$\Delta d = |\Delta \boldsymbol{r}| \tag{4.4.1}$$

where $\boldsymbol{r}$ is the position vector of the phosphate residue. For a ssDNA connection with $q$ nt, starting from $p$ nt, $\Delta d$ is given as

$$\Delta d = |\boldsymbol{r}_{p+q} - \boldsymbol{r}_{p-1}| \tag{4.4.2}$$

#### S4.4.2 Definition of stretch

The stretch ($\xi$) of a joint formed by ssDNA is defined as deviational distance ($\Delta d$) over distance ($\Delta s$).

$$\xi = \frac{\Delta d}{\Delta s} \tag{4.4.3}$$

For a ssDNA joint with a length of $q$ nt from $p$ nt on the scaffold (Figure S19), $\xi$ is defined as

$$\xi = \frac{|\boldsymbol{r}_{p+q} - \boldsymbol{r}_{p-1}|}{\sum_{i=p}^{i=p+q} |\boldsymbol{r}_i - \boldsymbol{r}_{i-1}|} \tag{4.4.4}$$

Some pairs of edges are connected by multiple ssDNA segments. The maximum $\xi$ of all segments is selected to represent the stretch between each edge pair as the rest of the segments are too loose to contribute to the stretch level significantly. The reasons for having multiple connection include linking the scaffold segments. More importantly, ssDNA segments between the same edges can take turns to act as the tightest so that the linkage is potentially favor for a wide range of angle.

#### S4.4.3 Criteria on stretch of the joints in the re-entrant triangle

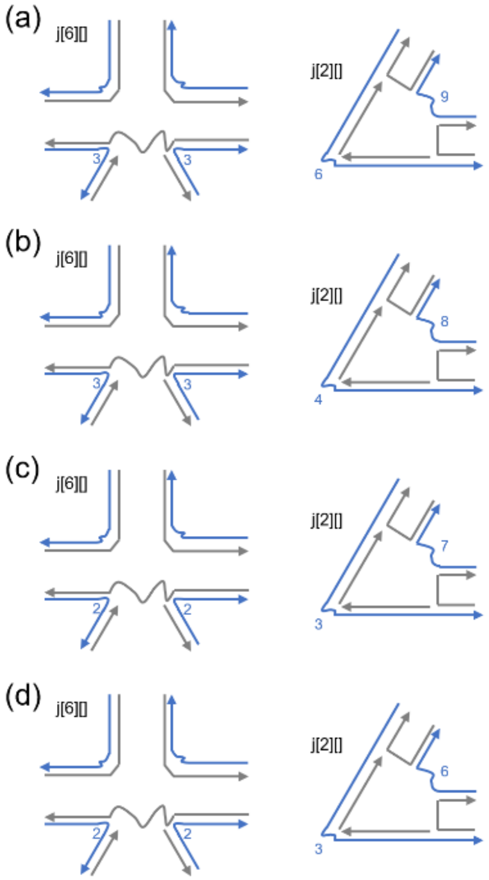


**Figure S20.** Schematics of joint designs in the re-entrant triangle. Here the length of the unpaired scaffold segment is varied, which are different from the designs in Figure S5b and d. Different length yields different stretch: (a) 55 %, (b) 60 %, (c) 65 %, (d) 70 %.

To examine the various extent of stretch at a joint, the lengths of ssDNA scaffold domains are modified as shown in Figure S20 (also see Figure S5). Once the stretch is determined, $\Delta d$ and $\delta_f/l$ are extracted and plotted for comparison.

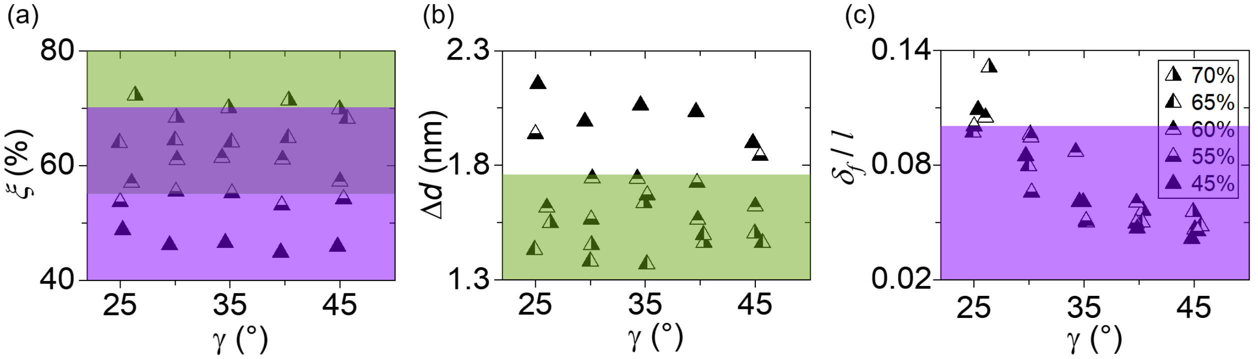


**Figure S21.** Computed stretch ($\xi$), deviational distance ($\Delta d$), and dimensionless flexure ($\delta_f/l$) of the re-entrant triangle. (a)–(c) share the same figure legends, shown in (c). The olive region in (a) and (b) indicates the desired region for $\xi$ based on $\Delta d$. The purple area in (a) and (c) shows the preferred range for $\xi$ based on $\delta_f/l$. The common area of olive and purple in (a) marks the recommended range for $\xi$, which is from 55 % to 70 %.

To maintain structural accuracy, $\Delta d$ must not be too large. For example, it shall be less than the minimum thickness of the edge ($t_{\min}$). Thus, we propose

$$\Delta d \leq 0.8 \cdot t_{\min} \tag{4.4.5}$$

In the re-entrant triangle, $\Delta d \leq 0.8 \times 2.2$ nm = 1.76 nm. As shown in Figure S21b, $\xi = 45$ % is outside the range; $\xi = 55$ % is on the boundary. Therefore, $\xi \geq 55$ % is reasonable in Figure S21a.

To ensure enough flexibility at the joint, $\delta_f/l$ of the connected edges must not be significant. Similar to equation (4.3.12), here we propose 10 % to be the limit.

$$\delta_f/l \leq 10\ \% \tag{4.4.6}$$

Figure S21c shows that while $\xi = 70$ % exceeds the limit at small $\gamma$, most data are within the range. Thus, we set $\xi \leq 70$ % as the upper limit. By combining $\xi \geq 55$ % and $\xi \leq 70$ %, the final design recommendation is

$$55\% \leq \xi \leq 70\% \tag{4.4.7}$$

**S4.4.4 Verification of Criteria on other structures**

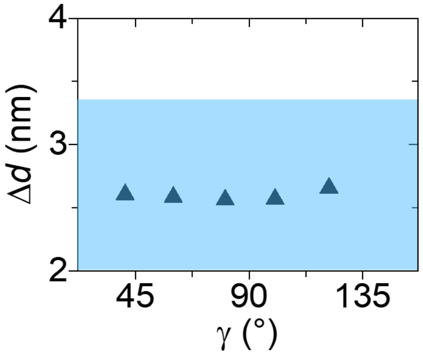


**Figure S22.** Deviational distance ($\Delta d$) of the rotating square from the MD simulations. Equation (4.4.5) yields $\Delta d \leq 0.8 \times 4.2$ nm = 3.36 nm for the rotating square. The result indicates that the reinforcement satisfies the design criterion on $\Delta d$.

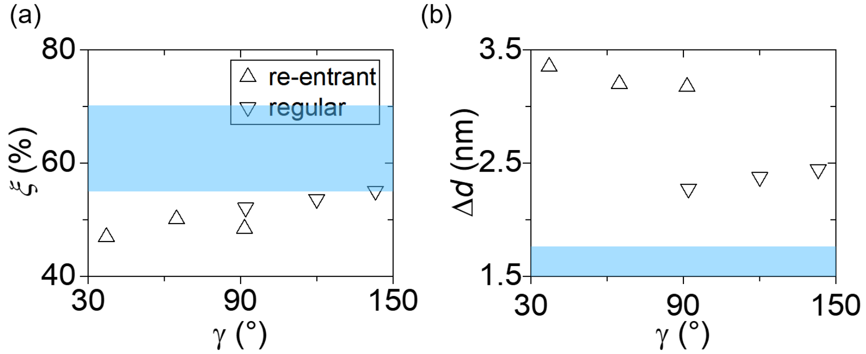


**Figure S23.** Computed stretch ($\xi$) and deviational distance ($\Delta d$) of honeycombs. (a)–(b) share the same figure legends, shown in (a). The blue shade indicates the recommended design requirements: 55 % $\leq \xi \leq$ 70 % and $\Delta d \leq 0.8 \times 2.2$ nm = 1.76 nm in both honeycombs. Note that equation (4.4.6) is already satisfied as shown in Figure S18a. With the data points out of the criterion on $\Delta d$, the honeycombs do not follow the design principles, thus showing loose joints and some curved edges (Figure 1g).

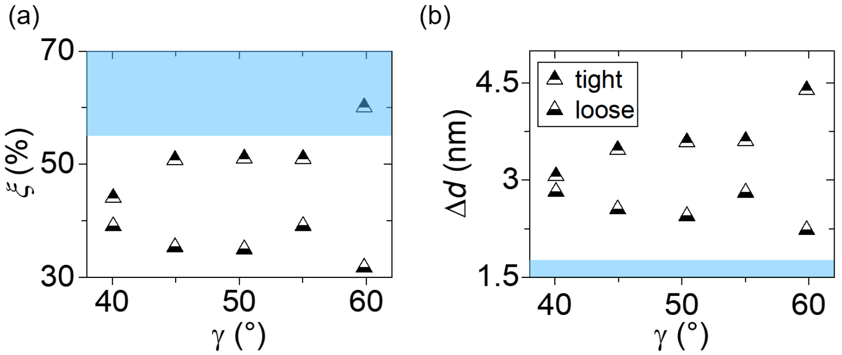


**Figure S24.** Computed stretch ($\xi$) and deviational distance ($\Delta d$) of the control design of the re-entrant triangle. The control design includes two types of joint designs: tight and loose joints. (a)–(b) share the same figure legends, shown in (b). The blue region indicates that the control design is mostly out of the recommended design criteria (55 % $\leq \xi \leq$ 70 %, $\Delta d \leq 0.8 \times 2.2$ nm = 1.76 nm). Note $\delta_f/l$ is partially out of the range set by equation (4.4.6) as shown in Figure S18b, thus its structural integrity is not maintained.

#### S4.5 Thermodynamic model for chemical deformation

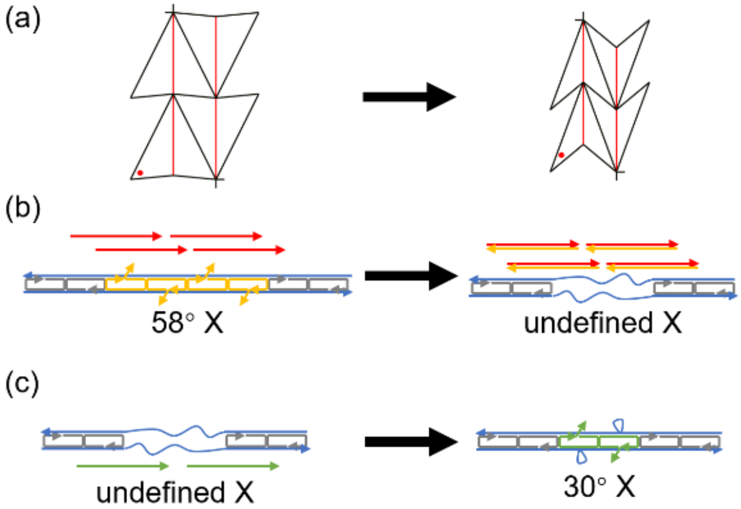


**Figure S25.** Schematics of chemical deformation of the re-entrant triangle from 58° to 30°. (a) The overall reconfiguration. (b) Release process via toehold-mediated strand displacement. A jack edge X will transition from 58° conformation to an undefined conformation after strand displacement of releasing strands (red) removing target strands with toeholds (orange). (c) Recombination process via reannealing. After removing the chemical waste (dsDNA in red and orange), new staples (green) are added and X will have a new length for 30° conformation.

The force needed for auxetic reconfiguration can be calculated theoretically using nearest neighbor model (NNM) and/or online tools such as NUPACK. As illustrated in Figure S25, the reconfiguration process includes two steps: (i) release of staples from jack edges and (ii) recombination of new staples for a different jack length. Considering the displacement of the edges occurs mostly in step (ii), the force is determined by the free energy change in the step.

$$F = \frac{-\Delta G}{\Delta x} \tag{4.5.1}$$

From the design, $\Delta x$ is about 14 nm per jack edge X. The free energy change ($\Delta G$) for each edge X may be determined by NNM and NUPACK. Both approaches yield similar results as shown in the chart below. Since the free energy change is the maximum energy available for reconfiguration, ~55 pN per edge X is the theoretical maximum force involved in chemical deformation.

**Table S4.3.** The free energy change ($\Delta G$) and theoretical maximum force per edge of the re-entrant triangle.

| Method | $\Delta G$ (/mol edge) | $\Delta G$ (J/edge) | $F$ (pN/edge) |
|---|---|---|---|
| NNM | -459.91 kJ | $-7.64\times10^{-19}$ | 55 |
| NUPACK | -108.51 kcal | $-7.54\times10^{-19}$ | 54 |

Similarly, theoretical maximum force involved in chemical deformation of the rotating square from 140° to 37° can be calculated using NNM and NUPACK calculations. $\Delta x$ is approximately 35 nm per jack edge A. The theoretical maximum force is ~72 pN per edge A.

**Table S4.4.** The free energy change ($\Delta G$) and theoretical maximum force per edge of the rotating square.

| Method | $\Delta G$ (/mol edge) | $\Delta G$ (J/edge) | $F$ (pN/edge) |
|---|---|---|---|
| NNM | -1449.4 kJ | $-2.41\times10^{-18}$ | 69 |
| NUPACK | -379.34 kcal | $-2.63\times10^{-18}$ | 75 |

### S5. MD Simulations with oxDNA

Coarse-grained (CG) molecular dynamics (MD) simulations were performed with oxDNA to investigate equilibrium conformations and mechanical deformations of DNA structures. Specifically, oxDNA2 model was used with the version 2.3, an update published in February 2018. The salt concentration was set to be $[Na^+] = 0.5$ M.[64] Since the DNA structures were deposited on mica surfaces at 4 °C, the simulations were also performed at 4 °C. The simulations of mechanical deformation were performed at 27 °C as the structure was reannealed from 40 °C to 20 °C.

Static structures in distinct conformations were created in caDNAno2, which were then converted into topology and configuration files for later use in oxDNA via 'cadnano_interface.py' using actual sequence information. Since the free loops of jack edges were rarely seen interacting with other edges under AFM, they were removed (by manually deleting them in the topology and configuration files) after the conversion. Then, the topology and configuration files were processed following the method described in 'NEW_RELAX_PROCEDURE'. After that, MD simulations were performed. The computations were in the range of $10^7$ steps and were suspended when the relative fluctuations were less than 3%. The total running time required for each conformation was about a week. After the simulations, the results were analyzed by extracting the directional and positional information. The results were also visualized in cogli1, a software designed to work with oxDNA.

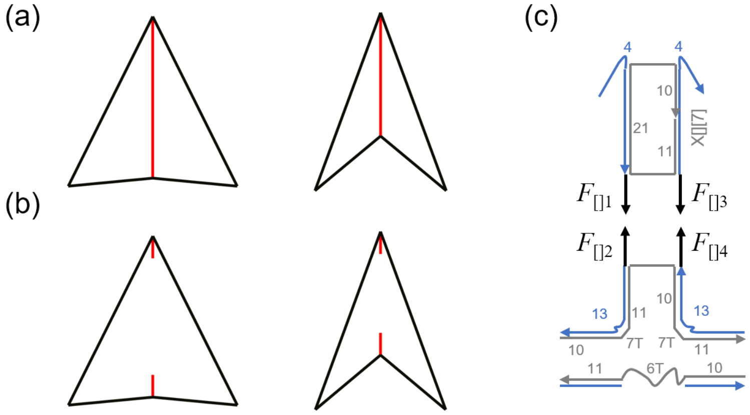


**Figure S26.** MD simulations for mechanical deformation of the re-entrant triangle. (a) One intact unit in the re-entrant triangle at 58° and 30°. (b) The unit in (a) with the jack edge X cut. Only staple X[][7] and corresponding scaffold segments were left. (c) Zoom-in view of the cut parts. Note that the joints are intact. To enable the dynamic reconfiguration, forces (black arrows, $F_{[]1}$ through $F_{[]4}$) were applied to the tip of the scaffold of X residue. The forces were pulling inward for the structural transition from 58° to 30° (as shown).

Figure S26 illustrates our MD simulations to study mechanical deformation of the re-entrant triangle. The X edges were removed (by manually deleting) so that the forces needed for structural deformation could be calculated. Harmonic traps were placed at the tip of all X residues to introduce pushing or pulling loads[64]. The stiffness of the traps was 5.7 pN/nm and the averaged (among all 4×4 traps) rate was $6.37\times10^6$ nm/s, yielding the force loading rate of $3.63\times10^7$ pN/s. Since the displacement of each X was small (~14 nm), a slower rate would not take too much time for simulations, while providing more stable forces.

When the simulations were complete, the forces were extracted by calculating the product of the trap stiffness and the displacement of each tip of the scaffold with respect to the corresponding harmonic trap. Since two pairs of forces ($F_{[]1}$ and $F_{[]2}$; $F_{[]3}$ and $F_{[]4}$) were applied on each X edge, the sum of the pairs or $(F_{[]1} + F_{[]2})/2 + (F_{[]3} + F_{[]4})/2$ is reported as the force per edge.

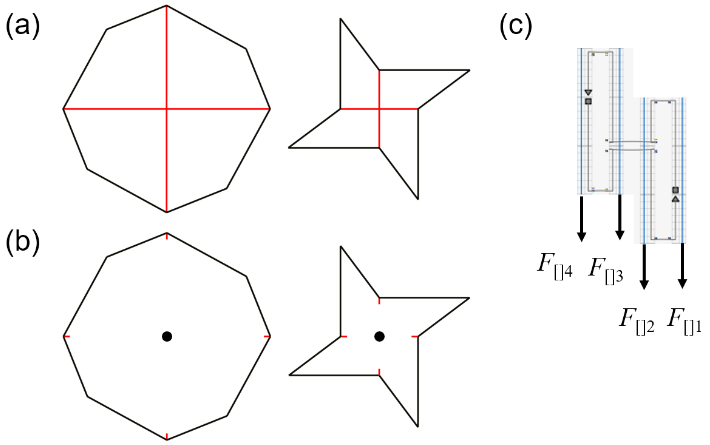


**Figure S27.** MD simulations for mechanical deformation of the rotating square. (a) The rotating square at 140° and 37°. (b) The jack edge As are removed. Only two staples close to the terminal of the jack and corresponding scaffold segments were left (which are shown in red). The center is noted as a black dot. (c) Zoom-in view of the part that has been cut. To enable the dynamic reconfiguration, forces (black arrows, $F_{1[]}$ through $F_{4[]}$) were applied to the tip of the scaffold of A residue. The forces were applied toward the center for the structural transition from 140° to 37° (as shown).

Figure S27 illustrates our simulation domain for the mechanical deformation of the rotating square. The jack edges (As) were removed for calculation of the forces needed for structural deformation. The stiffness of the harmonic traps used was also 5.7 pN/nm and the average (among all 4×4 traps) rate was $7.26\times10^6$ nm/s, yielding the force loading rate of $4.14\times10^7$ pN/s. Since the displacement of each A (~35 nm) was more than twice of the displacement of each X of the re-entrant triangle, a slightly faster rate was used to balance between simulation time and stable data on forces. Since four forces ($F_{[]1}$ through $F_{[]4}$) were pulling in the same direction on each A edge, the sum ($F_{1[]} + F_{2[]} + F_{3[]} + F_{4[]}$) is reported as the force per edge.

By using the harmonic traps, we also obtain the force perpendicular to the pulling direction (termed circular force, $F_{c,\mathrm{acc}}$). For each of the four forces in Figure S27, there is a corresponding circular force. For example, $F_{[]1c}$ is the circular force together with $F_{[]1}$. As noted in the discussion of Figure S14, this circular force is caused by the fact that $F_1 \neq F_2$. Note that $F_2 > F_1$, then in the simulation,

$$F_{[]1} + F_{[]2} + F_{[]3} + F_{[]4} = F = F_1 + F_2 \quad (5.1)$$

$$F_{[]1c} + F_{[]2c} + F_{[]3c} + F_{[]4c} = F_c = -F_1 + F_2 \quad (5.2)$$

$F_1$ and $F_2$ are then calculated.

$$F_1 = (F - F_c)/2 \quad (5.3)$$

$$F_2 = (F + F_c)/2 \quad (5.4)$$

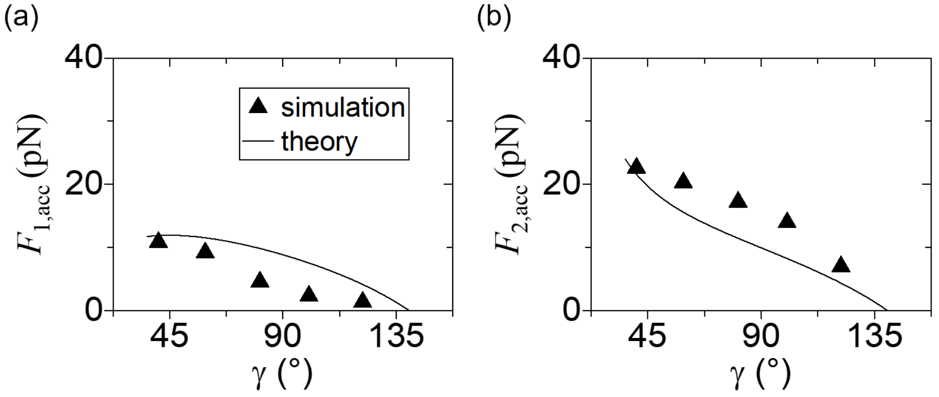

**Figure S28.** Accumulated forces ($F_{1,acc}$ and $F_{2,acc}$) in the rotating square as a function of angle $\gamma$. Solid lines represent theoretical predictions from the finite model, while solid triangles denote the simulation results of structural deformation by mechanical loading. The simulation data reasonably follows the theoretical predictions. The sum of (a) and (b) yields Figure 5c.

**S6. Supporting AFM Images**
**Re-entrant honeycomb**

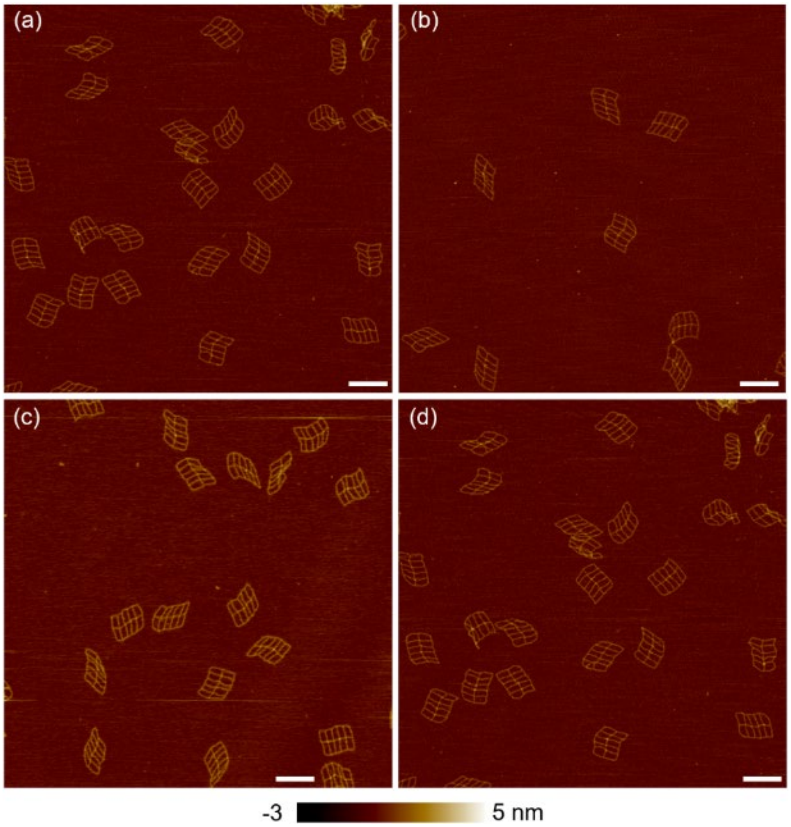


**Figure S29.** AFM images of the re-entrant honeycomb at 90°. Scale bar: 200 nm.

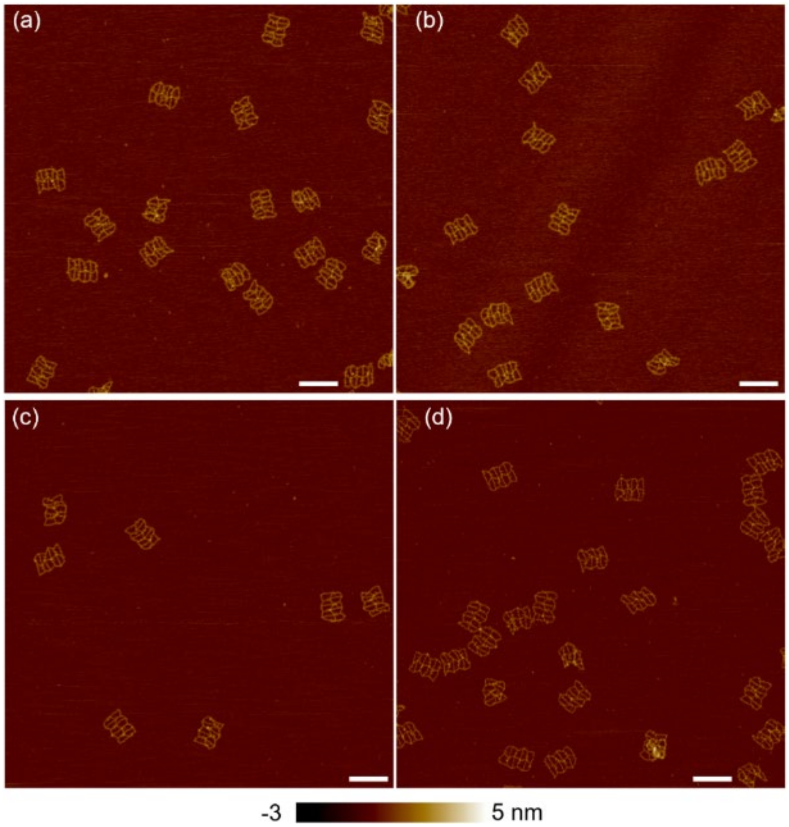


**Figure S30.** AFM images of the re-entrant honeycomb at 60°. Scale bar: 200 nm.

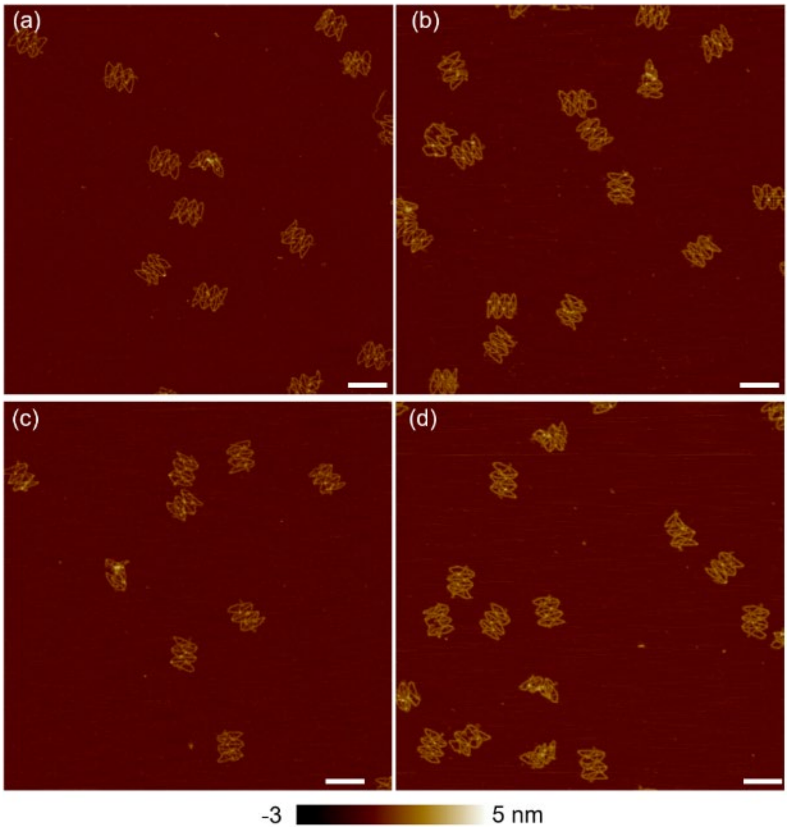

**Figure S31.** AFM images of the re-entrant honeycomb at 29°. Scale bar: 150 nm.

**Regular honeycomb**

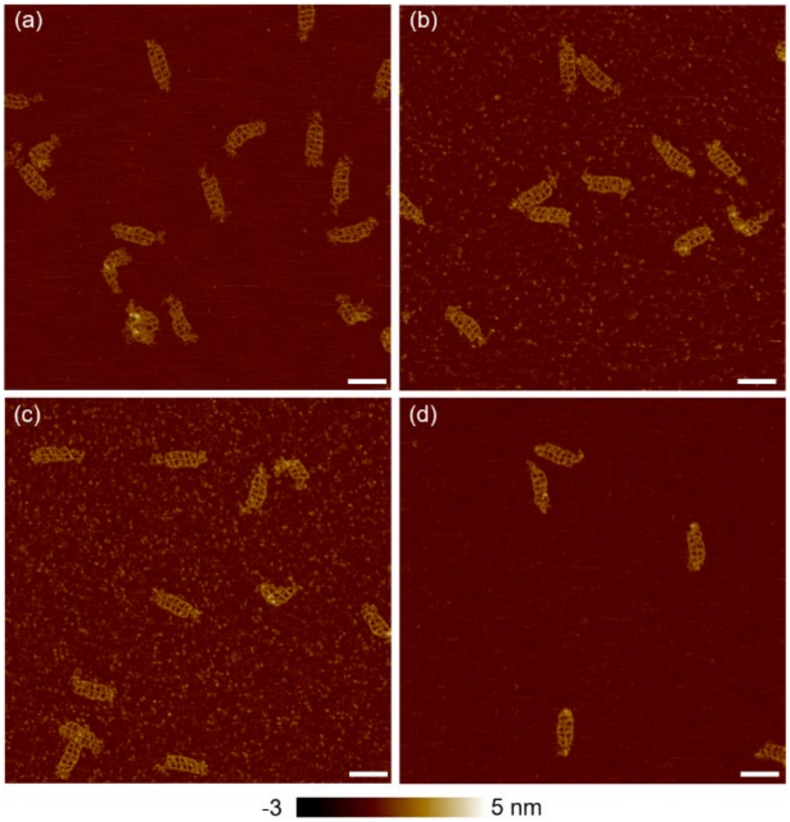


**Figure S32.** AFM images of the regular honeycomb at 90°. Scale bar: 150 nm.

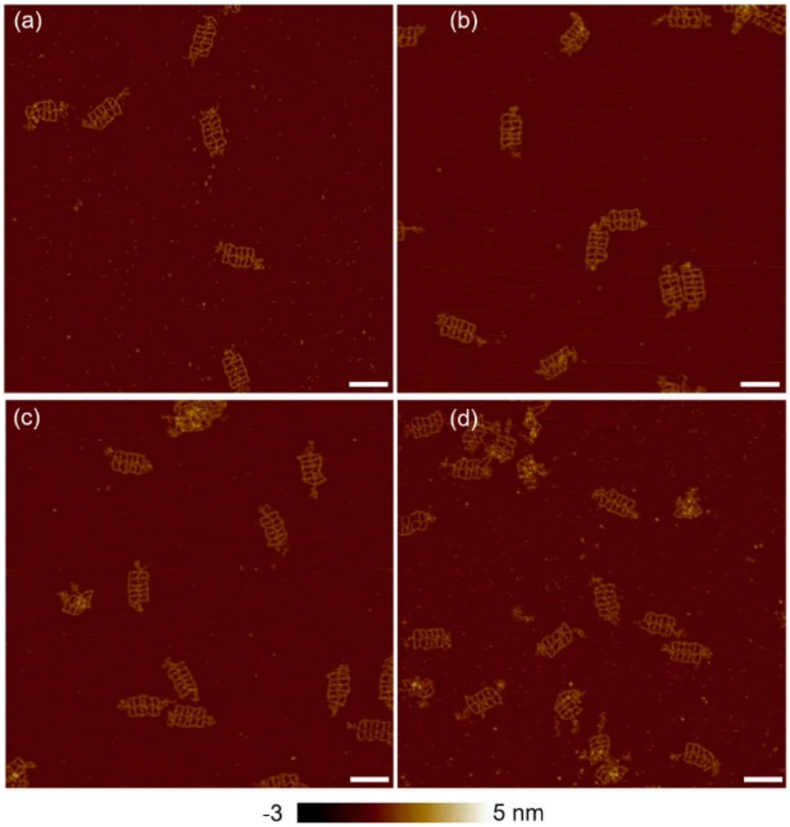


**Figure S33.** AFM images of the regular honeycomb at 120°. Scale bar: 150 nm.

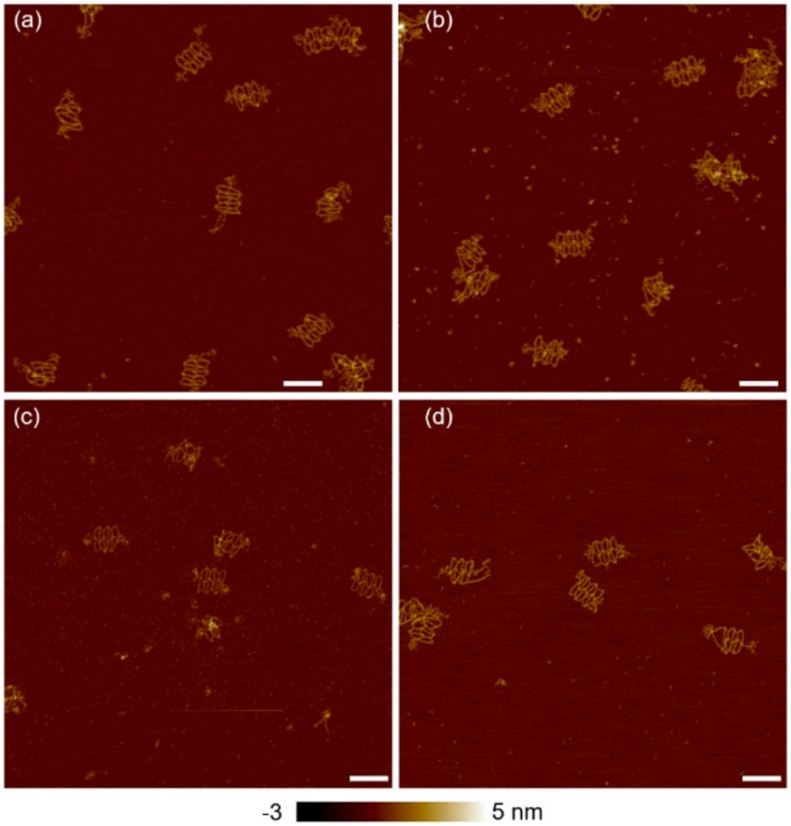

**Figure S34.** AFM images of the regular honeycomb at 146°. Scale bar: 150 nm.

**Re-entrant triangle**

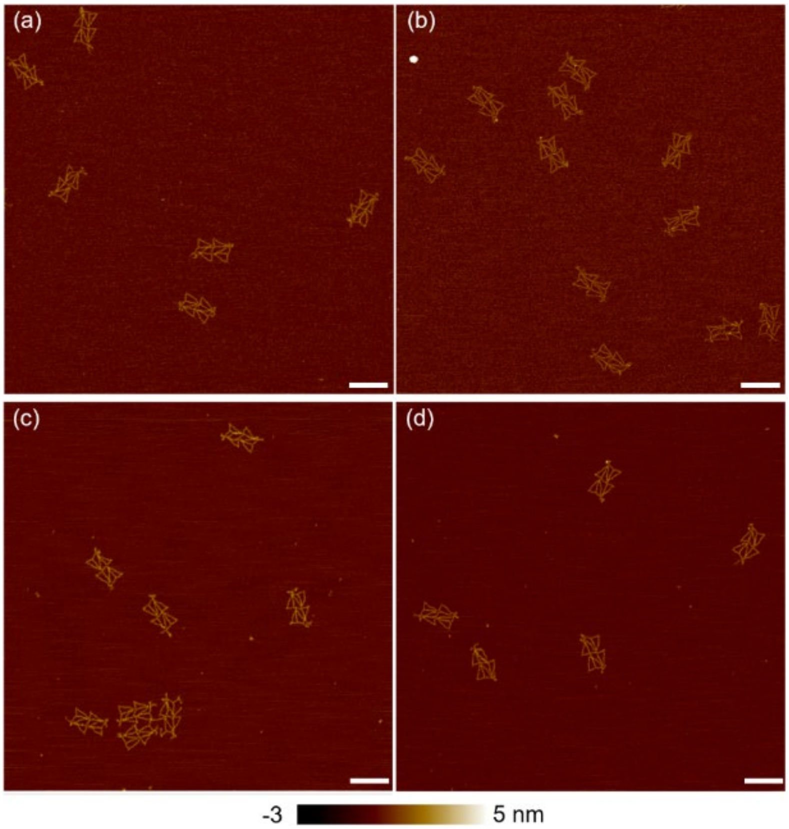


**Figure S35.** AFM images of the re-entrant triangle at 58°. (a)–(b) Static structures. (c)–(d) Dynamic structures reconfigured from the conformation at $\gamma = 30°$. Scale bar: 200 nm.

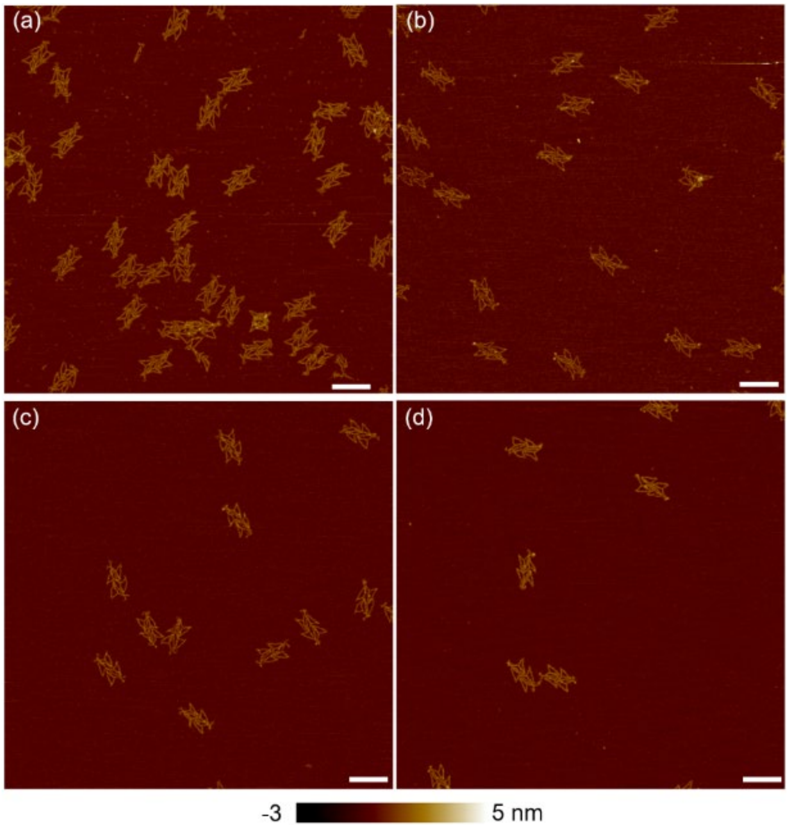


**Figure S36.** AFM images of the re-entrant triangle at 30°. (a)–(b) Static structures. (c)–(d) Dynamic structures reconfigured from the conformation at $\gamma$ = 58°. Scale bar: 200 nm.

**Rotating square**

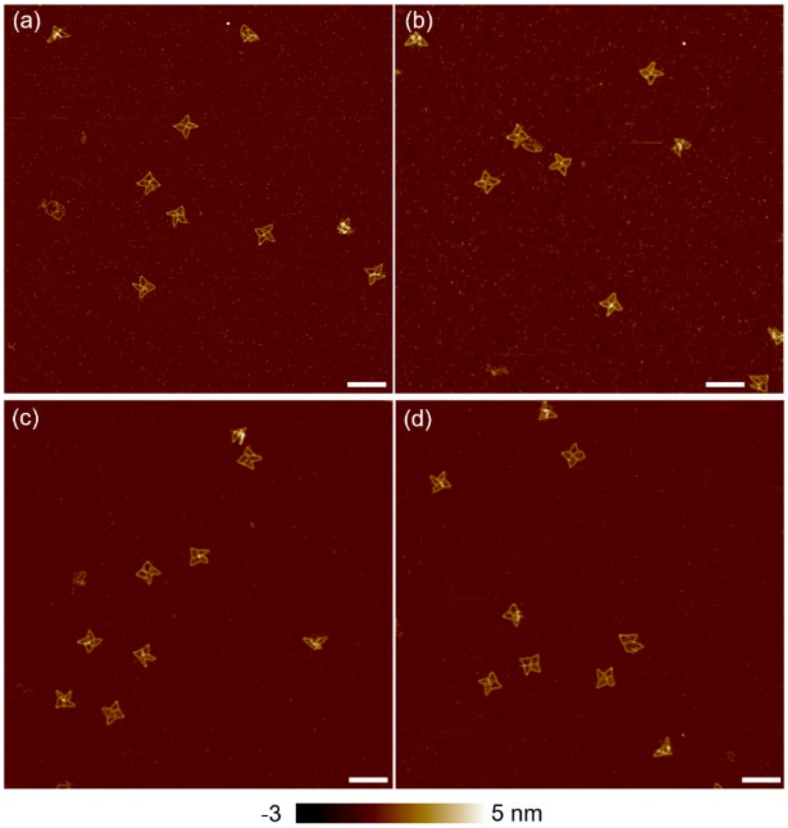


**Figure S37.** AFM images of the rotating square at 37°. Scale bar: 200 nm.

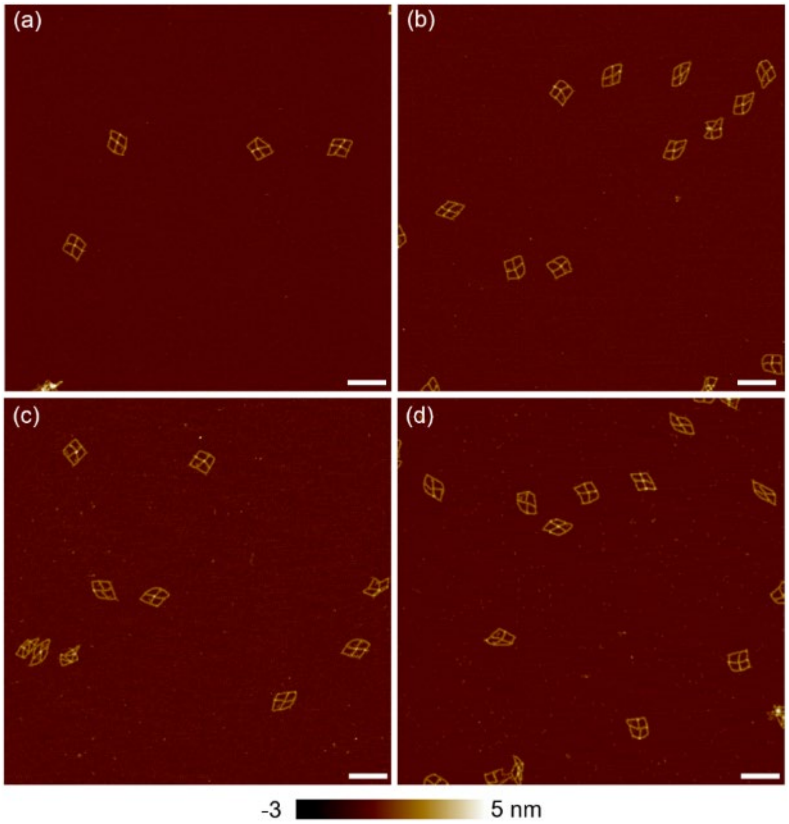


**Figure S38.** AFM images of the rotating square at 88°. Scale bar: 200 nm.

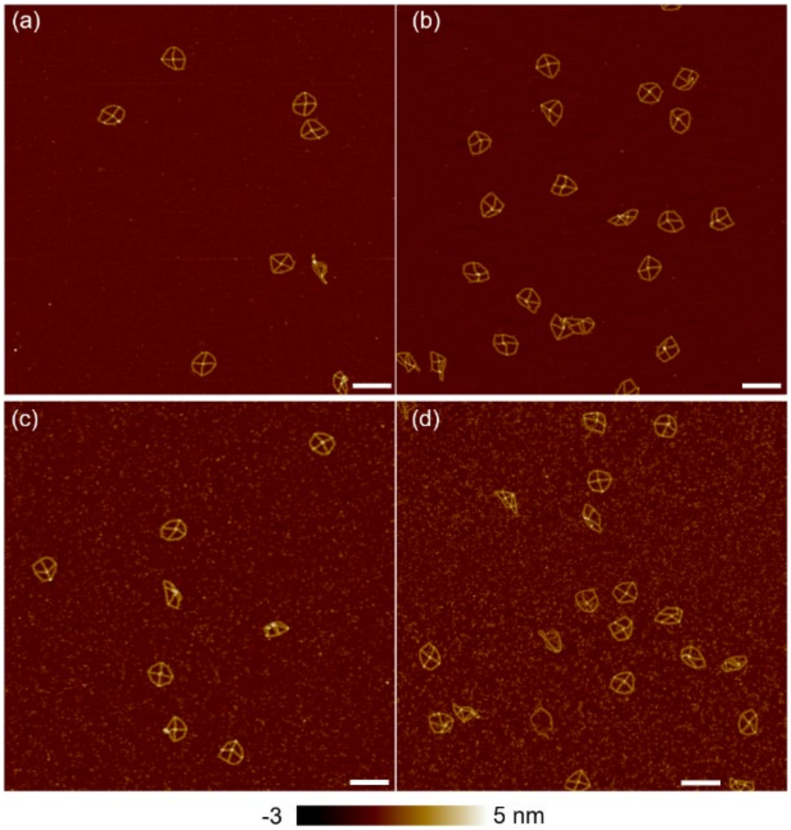


**Figure S39.** AFM images of the rotating square at 140°. Scale bar: 200 nm.

**Re-entrant triangle, control**

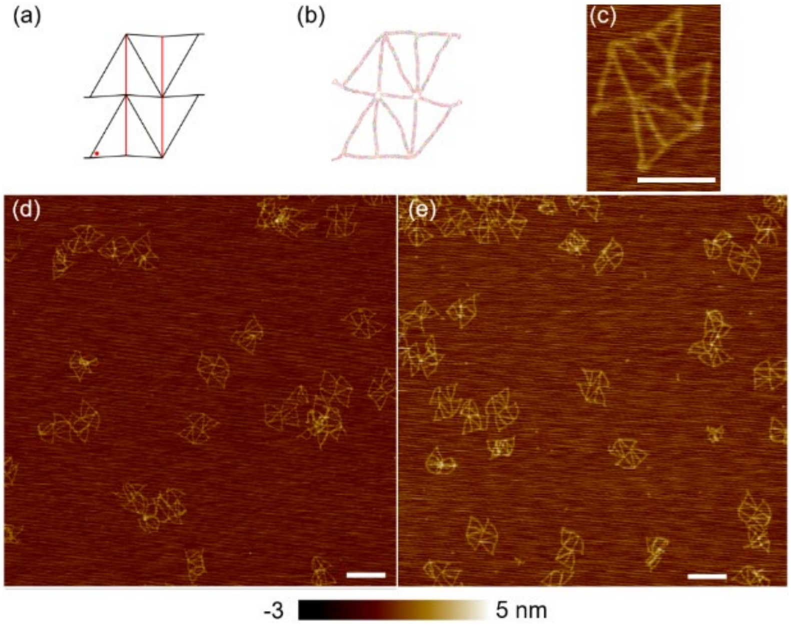


**Figure S40.** Re-entrant triangle at $\gamma = 56°$. (a) Schematic of the structure. (b) MD simulation result. The color scheme is the default in cogli1. (c) AFM image of an individual structure. Scale bar: 100 nm (d)–(e) AFM images at large scale. Scale bar: 200 nm.

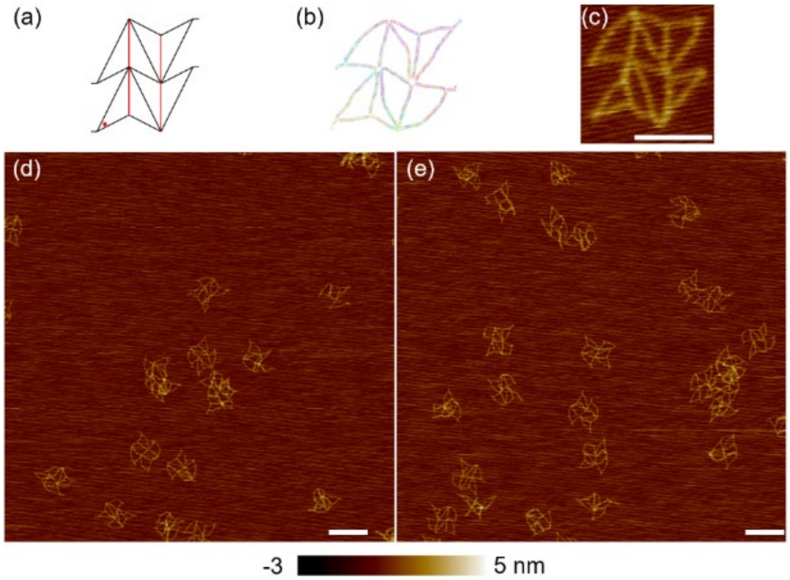

**Figure S41.** Re-entrant triangle at $\gamma = 36°$. (a) Schematic of the structure. (b) MD simulation result. (c) AFM image of an individual structure. Scale bar: 100 nm (d)–(e) AFM image at large scale. Scale bar: 200 nm.

**S7. Origami Sequence Information**
In this part, the sequences of all the DNA strands are listed. The name specifies the position of the strands, as illustrated in S3. These strands include staples and releasers.

**Re-entrant honeycomb**

| Re-entrant honeycomb staples | |
|---|---|
| Name | Sequence (5' to 3') |
| L[0][0] | GTCTGGAAGTTGACGACGACAGTATCGGCCTCAAAGTACGGT |
| L[0][1] | CAGTTTGAGGGTCATTCCATATAACAGTTGATGTGCATCTGC |
| L[0][2] | GCGAACGAGTAGTAGATGGGCGCATCGTAACCTCCCAATTCT |
| L[0][3] | GTCACGTTGGTGATTTAGTTTGACCATTAGATAATGGGATAG |
| L[0][4] | CCTGTAGCCAGTCTACTAATAGTAGTAGCATTGTCTGGCCTT |
| L[0][5] | GTGGCATCAATCTTTCATCAACATTAAATGTGAGCTGAAAAG |
| L[0][6] | AACCCGTCGGATATATTTTCATTTGGGGCGCGAGCGAGTAAC |
| L[0][7] | ACCTGTTTAGCTTCTCCGTGGGAACAAACGGCATGGTCAATA |
| Ls[0] | GGATTGACCGTACATTTCGCAA |
| L[1][0] | GGCCTCTTCGCGCGCCGCTACAGGGCGCGTACGATCGGTGCG |
| L[1][1] | CCGCGCTTAATTATTACGCCAGCTGGCGAAAGACCACACCCG |
| L[1][2] | TGCAAGGCGATGCGGTCACGCTGCGCGTAACCGGGGATGTGC |
| L[1][3] | TGGCAAGTGTATAAGTTGGGTAACGCCAGGGTGCTAGGGCGC |
| L[1][4] | TAAAGGGAGCCCCGGGTACCGAGCTCGAATTCATCGGAACCC |
| L[1][5] | CTAGAGGATCCCCCGATTTAGAGCTTGACGGGCAGGTCGACT |
| L[1][6] | GAACGTGGCGAAGTGCCAAGCTTGCATGCCTGGAAAGCCGGC |
| L[1][7] | AAACGACGGCCGAAAGGAAGGGAAGAAAGCGACGACGTTGTA |
| Ls[1] | AAGGAGCGGGCTTTCCCAGTCA |
| L[2][0] | CTGAGAAGTGTTGGAAGGGTTAGAACCTACCACGCCAGAATC |
| L[2][1] | CTTCTGAATAATTTTATAATCAGTGAGGCCACTTGGATTATA |
| L[2][2] | AGTCTGTCCATCATCAATATAATCCTGATTGTCGAGTAAAAG |
| L[2][3] | TGATGGCAATTCACGCAAATTAACCGTTGTAGGATTATCAGA |
| L[2][4] | TTATTAATTTTTCCAGAACAATATTACCGCCATGCCCGAACG |
| L[2][5] | TGCTGGTAATAAAAAGTTTGAGTAACATTATCCTATCGGCCT |
| L[2][6] | ACAAAGAAACCGCCTGAGTAGAAGAACTCAAAATTTTGCGGA |
| L[2][7] | TAACATCACTTACCAGAAGGAGCGGAATTATCTGATTAGTAA |
| Ls[2] | ATCATATTCCTCAATACTTCTT |
| L[3][0] | CGGATTCGCCTCTCATCGAGAACAAGCAAGCCGAAACAATAA |
| L[3][1] | CAAGTACCGCAGATTGCTTTGAATACCAAGTTGGTATTAAAC |
| L[3][2] | GCAGAGGCGAATTCCTTATCATTCCAAGAACGACAAAATCGC |
| L[3][3] | ATCGGCTGTCTTTATTCATTTCAATTACCTGACCAATCAATA |
| L[3][4] | ATGCAGAACGCTTTTAATGGAAACAGTACATATGTTCAGCTA |

| | |
|---|---|
| L[3][5] | TGAATTACCTTGCCTGTTTATCAACAATAGATCAATTTCATT |
| L[3][6] | CAAGAAAAATAAACAAAATTAATTACATTTAAAAGTCCTGAA |
| L[3][7] | AACATCAAGAAATATCCCATCCTAATTTACGATGATGAAACA |
| Ls[3] | GCATGTAGAAAGCAAAAGAAGA |
| L[4][0] | TTGCGGGAGGTTAGCCCCCTTATTAGCGTTTGACCTCCCGAC |
| L[4][1] | ATTTTCGGTCATTTGAAGCCTTAAATCAAGATTTTCATCGGC |
| L[4][2] | TTTGCACCCAGTAGCGTCAGACTGTAGCGCGTTAGTTGCTAT |
| L[4][3] | AAGTTTGCCTTCTACAATTTTATCCTGAATCTCGACAGAATC |
| L[4][4] | GCAAAATCACCATCCAAATAAGAAACGATTTTATTAGAGCCA |
| L[4][5] | TATTTATCCCAAGTAGCACCATTACCATTAGCACAGCCATAT |
| L[4][6] | ACGTCACCAATAATTTGCCAGTTACAAAATAAAAGGCCGGAA |
| L[4][7] | TTCCAGAGCCTGAAACCATCGATAGCAGCACCACGAGCGTCT |
| Ls[4] | GTAATCAGTAGTACCAACGCTA |
| L[5][0] | CACCAGAACCAGCCAAAAGGAATTACGAGGCATCAGAGCCGC |
| L[5][1] | AGATACATAACCCACCAGAGCCGCCGCCAGCACAACTAATGC |
| L[5][2] | GTTGAGGCAGGGAGATTTAGGAATACCACATTTTGACAGGAG |
| L[5][3] | ATTCATCAGTTTCAGACGATTGGCCTTGATATAGGTAGAAAG |
| L[5][4] | TTAAGAACTGGATACATGGCTTTTGATGATACTTATGCGATT |
| L[5][5] | CAGTAAGCGTCCTCATTATACCAGTCAGGACGTTTACCGTTC |
| L[5][6] | AAATCTACGTTATGGAAAGCGCAGTCTCTGAATTGGGAAGAA |
| L[5][7] | TTAAAGCCAGAAATAAAACGAACTAACGGAACTAAATCCTCA |
| Ls[5] | AACATTATTACTCACAAACAAA |
| L[6][0] | GGCTGACCTTCAGGAAACCGAGGAAACGCAATCGCATAGGCT |
| L[6][1] | AAGTTACCAGAATCAAGAGTAATCTTGACAAGTAGCCGAACA |
| L[6][2] | TCATTACCCAATTTTTAAGAAAAGTAAGCAGAAACCGGATAT |
| L[6][3] | TACCGAAGCCCATCAACGTAACAAAGCTGCTCATAGCTATCT |
| L[6][4] | GGGTAATTGAGCAACTTTAATCATTGTGAATTCAAAGTCAGA |
| L[6][5] | TGGTTTAATTTCGCTAATATCAGAGAGATAACGGGCTTGAGA |
| L[6][6] | TGAGTTAAGCCCACCAGAACGAGTAGTAAATTCCACAAGAAT |
| L[6][7] | CTGACGAGAAACAATAATAAGAGCAAGAAACAAAGGCTTGCC |
| Ls[6] | ATGAAATAGCAATTCAGTGAAT |
| L[7][0] | AGACACCACGGTAATTTCATCTTCTGACCTAAAGAAACGCAA |
| L[7][1] | TATATTTTAGTAATAAGTTTATTTTGTCACAACTTTTTCAAA |
| L[7][2] | ATTCATATGGTAAGACAAAGAACGCGAGAAAATCAATAGAAA |
| L[7][3] | AATCCAATCGCTTACCAGCGCCAAAGACAAAATGCTGATGCA |
| L[7][4] | CAATAGTGAATGTCACCGACTTGAGCCATTTGTGAGAAGAGT |
| L[7][5] | GAATTATCACCTTATCAAAATCATAGGTCTGACATTAAAGGT |
| L[7][6] | TTTTAACCTCCTAAATATTGACGGAAATTATTGAGACTACCT |

| | |
|---|---|
| L[7][7] | GGGAGGGAAGGGGCTTAGGTTGGGTTATATAAAACCGATTGA |
| Ls[7] | CTATATGTAAAGGGCGACATTC |
| L[8][0] | ATTCTTACCAGCACCAGCAGAAGATAAAACAGCGTTATACAA |
| L[8][1] | ACCGAACGAACTATAAAGCCAACGCTCAACAGCATTAAAAAT |
| L[8][2] | TTGAGAATCGCCTGATAGCCCTAAAACATCGCTAGGGCTTAA |
| L[8][3] | TAATGCGCGAACATATTTAACAACGCCAACATTATTAGTCTT |
| L[8][4] | AAAGGGACATTTCCAGACGACGACAATAAACAACCAGTAATA |
| L[8][5] | AAGTAATTCTGCTGGCCAACAGAGATAGAACCACAAAAGGTA |
| L[8][6] | GAAAGCGTAAGATAAGAGAATATAAAGTACCGCTTCTGACCT |
| L[8][7] | TCGAGCCAGTAAATACGTGGCACAGACAATATAGAGGCATTT |
| Ls[8] | TTTTGAATGGCGTAATTTAGGC |
| L[9][0] | TCAATATCTGGCAACAGCTGATTGCCCTTCACAAACCCTCAA |
| L[9][1] | AGTGAGACGGGTCAGTTGGCAAATCAACAGTTTCTTTTCACC |
| L[9][2] | GAGGAAGGTTAATTGGGCGCCAGGGTGGTTTTGAAAGGAATT |
| L[9][3] | GCGGTTTGCGTTCTAAAATATCTTTAGGAGCAGCGGGGAGAG |
| L[9][4] | ACTCACATTAATTCGACAACTCGTATTAAATCGAGTGAGCTA |
| L[9][5] | TTTACAAACAATTGCGTTGCGCTCACTGCCCGAGTATTAGAC |
| L[9][6] | GGGAAACCTGTTAATACATTTGAGGATTTAGACTTTCCAGTC |
| L[9][7] | CCGTCAATAGACGTGCCAGCTGCATTAATGAATAGATTAGAG |
| Ls[9] | TCGGCCAACGCCTAACAACTAA |
| L[10][0] | GAATAGCCCGAATTCAACCGTTCTAGCTGATATAAATCAAAA |
| L[10][1] | ATCAATATGATGATAGGGTTGAGTGTTGTTCCTCAAATCACC |
| L[10][2] | AAGAGTCCACTGGTGAGAAAGGCCGGAGACAGAGTTTGGAAC |
| L[10][3] | AGATTCAAAAGATTAAAGAACGTGGACTCCAAGTGTAGGTAA |
| L[10][4] | ACTTTTGCGGGGGTCGAGGTGCCGTAAAGCACACCCTGTAAT |
| L[10][5] | AAGTTTTTTGGAGAAGCCTTTATTTCAACGCACACCCAAATC |
| L[10][6] | TTTTTAGAACCATGGCCCACTACGTGAACCATAGGATAAAAA |
| L[10][7] | CTATCAGGGCGCTCATATATTTTAAATGCAATGAAAAACCGT |
| Ls[10] | GCCTGAGTAATCGTCAAAGGGC |
| L[11][0] | GCATGTCAATCGGTGAATTTCTTAAACAGCTTCGTAAAACTA |
| L[11][1] | CTTGCTTTCGAATATGTACCCCGGTTGATAATGGTTTATCAG |
| L[11][2] | CCAAAAACAGGAAAGGAGCCTTTAATTGTATCCAGAAAAGCC |
| L[11][3] | AAAAGGCTCCAAAGATTGTATAAGCAAATATTATCTCCAAAA |
| L[11][4] | TAAACAACTTTGGAACGCCATCAAAAATAATTGGGATTTTGC |
| L[11][5] | TTTAACCAATACAACAGTTTCAGCGGAGTGAGCAGCTCATTT |
| L[11][6] | AACAACTAAAGCGCATTAAATTTTTGTTAAATAATAGAAAGG |
| L[11][7] | TTGTTAAAATTGAATTGCGAATAATAATTTTTCGTTAATATT |
| Ls[11] | TCACGTTGAAATAAATTGTAAA |

| | |
|---|---|
| S[0][0] | GCCAGCTTTCCTAGCTCAACATGTTTTAAATATCGCACTCCA |
| S[0][1] | TAAGAGGTCATCCATTCAGGCTGCGCAACTGTCTCCTTTTGA |
| S[0][2] | AGCGCCATTCGTTTTGCGGATGGCTTAGAGCTAACCAGGCAA |
| S[0][3] | ATATAATGCTGGGCACCGCTTCTGGTGCCGGATAATTGCTGA |
| S[1][0] | CGTATAACGTGCAGGTCAGGATTAGAGAGTACTTGACGAGCA |
| S[1][1] | TATCGCGTTTTATTAAAGGGATTTTAGACAGGAGACTTCAAA |
| S[1][2] | ACAGGAGGCCGAATTCGAGCTTCAAAGCGAACCGGGAGCTAA |
| S[1][3] | GCAAACTCCAACTTTCCTCGTTAGAATCAGAGCAGACCGGAA |
| S[2][0] | CGTAAAACAGATGCATCAAAAAGATTAAGAGGATTATTTGCA |
| S[2][1] | AAATCAAAAATATATACAGTAACAGTACCTTTGAATGACCAT |
| S[2][2] | ACGTCAGATGACAGGTCTTTACCCTGACTATTTTCAGGTTTA |
| S[2][3] | GCAAAGCGGATAATAAAGAAATTGCGTAGATTATAGTCAGAA |
| S[3][0] | AATCATTACCGCTCAAATGCTTTAAACAGTTCTCATCGTAGG |
| S[3][1] | GACTGGATAGCATTCTAAGAACGCGAGGCGTTAAAATGTTTA |
| S[3][2] | GCTTATCCGGTGTCCAATACTGCGGAATCGTCGATATAGAAG |
| S[3][3] | ATTGAATCCCCCGCCCAATAGCAAGCAAATCAATAAATATTC |
| S[4][0] | AAATCACCGGAAAAAGAAGTTTTGCCAGAGGGTTCATAATCA |
| S[4][1] | TAACCCTCGTTCTCAGAACCGCCACCCTCAGAAACACTATCA |
| S[4][2] | GAGCCGCCACCTACCAGACGACGATAAAAACCGCCTCCCTCA |
| S[4][3] | GAGGCTTTTGCACCAGAGCCACCACCGGAACCAAAATAGCGA |
| S[5][0] | CAGGCAAGGCATTCCAGACGTTAGTAAATGAAATAAATCATA |
| S[5][1] | CATTCCACAGACTAAATCGGTTGTACCAAAAAACGCCTGTAG |
| S[5][2] | AGAGCATAAAGCAGCCCTCATAGTTAGCGTAAATAAAGCCTC |
| S[5][3] | TTTTGTCGTCTAAGAATTAGCAAAATTAAGCACGATCTAAAG |
| S[6][0] | TTCCTGTGTGACACTGAGTTTCGTCACCAGTATCATAGCTGT |
| S[6][1] | CACCACCCTCAAGCATAAAGTGTAAAGCCTGGCCCTCAGAGC |
| S[6][2] | TACGAGCCGGATTTTCAGGGATAGCAAGCCCACCACACAACA |
| S[6][3] | ATGTACCGTAAAATTGTTATCCGCTCACAATTATAGGAACCC |
| S[7][0] | AACGCTCATGGCCCTCAGAACCGCCACCCTCACAACAGGAAA |
| S[7][1] | AAGTATAGCCCTTACATTGGCAGATTCACCAGGGGTTGATAT |
| S[7][2] | AAATGGATTATGGAATAGGTGTATCACCGTACCAATCGTCTG |
| S[7][3] | TAGTACCGCCAAAATACCTACATTTTGACGCTTCAGGAGGTT |
| S[8][0] | TAACCTTGCTTTCAGTACCAGGCGGATAAGTGTGTGAGTGAA |
| S[8][1] | TATTAAGAGGCGAAAACATAGCGATAGCTTAGAACATGAAAG |
| S[8][2] | TTAGAATCCTTTGAGACTCCTCAAGAGAAGGATAATTTTCCC |
| S[8][3] | CGGGGTTTTGCCTGTAAATCGTCGCTATTAATTTAGGATTAG |
| S[9][0] | TGAAAATAGCACCCTGCCTATTTCGGAACCTAACGTCAAAAA |
| S[9][1] | TTTTAACGGGGGACGGGAGAATTAACTGAACATGGTAATAAG |

| | |
|---|---|
| S[9][2] | GAAGCGCATTATCAGTGCCTTGAGTAACAGTGTAAAAACAGG |
| S[9][3] | CAGTTAATGCCGCCTTTACAGAGAGAATAACACCCGTATAAA |
| S[10][0] | CAATGACAACAGGAGCAAACAAGAGAATCGATGTTGCGCCGA |
| S[10][1] | GTAGCTATTTTTGCAGGGAGTTAAAGGCCGCTGCCGGAGAGG |
| S[10][2] | TCGCTGAGGCTTGAGAGATCTACAAAGGCTATATATATTCGG |
| S[10][3] | CCTGAGAGTCTACCATCGCCCACGCATAACCGCAGGTCATTG |
| S[11][0] | CAGCAGCGAAAGTGGTTCCGAAATCGGCAAAATCGTCACCCT |
| S[11][1] | GCAGCAAGCGGAGGACTAAAGACTTTTTCATGTGAGAGAGTT |
| S[11][2] | CAGAGGCTTTGTCCACGCTGGTTTGCCCCAGCGCAACGGCTA |
| S[11][3] | CCTGTTTGATGGACAGCATCGGAACGAGGGTAAGGCGAAAAT |
| S[12][0] | GGTAAAATACGTAAAGCATCACCTTGCTGAACCCATTAAACG |
| S[12][1] | TTAACACCGCCAAGAATACACTAAAACACTCAGCGGTCAGTA |
| S[12][2] | GAAAGAGGCAATGCAACAGTGCCACGCTGAGAAACCTAAAAC |
| S[12][3] | AATGAAAAATCTAATGCCACTACGAAGGCACCGCCAGCAGCA |
| S[13][0] | ATACCAAGCGCCTAGAAAAAGCCTGTTTAGTACCCAGCGATT |
| S[13][1] | GACCGTGTGATTGTGTCGAAATCCGCGACCTGTTGAAATACC |
| S[13][2] | GCCTGATAAATAAATAAGGCGTTAAATAAGAATTGTATCATC |
| S[13][3] | AATCATAATTAGAAACAAAGTACAACGGAGATTAAACACCGG |
| S[14][0] | AACGAGGCGCAAAATACATACATAAAGGTGGCACTTAGCCGG |
| S[14][1] | AAGAACTGGCAAGGACAGATGAACGGTGTACAGAATACCCAA |
| S[14][2] | AACTTTGAAAGTGATTAAGACTCCTTATTACGCGAACTGACC |
| S[14][3] | GCAAACGTAGAGACGGTCAATCATAAGGGAACCAGTATGTTA |
| mL[0][0] | AACATTAAATGTCGCATCGTAACCTCCCAATT |
| mL[0][1] | CTGCGAACGAGTGGTGGCATCAATCTTTCATC |
| mL[2][0] | GAGTAACATTATAATCCTGATTGTCGAGTAAA |
| mL[2][1] | AGAGTCTGTCCATTGCTGGTAATAAAAAGTTT |
| mL[4][0] | CATTACCATTAGACTGTAGCGCGTTAGTTGCT |
| mL[4][1] | ATTTTGCACCCATTATTTATCCCAAGTAGCAC |
| mL[6][0] | TCAGAGAGATAAAAAGTAAGCAGAAACCGGAT |
| mL[6][1] | ATTCATTACCCAATGGTTTAATTTCGCTAATA |
| mL[8][0] | CAGAGATAGAACCTAAAACATCGCTAGGGCTT |
| mL[8][1] | AATTGAGAATCGAAAGTAATTCTGCTGGCCAA |
| mL[10][0] | TTATTTCAACGCGGCCGGAGACAGAGTTTGGA |
| mL[10][1] | ACAAGAGTCCACCAAGTTTTTTGGAGAAGCCT |
| sL[0][0] | TTCCTGTAGCCAAGTATCGGCCTCAAAGTACG |
| sL[0][1] | GTGTCTGGAAGTAGTAGTAGCATTGTCTGGCC |
| sL[2][0] | CGTTATTAATTTTAGAACCTACCACGCCAGAA |
| sL[2][1] | TCCTGAGAAGTGATATTACCGCCATGCCCGAA |

| | |
|---|---|
| sL[4][0] | CAGCAAAATCACTATTAGCGTTTGACCTCCCG |
| sL[4][1] | ACTTGCGGGAGGAGAAACGATTTTATTAGAGC |
| sL[6][0] | GAGGGTAATTGAAGGAAACGCAATCGCATAGG |
| sL[6][1] | CTGGCTGACCTTTCATTGTGAATTCAAAGTCA |
| sL[8][0] | TAAAAGGGACATAAGATAAAACAGCGTTATAC |
| sL[8][1] | AAATTCTTACCACGACAATAAACAACCAGTAA |
| sL[10][0] | ATACTTTTGCGGTTCTAGCTGATATAAATCAA |
| sL[10][1] | AAGAATAGCCCGGCCGTAAAGCACACCCTGTA |

**Regular honeycomb**

| Regular honeycomb staples | |
|---|---|
| Name | Sequence (5' to 3') |
| L[0][0] | CCCCCGATTTATCTGGAAGTTTTGGGGCGCGCTAAAGGGAG |
| L[0][1] | GTGGCATCAATTAAAGAACGTAAATCGGAACCAGCTGAAAAG |
| L[1][0] | GTGTAGCGGTCGAAAGACTTCAAATATCGCGTGCGCTGGCAA |
| L[1][1] | AGAGGAAGCCCACGCTGCGCGTAACCACCACAAAAAAGATTA |
| L[1][2] | TTAATGCGCCGAGAAGCAAAGCGGATTGCATCCCCGCCGCGC |
| L[2][0] | TAAAATATCTTGACCAGGCGCACGAGAAACACGAAGGTTATC |
| L[2][1] | AGTAAATTGGGCAACAGTGCCAAGGAATTGAGCAGAACGAGT |
| L[3][0] | ATTCGACAACTCAACCTAAAACGAAAGAGGCATTTACAAACA |
| L[3][1] | TACGAAGGCACCGTATTAAATCCTTTGCCCGAGTAATGCCAC |
| L[3][2] | TTTTAAAAGTTTCCATTAAACGGGTAAAATACACGTTATTAA |
| L[4][0] | GTGTGATAAATTCAGGAGGTTACTGAGTTTCGAATACCGACC |
| L[4][1] | AAACTACAACGTAGGTTGGGTTAATGGTTTGATCACCAGTAC |
| L[5][0] | TACCAGTATAAAACATGAAAGTATTAAGAGGCTACAAATTCT |
| L[5][1] | TATTATTCTGAAGCCAACGCTCAACAGTAGGGTTTCGGAACC |
| L[5][2] | AATCGCCATATACAGTTAATGCCCCCTGCCTACTTAATTGAG |
| L[6][0] | ACCAAGTACCGATCATTACCGTCAAGATTAGTCGGGTATTAA |
| L[6][1] | CACCCAGCTACAATGCAGAACCATTCCAAGAATGCTATTTTG |
| L[7][0] | TTCCCTTAGAAAATAATTTTTTCACGTTGAAATTAATTAATT |
| L[7][1] | GAATTGCGAATTCCTTGAAAACATAGCGATAGACAACTAAAG |
| L[7][2] | GACGCTGAGAAGCGGAGTGAGAATAGAAAGGACTTAGATTAA |
| L[8][0] | TAACGGATTCGTCTTAAACAGATCGTCACCCTGGAGAAACAA |
| L[8][1] | AGACAGCATCGTGAATAATGGCTTTTACATCGCAGCAGCGAA |
| L[9][0] | TATTAGTCTTTAGGAATACCACATTCAACTAATTTGAATGGC |
| L[9][1] | AGTTGAGATTTAATGCGCGAACTGATAGCCCTAAGATTCATC |
| L[9][2] | CATTAAAAATAAACAACATTATTACAGGTAGAAAAACATCGC |
| L[10][0] | TAGCAATACTTGACGACGATAATTCATTGAATTTAACCGTTG |
| L[10][1] | TGCTTTAAACATCAGAGCGGGCATCACGCAAACCCCCTCAAA |
| L[11][0] | TGCCCTTCACCAACGCGCGGGGAGAGGCGGTTAACAGCTGAT |
| L[11][1] | ATGAATCGGCCGCCTGGCCCTGAGAGAGTTGCAGCTGCATTA |
| L[11][2] | CCACGCTGGTTAGTCGGGAAACCTGTCGTGCCAGCAAGCGGT |
| mL[0][2] | GATTTAGTTTGATCAGGGCGAGTCGAGGTGCCCGAACGAGTA |
| mL[0][3] | AAAAACCGTCTACCATTAGATTTTGGGGCGCGTCAAAGGGCG |
| mL[2][2] | CGGATATTCATAAGCATCACCAGTTGGCAAATTGACAAGAAC |
| mL[2][3] | TGAAAAATCTATACCCAAATCACGAGAAACACCAGCAGCAAA |
| mL[4][2] | CCTCAGAGCCACAATCGCAAGTTTCATCTTCTGAACCGCCAC |

| | |
|---|---|
| mL[4][3] | TGATGCAAATCCCACCCTCATACTGAGTTTCGATGTAAATGC |
| mL[6][2] | ATTCTAAGAACACAAGAAAAATAATCGGCTGTCTTATCCGGT |
| mL[6][3] | ATAAGTCCTGAGCGAGGCGTTTCAAGATTAGTTCAACAATAG |
| mL[8][2] | CCACGCATAACGCACGTAAAAATGAATATACACAACCATCGC |
| mL[8][3] | CAAAATTATTTCGATATATTCATCGTCACCCTCCTACCATAT |
| mL[10][2] | AGAGGGGGTAAGACAGGAACGCACCGAGTAAAAAGTTTTGCC |
| mL[10][3] | AAGGGATTTTATAGTAAAATGATTCATTGAATAGGCCGATTA |
| fL[0][0] | TCATTCCATATCTAAAGGGAGCCCCCGATTTATCTGGAAGTT |
| fL[0][1] | AAATCGGAACCAACAGTTGATTCCCAATTCTGGTAAAGCACT |
| fL[0][2] | GATTTAGTTTGGTTTTTTGGGGTCGAGGTGCCCGAACGAGTA |
| fL[0][3] | ACCCAAATCAAACCATTAGATACATTTCGCAAGTGAACCATC |
| fL[0][4] | ACCTGTTTAGCATCAGGGCGATGGCCCACTACATGGTCAATA |
| fL[0][5] | AAAAACCGTCTTATATTTTCATTTGGGGCGCGTCAAAGGGCG |
| fL[0][6] | GTGGCATCAATTAAAGAACGTGGACTCCAACGAGCTGAAAAG |
| fL[0][7] | GAGTCCACTATTCTACTAATAGTAGTAGCATTTTTGGAACAA |
| fL[2][0] | ATAGGCTGGCTGAAGGTTATCTAAAATATCTTGACCAGGCGC |
| fL[2][1] | AAGGAATTGAGGACCTTCATCAAGAGTAATCTCAACAGTTGA |
| fL[2][2] | CGGATATTCATATATCTGGTCAGTTGGCAAATTGACAAGAAC |
| fL[2][3] | ACCCTCAATCATACCCAAATCAACGTAACAAACAAATATCAA |
| fL[2][4] | CAGTGAATAAGAAGCATCACCTTGCTGAACCTGCTGCTCATT |
| fL[2][5] | TGAAAAATCTAGCTTGCCCTGACGAGAAACACCAGCAGCAAA |
| fL[2][6] | AGTAAATTGGGCAACAGTGCCACGCTGAGAGCCAGAACGAGT |
| fL[2][7] | AACACCGCCTGCTTGAGATGGTTTAATTTCAAGGTCAGTATT |
| fL[4][0] | TAGTACCGCCAAATACCGACCGTGTGATAAATTCAGGAGGTT |
| fL[4][1] | TAATGGTTTGACCCTCAGAACCGCCACCCTCAGACCTAAATT |
| fL[4][2] | CCTCAGAGCCATTTTAGTTAATTTCATCTTCTGAACCGCCAC |
| fL[4][3] | TTTCAAATATACCACCCTCATTTTCAGGGATAGAGAAAACTT |
| fL[4][4] | TAGGAACCCATCAATCGCAAGACAAAGAACGCGCAAGCCCAA |
| fL[4][5] | TGATGCAAATCGTACCGTAACACTGAGTTTCGATGTAAATGC |
| fL[4][6] | AAACTACAACGTAGGTTGGGTTATATAACTATTCACCAGTAC |
| fL[4][7] | AACCTCCGGCTCCTGTAGCATTCCACAGACAGCTACCTTTTT |
| fL[6][0] | CGCCCAATAGCCGGGTATTAAACCAAGTACCGATCATTACCG |
| fL[6][1] | CATTCCAAGAAAAGCAAATCAGATATAGAAGGCTTTCCTTAT |
| fL[6][2] | ATTCTAAGAACAACCAATCAATAATCGGCTGTCTTATCCGGT |
| fL[6][3] | GAGCATGTAGAGCGAGGCGTTTTAGCGAACCTCCTAATTTAC |
| fL[6][4] | GGGAGGTTTTGACAAGAAAAATAATATCCCATCCCGACTTGC |
| fL[6][5] | ATAAGTCCTGAAAGCCTTAAATCAAGATTAGTTCAACAATAG |
| fL[6][6] | CACCCAGCTACAATGCAGAACGCGCCTGTTTATGCTATTTTG |

| | |
|---|---|
| fL[6][7] | CATGTTCAGCTAATTTTATCCTGAATCTTACCCAATAAACAA |
| fL[8][0] | CTTGATACCGAGGAGAAACAATAACGGATTCGTCTTAAACAG |
| fL[8][1] | CTTTTACATCGTAGTTGCGCCGACAATGACAAGTAACAGTAC |
| fL[8][2] | CCACGCATAACTTAACGTCAGATGAATATACACAACCATCGC |
| fL[8][3] | GATTTTCAGGTCGATATATTCGGTCGCTGAGGAAATTGCGTA |
| fL[8][4] | GTTAAAGGCCGGCACGTAAAACAGAAATAAAGCTTGCAGGGA |
| fL[8][5] | CAAAATTATTTCTTTTGCGGGATCGTCACCCTCCTACCATAT |
| fL[8][6] | AGACAGCATCGTGAATAATGGAAGGGTTAGAACAGCAGCGAA |
| fL[8][7] | GATTATACTTCGAACGAGGGTAGCAACGGCTATGATTGTTTG |
| fL[10][0] | AAAACCAAAATTTAACCGTTGTAGCAATACTTGACGACGATA |
| fL[10][1] | CATCACGCAAAAGCGAGAGGCTTTTGCAAAAGAGAGTCTGTC |
| fL[10][2] | AGAGGGGGTAATCAGTGAGGCCACCGAGTAAAAAGTTTTGCC |
| fL[10][3] | TGTTTTTATAATAGTAAAATGTTTAGACTGGATCCTGAGAAG |
| fL[10][4] | TACTGCGGAATGACAGGAACGGTACGCCAGAATAGCGTCCAA |
| fL[10][5] | AAGGGATTTTACGTCATAAATATTCATTGAATAGGCCGATTA |
| fL[10][6] | TGCTTTAAACATCAGAGCGGGAGCTAAACAGGCCCCCTCAAA |
| fL[10][7] | CCTCGTTAGAAGTTCAGAAAACGAGAATGACCAACGTGCTTT |
| S[0][0] | GAAAGCCGGCGCATGTTTTAAATATGCAACTAGCTTGACGGG |
| S[0][1] | CTGTAGCTCAAAACGTGGCGAGAAAGGAAGGGGAATATAATG |
| S[0][2] | AAGGAGCGGGCATGGCTTAGAGCTTAATTGCTAAGAAAGCGA |
| Sb[1][0] | AAGCGAACCAGTTTTGATAAGAGGTCATTTTTTCGAGCTTCA |
| Sm[1][0] | GAAAGAGGGATTAGAGAGTACCTTAACTCCAA |
| Sm[1][1] | CAGGTCAGACAGATGAACGGTGTACCAACTTT |
| Ss[1] | TAATTGCTCCACCGGAAGCA |
| S[2][0] | CAACTAATAGAACGGTCAATCATAAGGGAACCGGAGCACTAA |
| S[2][1] | ACGAGGCGCAGTTAGAGCCGTCAATAGATAATCTTAGCCGGA |
| S[2][2] | ATTTAGAAGTATCCGCGACCTGCTCCATGTTAACATTTGAGG |
| Sb[3][0] | CACTCATCTTTCATCGCCTGATAAATTGTGTCTACACTAAAA |
| Sm[3][0] | AGCCCGGAAAACAAAGTACAACGGGATTATAC |
| Sm[3][1] | CAAGCGCGATAGGTGTATCACCGTATAAGTAT |
| Ss[3] | AGATTTGTATGACCCCCAGC |
| S[4][0] | AAGAATAAACAGCGGATAAGTGCCGTCGAGAGGGCGTTAAAT |
| S[4][1] | CTCAGTACCAGCCGGAATCATAATTACTAGAACGGGGTTTTG |
| S[4][2] | TAGTATCATATCAAGAGAAGGATTAGGATTAGAAAGCCTGTT |
| Sb[5][0] | AATCCTGTTTGGATAGGGTTGAGTGTTGTTCCAGCAGGCGAA |
| Sm[5][0] | CTATGGTTCCTTATAAATCAAAAGCGAAATCG |
| Sm[5][1] | GCAAAATCGCTTTGACGAGCACGTGGCGCGTA |
| Ss[5] | AATAGCCCGAATGGTGGTTC |

| | |
|---|---|
| Sb[6][0] | TGAATTACCTTATCTACGTTAATAAAACGAACTTAATCATTG |
| Sm[6][0] | AATCAGGTTACCAGTCAGGACGTTAAGAACTG |
| Sm[6][1] | GCTCATTACTTTACCCTGACTATTAAATCAAA |
| Ss[6] | GGGAAGAAAAATGCGATTTT |
| Sb[7][0] | TTTGCGGAACAATGGCAATTCATCAATATAATACATTATCAT |
| Sm[7][0] | AGCAGAAGATCATCATATTCCTGACAGAAGGA |
| Sm[7][1] | GCGGAATTATAAAACAGAGGTGAGGAACCACC |
| Ss[7] | TTATCAGATGAAGAAACCAC |
| Sb[8][0] | GCGTAACGATCTTTGCTAAACAACTTTCAACACTCATAGTTA |
| Sm[8][0] | GAGGACTATAGTAAATGAATTTTCTCGTCTTT |
| Sm[8][1] | CCAGACGTAAGACTTTTTCATGAGGAGGCTTT |
| Ss[8] | TGTATGGGATTAAAGTTTTG |
| Sb[9][0] | GTAATTTAGGCAAGTAATTCTGTCCAGACGACACGCCAACAT |
| Sm[9][0] | TTTATCAAAGAATATAAAGTACCGTCGAGCCA |
| Sm[9][1] | GTAATAAGAATCATAGGTCTGAGAATAGTGAA |
| Ss[9] | ACAAAAGGTAAGAGGCATTT |
| Sb[10][0] | ATAACATCACTATATTACCGCCAGCCATTGCATTGATTAGTA |
| Sm[10][0] | GTGGTTTTCGGCCTTGCTGGTAATGAAGAACT |
| Sm[10][1] | CAAACTATTCTTTTCACCAGTGAGGCGCCAGG |
| Ss[10] | ATCCAGAACATGCCTGAGTA |
| S[11][0] | CAAAAGGAATTGAAATGGATTATTTACATTGGTACATAACGC |
| S[11][1] | CTCAATCGTCTACGAGGCATAGTAAGAGCAACCATTTTGACG |
| S[11][2] | CCCTCGTTTACAAACGCTCATGGAAATACCTAACTATCATAA |
| Sb[12][0] | ACACGACCAGTGTAAGAATACGTGGCACAGACTTCACCAGTC |
| Sm[12][0] | TTGAATACAGATAGAACCCTTCTGACATTCTG |
| Sm[12][1] | GCCAACAGCAAGTTACAAAATCGCTGATTGCT |
| Ss[12] | ACCTGAAAGCAATAAAAGGG |
| S[13][0] | CTCCAAAAGGAAACAAACATCAAGAAAACAAAAAAAAAAAGG |
| S[13][1] | GAAGATGATGAGCCTTTAATTGTATCGGTTTACTGAGCAAAA |
| S[13][2] | TTCGAGGTGAACGAATTATTCATTTCAATTACTCAGCTTGCT |
| Sb[14][0] | ACAATTTCATTTAACCTTGCTTCTGTAAATCGATTACATTTA |
| Sm[14][0] | GAACAAGCTACATAAATCAATATATTTTTAAT |
| Sm[14][1] | GGAAACAGAAGCCGTTTTTATTTTCTCATCGA |
| Ss[14] | TGTGAGTGAATGAATTACCT |
| Ef[0] | ATCATACAGGCTTGCGTTGCGCTCACTGCCCGCATCCAATAA |
| Ef[1] | CGTCTTTCCAGGTGCCTTGAGTAACAGTGCCCCGCTAACGAG |

**Re-entrant triangle**

For staples named X[][2] through X[][5], mX[][0], and mX[][1], toeholds are placed at the 3' end for the reconfiguration. Their releasers are fully complementary to the target staples for perfect base-pairing. To highlight their purposes, they are named after their targets, with 'r' in the front.

| Re-entrant triangle staples | |
|---|---|
| Name | Sequence (5' to 3') |
| M[0][0] | CATTGTGAATTGTACAACGGAGATTTGTATCACAACTTTAAT |
| M[0][1] | CGCGAAACAAAACCTTATGCGATTTTAAGAACTTATACCAAG |
| M[0][2] | TACCAGTCAGGCTCATCTTTGACCCCCAGCGATGGCTCATTA |
| M[0][3] | ACACTAAAACAACGTTGGGAAGAAAAATCTACGCAAAAGAAT |
| M[1][0] | ACCTCCCGACTCTGAGAGTCTGGAGCAAACAAGTTTTAGCGA |
| M[1][1] | CAGGTCATTGCTGCGGGAGGTTTTGAAGCCTTCAAAGGCTAT |
| M[1][2] | TAGTTGCTATTTAGCTATTTTTGAGAGATCTAAAATCAAGAT |
| M[1][3] | GCCGGAGAGGGTTGCACCCAGCTACAATTTTAATAAATTAAT |
| M[2][0] | CCGCTTCTGGTTATAAGCAAATATTTAAATTGCTTTCCGGCA |
| M[2][1] | CAGGAAGATTGGCCGGAAACCAGGCAAAGCGCGCCCCAAAAA |
| M[2][2] | TCAGGCTGCGCCCCCGGTTGATAATCAGAAAACATTCGCCAT |
| M[2][3] | AATCATATGTAAACTGTTGGGAAGGGCGATCGCTAGCATGTC |
| M[3][0] | GGGCGCTGGCAAATCCTGTTTGATGGTGGTTCGCGGGCGCTA |
| M[3][1] | CAGCAGGCGAAAGTGTAGCGGTCACGCTGCGCTGGTTTGCCC |
| M[3][2] | CACCCGCCGCGGTTGCAGCAAGCGGTCCACGCGTAACCACCA |
| M[3][3] | GCCCTGAGAGACTTAATGCGCCGCTACAGGGCTCACCGCCTG |
| M[4][0] | CGCCTGTAGCAAACAGTTAATGCCCCCTGCCTCAAACTACAA |
| M[4][1] | GTGCCCGTATATTCCACAGACAGCCCTCATAGTTGAGTAACA |
| M[4][2] | GATCTAAAGTTAGTTTTAACGGGGTCAGTGCCTTAGCGTAAC |
| M[4][3] | TACTGGTAATATTGTCGTCTTTCCAGACGTTATACAGGAGTG |
| Mb[5][0] | TAAAGGTGGCACAATAGAAAATTCATATGGTTAATACATACA |
| Mb[5][1] | ATTACTAGAAATGGTTTGAAATACCGACCGTGCGGAATCATA |
| Mm[5][0] | AAGGCGTTACGGAATAAGTTTATTGAAACGCA |
| Mm[5][1] | AAGACACCAAATAAGAATAAACACTGATAAAT |
| Ms[5] | TTGTCACAATACATATAAAA |
| Mb[6][0] | AGACGATTGGCGTCTCTGAATTTACCGTTCCAGAGGCAGGTC |
| Mb[6][1] | ACCTTTTACATAACAGAAATAAAGAAATTGCGCAGTAACAGT |
| Mm[6][0] | CAGGTTTATCATTAAAGCCAGAATACAAACAA |
| Mm[6][1] | ATAAATCCACGTCAGATGAATATATAGATTTT |
| Ms[6] | GGAAAGCGCACTTGATATTC |

| | |
|---|---|
| L[0][0] | CAATCATAAGGGAGTAGTAAATTGGGCTTGAGCGCAGACGGT |
| L[0][1] | AACACCAGAACGAACCGAACTGACCAACTTTGCCTGACGAGA |
| L[0][2] | GATGAACGGTGCATTCAGTGAATAAGGCTTGCAAAGAGGACA |
| L[0][3] | ACAAAGCTGCTTACAGACCAGGCGCATAGGCTAATCAACGTA |
| L[0][4] | AAATCAGGTCTTATTCATTGAATCCCCCTCAACATAAATCAA |
| L[0][5] | ATCGTCATAAATTACCCTGACTATTATAGTCAATACTGCGGA |
| L[0][6] | GGATTGCATCAGTTTAGACTGGATAGCGTCCAGAAGCAAAGC |
| L[0][7] | AATAGTAAAATAAAAGATTAAGAGGAAGCCCGCAGAGGGGGT |
| Lm[0][0] | AACAGTTCGACAAGAACCGGATATCATCAAGA |
| Lm[0][1] | GTAATCTTAGAAAACGAGAATGACATGCTTTA |
| Ls[0] | TCATTACCCAGGCTGACCTT |
| L[1][0] | CAACAGTAGGGAGTCCTGAACAAGAAAAATAAAGCCAACGCT |
| L[1][1] | AACAATAGATACTTAATTGAGAATCGCCATATCCTGTTTATC |
| L[1][2] | CAACATGTAATGTTCAGCTAATGCAGAACGCGTTAACAACGC |
| L[1][3] | ATAAACAACATTTAGGCAGAGGCATTTTCGAGGACGACGACA |
| L[1][4] | TAGACGGGAGATAACGTCAAAAATGAAAATAGGGAAGCGCAT |
| L[1][5] | GATTTTTTGTTATTAACTGAACACCCTGAACAAATAAGAAAC |
| L[1][6] | GTAATTGAGCGCATATTATTTATCCCAATCCAAAGTCAGAGG |
| L[1][7] | AAATAAACAGCCTAATATCAGAGAGATAACCCGCCAGTTACA |
| Lm[1][0] | ACAGAGAGGACAAAAGGTAAAGTAGAGAATAT |
| Lm[1][1] | AAAGTACCAATAACATAAAAACAGCAGCCTTT |
| Ls[1] | ATTCTGTCCACCAGTAATAA |
| L[2][0] | ACGCCATCAAAGTATCGGCCTCAGGAAGATCGACCAATAGGA |
| L[2][1] | GGACGACGACAAATAATTCGCGTCTGGCCTTCCAGTTTGAGG |
| L[2][2] | CTTTCATCAACGCATCGTAACCGTGCATCTGCCTGTAGCCAG |
| L[2][3] | TGTAGATGGGCATTAAATGTGAGCGAGTAACAGTCACGTTGG |
| L[2][4] | GAAACCTGTCGGTGCCTAATGAGTGAGCTAACTTCCAGTCGG |
| L[2][5] | TAAAGCCTGGGTGCCAGCTGCATTAATGAATCGCATAAAGTG |
| L[2][6] | CGGGGAGAGGCCACACAACATACGAGCCGGAAGGCCAACGCG |
| L[2][7] | GCTCACAATTCGGTTTGCGTATTGGGCGCCAGATTGTTATCC |
| Lm[2][0] | ATTGCGTTACGGCGGATTGACCGTTTCTCCGT |
| Lm[2][1] | GGGAACAAGCGCTCACTGCCCGCTTCACATTA |
| Ls[2] | AATGGGATAGACCCGTCGGA |
| L[3][0] | TTAGCGGGGTTCATGTACCGTAACACTGAGTTAGGATTAGGA |
| L[3][1] | CAATAGGAACCTTGCTCAGTACCAGGCGGATAATAGCAAGCC |
| L[3][2] | AGAGGGTTGATCCACCACCCTCATTTTCAGGGAGTGCCGTCG |
| L[3][3] | CACCCTCAGAGATAAGTATAGCCCGGAATAGGTCAGAACCGC |
| L[3][4] | GAACGAGGGTACTTGCAGGGAGTTAAAGGCCGGACAGCATCG |

| | |
|---|---|
| L[3][5] | GGTCGCTGAGGGCAACGGCTACAGAGGCTTTGCGATATATTC |
| L[3][6] | ACTTTTTCATGCAACCATCGCCCACGCATAACAGGACTAAAG |
| L[3][7] | GACAATGACAAAGGAAGTTTCCATTAAACGGGTAGTTGCGCC |
| Lm[3][0] | GGATCGTCTACCGCCACCCTCAGATACTCAGG |
| Lm[3][1] | AGGTTTAGACCCTCAGCAGCGAAACTTTTGCG |
| Ls[3] | ACCGCCACCCTGTATCACCG |
| L[4][0] | GCAGAGGCGAATTAGAATCCTTGAAAACATAGACAAAATCGC |
| L[4][1] | TTAATTTTCCCTTATTCATTTCAATTACCTGATCGCTATTAA |
| L[4][2] | ATGATGAAACATAACCTTGCTTCTGTAAATCGGCAAAAGAAG |
| L[4][3] | ATGTGAGTGAAAACATCAAGAAAACAAAATTAAAATCAATAT |
| L[4][4] | TCACCAATGAAATTTGGGAATTAGAGCCAGCAGCCGGAAACG |
| L[4][5] | CGACTTGAGCCACCATCGATAGCAGCACCGTATCACCGTCAC |
| L[4][6] | ACAGAATCAAGTTATTCATTAAAGGTGAATTAATCAGTAGCG |
| L[4][7] | ATTGACGGAAATTTGCCTTTAGCGTCAGACTGGAAGGTAAAT |
| Lm[4][0] | AGTAGCACTACCTTTTTAATGGAACAATTTC |
| Lm[4][1] | ATTTGAATCATTACCATTAGCAAGAAATCACC |
| Ls[4] | AACAGTACATATTACATTTA |
| L[5][0] | AACAAGAGTCCGTGGCGAGAAAGGAAGGGAAGTCCAGTTTGG |
| L[5][1] | AGCCGGCGAACACTATTAAAGAACGTGGACTCTGACGGGGAA |
| L[5][2] | GGGCGAAAAACGGGAGCCCCCGATTTAGAGCTCAACGTCAAA |
| L[5][3] | GGAACCCTAAACGTCTATCAGGGCGATGGCCCGCACTAAATC |
| L[5][4] | GCTCAATCGTCGAACAATATTACCGCCAGCCAACATTTTGAC |
| L[5][5] | GGTAATATCCATGAAATGGATTATTTACATTGCGGCCTTGCT |
| L[5][6] | CAGTCACACGAGAGTAGAAGAACTCAAACTATGCAGATTCAC |
| L[5][7] | ATCACTTGCCTCCAGTAATAAAAGGGACATTCTAGTAATAAC |
| Lm[5][0] | GGAAAAACGTTTTTTGGGGTCGAGCCATCACC |
| Lm[5][1] | CAAATCAAGCTCATGGAAATACCTTTGCAACA |
| Ls[5] | GTGCCGTAAAACTACGTGAA |
| Ef[0] | GTAGTAGCATTGAAAGGCCGGAGACAGTCAAATCTACTAATA |
| Ef[1] | AACTCCAACAGTAACCTGTTTAGCTATATTTTACCGGAAGCA |
| Ef[2] | AGATTAGAGCCACTTCTGAATAATGGAAGGGTAACAACTAAT |
| Ef[3] | AATATTTTTGAAATCAACAGTTGAAAGGAATTTGGCACAGAC |
| j[2][0]l | CTCCATGTTACTTAGCCGGAACGAGG |
| j[2][0]m | TCGCCTGATAAATTGTGTCGAAATCC |
| j[2][1]l | TTTTTGTTAAATCAGCTCATTTTTTA |
| j[2][1]m | TAAACGTTAATATTTTGTTAAAATTC |
| j[2][2]l | TTAAGAGGCTGAGACTCCTCAAGAGA |
| j[2][2]m | ATTTCGGAACCTATTATTCTGAAACA |

| | |
|---|---|
| j[2][3]l | TAGCCCGAGATAGGGTTGAGTGTTGT |
| j[2][3]m | CGAAATCGGCAAAATCCCTTATAAAT |
| j[3][0]f | GTAAATGAATTTTTTTTTTTGCTAAACCCACCAGAGCTTTTTTGACAGGAGG TT |
| j[3][0]s | GTAAGCGTCATTTTTTGCTTTTGATGA |
| j[3][1]f | GTGCGGGCCTTTTTTTTCCAGCTGGCGAAGAAGGCTTATTTTTTGAACGCGAG GC |
| j[3][1]s | GAGAATCGATTTTTTTTAATCGTAAAA |
| j[6][0]f | CATTTGGGGCTTTTTTTGTGGCATCAATTCACCATCAATTTTTTTCGTTCTAGCT G |
| j[6][0]ml | AAAGACTTCAAATATCGCGTTTTAATTCGAG |
| j[6][0]mr | ACGCTAACGAGCGTCTTTCCAGAGCCTAATTT |
| j[6][0]s | TCCTGAATCTTTTTTTAAGCGAACCAG |
| j[6][1]f | GTTAATAAAATTTTTTTCAACATTATTACTCCTTATTATTTTTTTCAAACGTAGA A |
| j[6][1]ml | TAAAATACGTAATGCCACTACGAAGGCACCA |
| j[6][1]mr | CAAAAGGGCGACATTCAACCGATTGAGGGAGG |
| j[6][1]s | TACCAGCGCCTTTTTTAAACGAAAGAG |
| j[6][2]f | GCGTACTATGTTTTTTTGCACGTATAACTTTTAGTTAATTTTTTTCCTAAATTTA A |
| j[6][2]ml | GGTGGTTTTTCTTTTCACCAGTGAGACGGGC |
| j[6][2]mr | CATATGCGTTATACAAATTCTTACCAGTATAA |
| j[6][2]s | AAGCCTGTTTTTTTTTCTGATTGCCCT |
| j[6][3]f | GAGGAAGGTTTTTTTTTTTAGGAGCACTTAGAACCTACTTTTTTTTTTGCACGTA A |
| j[6][3]ml | TGGCCAACAGAGATAGAACCCTTCTGACCTG |
| j[6][3]mr | CGGATTCGCCTGATTGCTTTGAATACCAAGTT |
| j[6][3]s | CGGGAGAAACTTTTTTGTAAGAATACG |
| X[0][0] | GATTCATCAGTAAAGAACTGGCATGATTAAGACAGGTAGAAA |
| X[0][1] | CGGAATACCCATGAGATTTAGGAATACCACATGCAATAATAA |
| X[0][2] | CAGATACATAACCAGAAGGAAACCGAGGAAACTCAACTAATGCGGTTTCC |
| X[0][3] | GAACAAAGTTACGCCAAAAGGAATTACGAGGCGCAGATAGCCCCACCTTC |
| X[0][4] | CAACACTATCAAGCCCTTTTTAAGAAAAGTAAATAGTAAGAGTTCTCTCA |
| X[0][5] | TATCTTACCGATAACCCTCGTTTACCAGACGATAGCAATAGCTCTGCTGA |
| X[0][6] | CAAAATAGCGATAAGAGCAAGAAACAATGAAACGATAAAAAC |
| X[0][7] | AAGCCCAATAAGAGGCTTTTGCAAAAGAAGTTGAATTGAGTT |
| X[1][0] | TGCTGCAAGGCAATAGCAAGCAAATCAGATATAAGGGGGATG |
| X[1][1] | TTACCGCGCCCGATTAAGTTGGGTAACGCCAGGTAGGAATCA |
| X[1][2] | GTCACGACGTTCAAGCCGTTTTTATTTTCATCGGTTTTCCCAATTTTCAA |
| X[1][3] | TCGAGAACAAGGTAAAACGACGGCCAGTGCCAACCGCACTCACGGGATCC |
| X[1][4] | CCTGCAGGTCGAGAACGGGTATTAAACCAAGTAGCTTGCATGAGTGGCGT |

| | |
|---|---|
| X[1][5] | TTATCATTCCAACTCTAGAGGATCCCCGGGTACTGTCTTTCCTTAGTTAG |
| X[1][6] | ATTCGTAATCATAGAAACCAATCAATAATCGGCCGAGCTCGA |
| X[1][7] | TTACGAGCATGTGGTCATAGCTGTTTCCTGTGCCATCCTAAT |
| X[2][0] | AGTTTCAGCGGTCAGAGCCGCCACCAGAACCAAACTTTCAAC |
| X[2][1] | AGCCACCACCCAGTGAGAATAGAAAGGAACAACCACCCTCAG |
| X[2][2] | TGCGAATAATAGAGCCGCCACCCTCAGAACCGCTAAAGGAATGTGTTGGG |
| X[2][3] | CGCCTCCCTCAATTTTTCACGTTGAAAATCTCCACCGGAACAACCCAGT |
| X[2][4] | GGCTCCAAAAGAAATCACCGGAACCAGAGCCACCAAAAAAAACCATCTTT |
| X[2][5] | TTTCATAATCAGAGCCTTTAATTGTATCGGTTTTTGCCATCTTGCCCTAG |
| X[2][6] | CTTTCGAGGTGGTCATAGCCCCCTTATTAGCGTATCAGCTTG |
| X[2][7] | CGGCATTTTCGAATTTCTTAAACAGCTTGATAGCGTTTTCAT |
| X[3][0] | CGTTAGAATCAGAGAAAACTTTTTCAAATATAGTGCTTTCCT |
| X[3][1] | ACAAAGAACGCGAGCGGGAGCTAAACAGGAGGCAATCGCAAG |
| X[3][2] | GGATTTTAGACATGTAAATGCTGATGCAAATCCCGATTAAAGGCGAATCC |
| X[3][3] | TATATAACTATAGGAACGGTACGCCAGAATCCTAGGTTGGGTCCCATGAG |
| X[3][4] | TTTTATAATCACTACCTTTTTAACCTCCGGCTTGAGAAGTGTACAGTCCG |
| X[3][5] | GGTCTGAGAGAGTGAGGCCACCGAGTAAAAGACAAAATCATAGTCTAAGA |
| X[3][6] | CACGCAAATTAAAGAGTCAATAGTGAATTTATGTCTGTCCAT |
| X[3][7] | AAGACGCTGAGACCGTTGTAGCAATACTTCTTAGCTTAGATT |
| mX[0][0] | CAGATACATAAAGCCCTTTTTACCGAGGAAACTCAACTAATGCCATACGT |
| mX[0][1] | TATCTTACCGACGCCAAAAGGTTACCAGACGATAGCAATAGCAATTCATC |
| mX[1][0] | GTCACGACGTTAGAACGGGTATTATTTTCATCGGTTTTCCCAGTGTTTAC |
| mX[1][1] | TTATCATTCCAGTAAAACGACATCCCCGGGTACTGTCTTTCCAATATTCT |
| mX[2][0] | TGCGAATAATAAAATCACCGGCCTCAGAACCGCTAAAGGAATTCAGAGCG |
| mX[2][1] | TTTCATAATCAATTTTTTCACTTGTATCGGTTTTTGCCATCTATATTCGT |
| mX[3][0] | GGATTTTAGACCTACCTTTTTTGATGCAAATCCCGATTAAAGCGGCATCA |
| mX[3][1] | GGTCTGAGAGAAGGAACGGTACGAGTAAAAGACAAAATCATATGAGGGTC |

| Re-entrant triangle releasers | |
|---|---|
| Name | Sequence (5' to 3') |
| rX[0][2] | GGAAACCGCATTAGTTGAGTTTCCTCGGTTTCCTTCTGGTTATGTATCTG |
| rX[0][3] | GAAGGTGGGGCTATCTGCGCCTCGTAATTCCTTTTGGCGTAACTTTGTTC |
| rX[0][4] | TGAGAGAACTCTTACTATTTACTTTTCTTAAAAAGGGCTTGATAGTGTTG |
| rX[0][5] | TCAGCAGAGCTATTGCTATCGTCTGGTAAACGAGGGTTATCGGTAAGATA |
| rX[1][2] | TTGAAAATTGGGAAAACCGATGAAAATAAAAACGGCTTGAACGTCGTGAC |
| rX[1][3] | GGATCCCGTGAGTGCGGTTGGCACTGGCCGTCGTTTTACCTTGTTCTCGA |
| rX[1][4] | ACGCCACTCATGCAAGCTACTTGGTTTAATACCCGTTCTCGACCTGCAGG |
| rX[1][5] | CTAACTAAGGAAAGACAGTACCCGGGGATCCTCTAGAGTTGGAATGATAA |

| | |
|---|---|
| rX[2][2] | CCCAACACATTCCTTTAGCGGTTCTGAGGGTGGCGGCTCTATTATTCGCA |
| rX[2][3] | ACTGGGTTGTTCCGGTGGAGATTTTCAACGTGAAAAAATTGAGGGAGGCG |
| rX[2][4] | AAAGATGGTTTTTTTTGGTGGCTCTGGTTCCGGTGATTTCTTTTGGAGCC |
| rX[2][5] | CTAGGGCAAGATGGCAAAAACCGATACAATTAAAGGCTCTGATTATGAAA |
| rX[3][2] | GGATTCGCCTTTAATCGGGATTTGCATCAGCATTTACATGTCTAAAATCC |
| rX[3][3] | CTCATGGGACCCAACCTAGGATTCTGGCGTACCGTTCCTATAGTTATATA |
| rX[3][4] | CGGACTGTACACTTCTCAAGCCGGAGGTTAAAAAGGTAGTGATTATAAAA |
| rX[3][5] | TCTTAGACTATGATTTTGTCTTTTACTCGGTGGCCTCACTCTCTCAGACC |
| rmX[0][0] | ACGTATGGCATTAGTTGAGTTTCCTCGGTAAAAAGGGCTTTATGTATCTG |
| rmX[0][1] | GATGAATTGCTATTGCTATCGTCTGGTAACCTTTTGGCGTCGGTAAGATA |
| rmX[1][0] | GTAAACACTGGGAAAACCGATGAAAATAATACCCGTTCTAACGTCGTGAC |
| rmX[1][1] | AGAATATTGGAAAGACAGTACCCGGGGATGTCGTTTTACTGGAATGATAA |
| rmX[2][0] | CGCTCTGAATTCCTTTAGCGGTTCTGAGGCCGGTGATTTTATTATTCGCA |
| rmX[2][1] | ACGAATATAGATGGCAAAAACCGATACAAGTGAAAAAATTGATTATGAAA |
| rmX[3][0] | TGATGCCGCTTTAATCGGGATTTGCATCAAAAAAGGTAGGTCTAAAATCC |
| rmX[3][1] | GACCCTCATATGATTTTGTCTTTTACTCGTACCGTTCCTTCTCTCAGACC |

**Rotating square**

| Rotating square staples | |
|---|---|
| Name | Sequence (5' to 3') |
| A[0][0] | AATAAGGCGTTAAATGTAAATCGAACGAGTAGATTGTAGAAA |
| A[0][1] | TGAAATACCGACCGAATCGCAAACAGTTGATTCCCGCTGTCT |
| A[0][2] | TCTGACCTAAATTTAGAAAACTTGCTCCGTTTCATAAGAACG |
| A[0][3] | CCAATCAATAATCGAATTCTGGCTGATGCAAATCCTGTGATA |
| A[0][4] | TTCCTTATCATTCCTCCATATAGACAAAGAACGCGAATGGTT |
| A[0][5] | GGTATTAAACCAAGCCTTTAATTTTTCAAATATATTTCATCT |
| A[1][0] | TTCGGTCGCTGAGGGTTAAAGGTCAGGATTAGAGACCTCAGA |
| A[1][1] | CGCCCACGCATAACATCGTCAACCGGAAGCAAACTAGCCACC |
| A[1][2] | GCCGACAATGACAAAGACAGCATCAAAATCAAAGCGGATAGC |
| A[1][3] | ACCGCCACCCTCAGCCAACAGGCCGCTTTTGCGGGCGATATA |
| A[1][4] | ACCCTCATTTTCAGGAACCAGCCCTCAGCAGCGAACAACCAT |
| A[1][5] | AAGCCCAATAGGAAACCATAAATCGGAACGAGGGTTAGTTGC |
| A[2][0] | CAGGAGGTTGAGGCGGAACCAATTACAGGTAGAAAAAGTATA |
| A[2][1] | CAGAGCCGCCGCCAACCGCCTACGAACTAACGGAATATCACC |
| A[2][2] | AGCCGCCACCAGAAACCCTCAGAGTAGTTCTACGTTAGTACC |
| A[2][3] | GCCCGGAATAGGTGCAACATTGAGCCACCACCGGAGCATTGA |
| A[2][4] | GTACTCAGGAGGTTTAATAAACCCTCAGAGCCGCCCCACCAC |
| A[2][5] | GCCACCCTCAGAACCCAGAACGAACCGCCACCCTCCCCTCAG |
| A[3][0] | ATAACCCACAAGAACCCAATATGCCCTGACGAGAAACTCATC |
| A[3][1] | TTGAGCGCTAATATCAATGAAGCTCATTCAGTGAACGTTTTT |
| A[3][2] | CTGAACAAAGTCAGCTTACCGACTGACCAACGTAAAATCATT |
| A[3][3] | GAGAACAAGCAAGCTAAGGCTATAAGAGCAAGAAACAGAGAG |
| A[3][4] | ATTTTCATCGTAGGCAAAGCTATAGCAATAGCTATAGGGTAA |
| A[3][5] | ACCGCGCCCAATAGGAACCGAAAGCCCTTTTTAAGGAACACC |
| M[0][0] | AGCGTAAGAATACGAAAGACAATAGTAGTAGCATTAAATCAT |
| M[0][1] | ATAGAACCCTTCTGTATTTTGAAGGTGGCATCAATGAATTAG |
| M[0][2] | GGGACATTCTGGCCAAATTCATCATTTGGGGCGCGAAAGCCT |
| M[0][3] | GTCACACGACCAGTGCCAAAGATAACCTGTTTAGCAAATCGG |
| M[0][4] | TTTACATTGGCAGATTCAACCGATACATTTCGCAATTATGAC |
| M[0][5] | CCTGTAATACTTTTACCATTAGATTGAGGGAGGGATGGATTA |
| M[0][6] | TTGTACCAAAAACAATGGTCAACAAAAGGGCGACATTCACCA |
| M[0][7] | CAGAGCATAAAGCTTATATTTTATGGTTTACCAGCAATAAAA |
| M[0][8] | CAAAATTAAGCAATAGCTGAATCACAATCAATAGAAACAGAG |
| M[0][9] | ACAGGCAAGGCAAATCTACTACCACGGAATAAGTTACCTGAA |
| M[1][0] | GGAACAAACGGCGGATCAATAACAGTTCAGAAAACAAGGGCG |

| | |
|---|---|
| M[1][1] | CAACCCGTCGGATTTGGATTAAATCCCCCTCAAATAGGGCGA |
| M[1][2] | ACATTAAATGTGAGGGAAGGGAATCGTCATAAATAAACCATC |
| M[1][3] | TCCTGTAGCCAGCTATCAAAAGGATAGCGTCCAATTTTTGGG |
| M[1][4] | AAAATAATTCGCGTAACAGAATAATAGTAAAATGTAAGCACT |
| M[1][5] | AAATCGGAACCCTAGAGGGGGATAAAGAAATTGCGGCCATCA |
| M[1][6] | GTCGAGGTGCCGTATTAGACTTTATTTGCACGTAACTGGCCT |
| M[1][7] | ACCCAAATCAAGTTACTGCGGTTAGAACCTACCATTTCATCA |
| M[1][8] | TGGCCCACTACGTGTTCATTGTACTTCTGAATAATCGAGTAA |
| M[1][9] | AAAAACCGTCTATCGCTTTAATAATCCTGATTGTTCTCCGTG |
| M[2][0] | GAGTTGCAGCAAGCGCGAAACTTGAGGATTTAGAACGCCATT |
| M[2][1] | CTTCACCGCCTGGCATTTGTAAGCCGTCAATAGATGTTGGGA |
| M[2][2] | GACGGGCAACAGCTATTGTGTCACTAACAACTAATGGGCCTC |
| M[2][3] | GGTTTTTCTTTTCATGCTCCATATCTAAAATATCTAGCTGGC |
| M[2][4] | TTGCGTATTGGGCGAACGAGGTTGAAAGGAATTGACTGCAAG |
| M[2][5] | GCGATTAAGTTGGGTCAACAGCGCAGACGGTCAATAGGCGGT |
| M[2][6] | GAAAGGGGGATGTGGGAAGGTTGTTACTTAGCCGGCCAGGGT |
| M[2][7] | TTCGCTATTACGCCTTAGGAGCGAAATCCGCGACCCCAGTGA |
| M[2][8] | AGGGCGATCGGTGCAGATTAGTCATCGCCTGATAAGATTGCC |
| M[2][9] | CAGGCTGCGCAACTAATACATAAAGTACAACGGAGCCTGAGA |
| M[3][0] | ATGAACGGTAATCGCAAACATCTTTTCATAATCAAGCGTACT |
| M[3][1] | CTGGAGCAAACAAGTAATTACCCCCTTATTAGCGTGAGCACG |
| M[3][2] | ATCAGGTCATTGCCTTTGAATATCGGCATTTTCGGCTCGTTA |
| M[3][3] | TTTGAGAGATCTACGAAACAGGTCAGACTGTAGCGGCTAAAC |
| M[3][4] | ATGCCGGAGAGGGTTATGTGAAGAATCAAGTTTGCAGGGATT |
| M[3][5] | TTAGACAGGAACGGTAGCGACGTGAATAACCTTGCTAAATTA |
| M[3][6] | AGGAGGCCGATTAACTTTAGCTACATAAATCAATAAGCTATT |
| M[3][7] | GAATCAGAGCGGGACGTTTTCTACCTTTTTTAATGAAAGGCT |
| M[3][8] | TATAACGTGCTTTCTCATAGCATTTAACAATTTCATGAGAGT |
| M[3][9] | ATGGTTGCTTTGACTTGCCATCAAGAAAACAAAATAGAATCG |
| L[0][0] | TAATGCGCCGCTACAGTTGAGGCAAAAGAAGATGACTAGCAT |
| L[0][1] | AACCACCACACCCGCATTCAATTATTCATTTCAATCCCCGGT |
| L[0][2] | TGTAGCGGTCACGCTAACGCCACAAAATCGCGCAGGCCCCAA |
| L[0][3] | GGGCGCTAGGGCGCGGCATAGGATTGCTTTGAATATATAAGC |
| L[0][4] | AGGGAAGAAAGCGATCATAACGAAACAATAACGGATAAACGT |
| L[0][5] | GGCGAACGTGGCGACGACGATACAGTACCTTTTACAATTCGC |
| L[0][6] | TAGAGCTTGACGGGCGAGAGGACGTCAGATGAATAAAATCAG |
| L[0][7] | CTCATTTTTTAACCAGGTTTACTTTTGCAAAAGAACCCGATT |
| L[0][8] | ATTAAATTTTTGTTTACAGTAAAAAACCAAAATAGGAAAGCC |

| | |
|---|---|
| L[0][9] | TAATATTTTGTTAAATCGGGACCTCGTTTACCAGAGAAAGGA |
| L[0][10] | AAATATTTAAATTGTTCGCCTTAAGAGCAACACTAAAGGAGC |
| L[0][11] | AAACAGGAAGATTGCCAAGTTAAAAGGAATTACGATGGCAAG |
| L[0][12] | TGATAATCAGAAAAAGGCGAACTAATGCAGATACATGCGCGT |
| L[0][13] | GTCAATCATATGTATACCTGAATTTAGGAATACCACCGCGCT |
| L[1][0] | CGGAAACCAGGCAATTTACAAGACCCCCAGCGATTTGGTTTG |
| L[1][1] | TTCCGGCACCGCTTCGTATTATACACTAAAACACTAATCCTG |
| L[1][2] | AAGATCGCACTCCAACGTTATAAAACGAAAGAGGCCGAAATC |
| L[1][3] | GACGACAGTATCGGTGAGTAACCACTACGAAGGCATAAATCA |
| L[1][4] | CATCTGCCAGTTTGGGAACAAAAACGGGTAAAATAGATAGGG |
| L[1][5] | GATGGGCGCATCGTGGAGCGGTTTCATGAGGAAGTAGTTTGG |
| L[1][6] | GGGATAGGTCACGTTCCTGATGGCTTTGAGGACTAATTAAAG |
| L[1][7] | AACGTGGACTCCAACTACAGATATCAGATGATGGCCCGTAAT |
| L[1][8] | AACAAGAGTCCACTAAGACTTAATTATCATCATATTGGTGTA |
| L[1][9] | TTGAGTGTTGTTCCTTCCATTAGAAACCACCAGAAAACCGTG |
| L[1][10] | AAAGAATAGCCCGACGTAATGCATTATCATTTTGCAGGGGAC |
| L[1][11] | GGCAAAATCCCTTACCAACCTTAATTTTAAAAGTTCCTCAGG |
| L[1][12] | TTTGATGGTGGTTCAAAAGAAAATCCTTTGCCCGAGCCAGCT |
| L[1][13] | CCCCAGCAGGCGAACATCTTTACAATTCGACAACTCTGGTGC |
| L[2][0] | GACGCTCAATCGTCAATATTGTAGGTTGGGTTATAAGAAGCC |
| L[2][1] | TCATGGAAATACCTTTAAAGGCTACCTTTTTAACCAGGATAA |
| L[2][2] | CCATTGCAACAGGACACCGACCAAAATCATAGGTCCTCATAT |
| L[2][3] | CCAGAACAATATTAAATTAGAAAGAGTCAATAGTGGCCTGAG |
| L[2][4] | TATCGGCCTTGCTGCAGTAGCAGCTTAGATTAAGAAGATTCA |
| L[2][5] | CCTGAGTAGAAGAACAAGGCCAATCCTTGAAAACAGCCGGAG |
| L[2][6] | GATTAGTAATAACATGAAACCTATTAATTAATTTTATCAATA |
| L[2][7] | TGATATTCAACCGTTCGTCGCATCGATAGCAGCACCTTCTTT |
| L[2][8] | ACAGTCAAATCACCCCCTTAGGGAAACGTCACCAATCACTTG |
| L[2][9] | AAAGGGTGAGAAAGTAGCGATACCATTACCATTAGCTCAAAC |
| L[2][10] | TAATGTGTAGGTAACGCTGAGGCCAGCAAAATCACGTAATAT |
| L[2][11] | ATTTTAAATGCAATAATTTATTTGAGCCATTTGGGCCGCCAG |
| L[2][12] | AAATTTTTAGAACCTGAGAGATGAATTATCACCGTAAAACGC |
| L[2][13] | TTTATTTCAACGCATCCGGCTACGGAAATTATTCAACATTTT |
| L[3][0] | AATTGTTATCCGCTCAGCAGAGTGGCAACATATAACACAACA |
| L[3][1] | TCATAGCTGTTTCCTGAGGCGGTAGAAAATACATAATAAAGT |
| L[3][2] | AGCTCGAATTCGTACGCCTGCTTACGCAGTATGTTGCCTAAT |
| L[3][3] | CTAGAGGATCCCCGGAGAGCCTGGCATGATTAAGAACATTAA |
| L[3][4] | TTGCATGCCTGCAGAATCTAATAACGGAATACCCACTGCCCG |

| | |
|---|---|
| L[3][5] | AAACGACGGCCAGTGAACCTCGAAACCGAGGAAACAACCTGT |
| L[3][6] | TTTCCCAGTCACGACAATCAAGCCGAACAAAGTTATAATGAA |
| L[3][7] | TCGGCCAACGCGCGGCAGATATATCTGGTCAGTTGCCAGGGT |
| L[3][8] | CGTGCCAGCTGCATCCAGAAGAAATATCAAACCCTCGTTGTA |
| L[3][9] | CTTTCCAGTCGGGAGCAATAAAGCATCACCTTGCTGCCAAGC |
| L[3][10] | TTGCGTTGCGCTCAAAAGAACAGCAGCAAATGAAAGTCGACT |
| L[3][11] | GAGTGAGCTAACTCCTCCTTAAACAGTGCCACGCTGGTACCG |
| L[3][12] | GTAAAGCCTGGGGTAGCAAACGTCAGTATTAACACATCATGG |
| L[3][13] | TACGAGCCGGAAGCCATAAAGAGATAAAACAGAGGTGTGTGA |
| Ef[0] | GAAGTGTTTTCACGCAAATTAACCGTTGTAGCAGAATCCTGA |
| Ef[1] | TTTGAATGGCGCCATTAAAAATACCGAACGACAGACAATATT |
| mA[0][0] | AATCGCCATATTTATGTAAATCGAACGAGTAGATTGTTTATC |
| mA[0][1] | CAACAGTAGGGCTTAATCGCAAACAGTTGATTCCCCCTGAAC |
| mA[0][2] | TACCAGTATAAAGCAGAAAACTCTGGAAGTTTCATCCCATCC |
| mA[0][3] | CATATGCGTTATACTTTAGTTATATGCAACTAAAGGTAGAAA |
| mA[0][4] | TAGAAAAAGCCTGTACCTAAACTGTAGCTCAACATGCTGTCT |
| mA[0][5] | AAACACCGGAATCAATACCGATTGCTCCTGCTGAAAAGAACG |
| mA[0][6] | AACAATAGATAAGTAATTCTGGCTGATGCAAATCCAATTGAG |
| mA[0][7] | AAGAAAAATAATATTCCATATAGACAAAGAACGCGCAACGCT |
| mA[0][8] | TAATTTACGAGCATTACGGTGTTTTTCAAATATATAAATTCT |
| mA[0][9] | CCAATCAATAATCGGTTTTAAAATTTCATCTTCTGTTAGTAT |
| mA[0][10] | TTCCTTATCATTCCTATAATGTTTAATGGTTTGAATAATTAC |
| mA[0][11] | GGTATTAAACCAAGCCTTTAACCGTGTGATAAATATAAGAAT |
| mA[1][0] | CAGCTTGATACCGAGACAATGGTCAGGATTAGAGACCTCAGA |
| mA[1][1] | TTTCGAGGTGAATTCCACGCAACCGGAAGCAAACTAGCCACC |
| mA[1][2] | TGTATCGGTTTATCGGTCGCTTCGAGCTTCAAAGCGGATAGC |
| mA[1][3] | GCTCCAAAAGGAGCGTTAAAGCTTCAAATATCGCGCCCATGT |
| mA[1][4] | TTGAAAATCTCCAAATCGTCAATTAAGAGGAAGCCTTTCGTC |
| mA[1][5] | GCGAATAATAATTTAGACAGCATCAAAAAATTGCATCAACGCC |
| mA[1][6] | ACCGCCACCCTCAGCCAACAGACAACAACCATCGCTCTTAAA |
| mA[1][7] | ACCCTCATTTTCAGGAACCAGTAACCGATATATTCAGCTTGC |
| mA[1][8] | AAGCCCAATAGGAATTTTAATGAGGCTTGCAGGGACTTTAAT |
| mA[1][9] | ACCGTAACACTGAGCGAAAGAGCCGCTTTTGCGGGAAAAAAG |
| mA[1][10] | ACCAGTACAAACTACAAAAAGCCCTCAGCAGCGAATTTCACG |
| mA[1][11] | TGTAGCATTCCACAACCATAAATCGGAACGAGGGTAGGAATT |
| mA[2][0] | ATGATACAGGAGTGGGAACCAATTACAGGTAGAAAGATTAGG |
| mA[2][1] | AAGCGTCATACATGACCGCCTACGAACTAACGGAAGCTCAGT |
| mA[2][2] | CTCTGAATTTACCGACCCTCAAGAAAAATCTACGTTGCCGTC |

| | |
|---|---|
| mA[2][3] | AGCCAGAATGGAAAAGAGCCAATACCAGTCAGGACAAGTATA |
| mA[2][4] | AAACAAATAAATCCGCCACCAGATTTTAAGAACTGTATCACC |
| mA[2][5] | ACGATTGGCCTTGAGCCGCCGGAGTAGTGAATTACTAGTACC |
| mA[2][6] | ATTAGCGGGGTTTTCAACATTGAGCCACCACCGGAGCTTTTG |
| mA[2][7] | ACCAGGCGGATAAGTAATAAACCCTCAGAGCCGCCTTCCAGT |
| mA[2][8] | GAGAGGGTTGATATGTTGGGAGAACCGCCACCCTCGCGCAGT |
| mA[2][9] | GCCCGGAATAGGTGGCTCATTCCACCCTCAGAGCCTCATTAA |
| mA[2][10] | GTACTCAGGAGGTTCTTATGCGAACCACCACCAGATATTCAC |
| mA[2][11] | GCCACCCTCAGAACCCAGAACCCAGCATTGACAGGAGGTCAG |
| mA[3][0] | CGGGAGAATTAACTAACAAAGTGCCCTGACGAGAAACTCATC |
| mA[3][1] | AAAACAGGGAAGCGAGCGCTAGCTCATTCAGTGAACGTTTTT |
| mA[3][2] | CTTTACAGAGAGAAACCCACACCAAATCAACGTAAAATCATT |
| mA[3][3] | GTCAAAAATGAAAACCCAATACAAGAACCGGATATCAAGCAA |
| mA[3][4] | AGAAACGATTTTTTCAATGAACTTCATCAAGAGTAGCTTATC |
| mA[3][5] | TTATTTATCCCAATCTTACCGACTGACCATAGGCTCGCGAGG |
| mA[3][6] | GAGAACAAGCAAGCTAAGGCTTCAGAGGGTAATTGCATTAGA |
| mA[3][7] | ATTTTCATCGTAGGCAAAGCTATATCAGAGAGATATAACATA |
| mA[3][8] | ACCGCGCCCAATAGTCATTACAGAATTGAGTTAAGTAGCAGC |
| mA[3][9] | ATCAGATATAGAAGATCTTGAATAAGAGCAAGAAAGTTTAAC |
| mA[3][10] | CGGTATTCTAAGAAGGCTGACATAGCAATAGCTATCCAAATA |
| mA[3][11] | CGTTTTAGCGAACCGAACCGAAAGCCCTTTTTAAGAGCCATA |
| fA[0][0] | AAGGTAAAGTAATTTGTAAATCGAACGAGTAGATTGACGACA |
| fA[0][1] | GAGAATATAAAGTAAATCGCAAACAGTTGATTCCCCAGCTAA |
| fA[0][2] | GCATTTTCGAGCCAAGAAAACTCTGGAAGTTTCATGTTTATC |
| fA[0][3] | CAACATGTAATTTATTTAGTTATATGCAACTAAAGCCTGAAC |
| fA[0][4] | AATCGCCATATTTAACCTAAACTGTAGCTCAACATCCCATCC |
| fA[0][5] | CAACAGTAGGGCTTATACCGAGCTTAATTGCTGAAGTAGAAA |
| fA[0][6] | TACCAGTATAAAGCAGGCGTTCATTTTTGCGGATGGCTGTCT |
| fA[0][7] | CATATGCGTTATACACCGGAATTGCTCCTTTTGATAAGAACG |
| fA[0][8] | GGTATTAAACCAAGCCTTTAATCATAATTACTAGATTAGTAT |
| fA[0][9] | TTCCTTATCATTCCAAGAGGTAAATAAGAATAAACAAATTCT |
| fA[0][10] | CCAATCAATAATCGGCTTAGACCGTGTGATAAATACAACGCT |
| fA[0][11] | TAATTTACGAGCATTATAATGTTTAATGGTTTGAAAATTGAG |
| fA[0][12] | AAGAAAAATAATATGTTTTAAAATTTCATCTTCTGACAACGC |
| fA[0][13] | AACAATAGATAAGTTACGGTGTTTTTCAAATATATGGCAGAG |
| fA[0][14] | TGCAGAACGCGCCTTCCATATAGACAAAGAACGCGGTAATAA |
| fA[0][15] | ATAAACAACATGTTAATTCTGGCTGATGCAAATCCCCGACAA |
| fA[1][0] | TGTATCGGTTTATCCGAGGTGGTCAGGATTAGAGACCTCAGA |

| | |
|---|---|
| fA[1][1] | GCTCCAAAAGGAGCCTTGATAACCGGAAGCAAACTAGCCACC |
| fA[1][2] | TTGAAAATCTCCAAGACAATGTCGAGCTTCAAAGCGGATAGC |
| fA[1][3] | GCGAATAATAATTTCCACGCACTTCAAATATCGCGCCCATGT |
| fA[1][4] | AAAGGAACAACTAAGGTCGCTATTAAGAGGAAGCCTTTCGTC |
| fA[1][5] | GTTTCAGCGGAGTGGTTAAAGAAAGCGGATTGCATCAACGCC |
| fA[1][6] | TTTGCTAAACAACTATCGTCACTGACTATTATAGTGACAGCC |
| fA[1][7] | AATGAATTTTCTGTAGACAGCATCAAAAATCAGGTAACGATC |
| fA[1][8] | TAAAGTTTTGTCGTACCATAAATCGGAACGAGGGTGTTAGTA |
| fA[1][9] | CTCATAGTTAGCGTCTTTACCCCCTCAGCAGCGAAATGGGAT |
| fA[1][10] | TGTAGCATTCCACACAGAAGCGCCGCTTTTGCGGGTTCAACA |
| fA[1][11] | ACCAGTACAAACTACAAAAAGGAGGCTTGCAGGGAAGAATAG |
| fA[1][12] | ACCGTAACACTGAGCGAAAGATAACCGATATATTCAGGAATT |
| fA[1][13] | AAGCCCAATAGGAATTTTAATACAACAACCATCGCTTTCACG |
| fA[1][14] | ACCCTCATTTTCAGGAACCAGCCGATAGTTGCGCCAAAAAAG |
| fA[1][15] | ACCGCCACCCTCAGCCAACAGAATTTCTTAAACAGCTTTAAT |
| fA[2][0] | GCCTATTTCGGAACGGAACCAATTACAGGTAGAAAGAAACAT |
| fA[2][1] | TATAAACAGTTAATACCGCCTACGAACTAACGGAAGCTGAGA |
| fA[2][2] | TGCCTTGAGTAACAACCCTCAAGAAAAATCTACGTGATTAGG |
| fA[2][3] | AATAAGTTTTAACGAGAGCCAATACCAGTCAGGACGCTCAGT |
| fA[2][4] | ATGATACAGGAGTGGCCACCAGATTTTAAGAACTGTGCCGTC |
| fA[2][5] | AAGCGTCATACATGGCCGCCGTCATTGTGAATTACAAGTATA |
| fA[2][6] | CTCTGAATTTACCGAGGTTGAATGGTTTAATTTCATATCACC |
| fA[2][7] | AGCCAGAATGGAAATTGGCCTGAGTAGTAAATTGGTAGTACC |
| fA[2][8] | GCCACCCTCAGAACCCAGAACTGATATTCACAAACTCATTAA |
| fA[2][9] | GTACTCAGGAGGTTGCTTGAGGGCAGGTCAGACGAGCGCAGT |
| fA[2][10] | GCCCGGAATAGGTGACTTTAACCAGCATTGACAGGTTCCAGT |
| fA[2][11] | GAGAGGGTTGATATCTTATGCGAACCACCACCAGAGCTTTTG |
| fA[2][12] | ACCAGGCGGATAAGGCTCATTCCACCCTCAGAGCCTACTGGT |
| fA[2][13] | ATTAGCGGGGTTTTGTTGGGAGAACCGCCACCCTCGGGTCAG |
| fA[2][14] | CTCCTCAAGAGAAGTAATAAACCCTCAGAGCCGCCGTGCCCG |
| fA[2][15] | GAAAGTATTAAGAGCAACATTGAGCCACCACCGGAGCCCCCT |
| fA[3][0] | CTTTACAGAGAGAAACAGGGATGCCCTGACGAGAAACTCATC |
| fA[3][1] | GTCAAAAATGAAAAGAGAATTGCTCATTCAGTGAACGTTTTT |
| fA[3][2] | AGAAACGATTTTTTAACAAAGCCAAATCAACGTAAAATCATT |
| fA[3][3] | TTATTTATCCCAATAGCGCTACAAGAACCGGATATCAAGCAA |
| fA[3][4] | GTTACAAAATAAACACCCACACTTCATCAAGAGTAGCTTATC |
| fA[3][5] | TTTCCAGAGCCTAACCCAATACAGGCGCATAGGCTCGCGAGG |
| fA[3][6] | TTACCAACGCTAACCAATGAAACAGATGAACGGTGTCCCGAC |

| | |
|---|---|
| fA[3][7] | GCTACAATTTTATCCTTACCGACTGACCAACTTTGGAAGCCT |
| fA[3][8] | TAAATCAAGATTAGGAACCGAAAGCCCTTTTTAAGGCACCCA |
| fA[3][9] | TTGCGGGAGGTTTTAAAGAGGATAGCAATAGCTATCTGAATC |
| fA[3][10] | CGTTTTAGCGAACCTACAGACATAAGAGCAAGAAAGAGCGTC |
| fA[3][11] | CGGTATTCTAAGAAGGCTGACAGAATTGAGTTAAGTTTGCCA |
| fA[3][12] | ATCAGATATAGAAGATCTTGAATATCAGAGAGATAAGCCATA |
| fA[3][13] | ACCGCGCCCAATAGTCATTACTCAGAGGGTAATTGCCAAATA |
| fA[3][14] | ATTTTCATCGTAGGCAAAGCTAACTGAACACCCTGGTTTAAC |
| fA[3][15] | GAGAACAAGCAAGCTAAGGCTAGCGCATTAGACGGTAGCAGC |

**Re-entrant triangle, control**

| Re-entrant triangle, control, staples | |
|---|---|
| Name | Sequence (5' to 3') |
| M[0][0] | CGCCTCCCTCAACCAGGCGGATAAGTGCCGTCCCACCGGAAC |
| M[0][1] | ATATAAGTATAAAATCACCGGAACCAGAGCCAGAGAGGGTTG |
| M[0][2] | TTTCATAATCAGCCCGGAATAGGTGTATCACCTTTGCCATCT |
| M[0][3] | GGTTTAGTACCGTCATAGCCCCCTTATTAGCGGTACTCAGGA |
| M[0][4] | CGGCATTTTCGGCCACCCTCAGAACCGCCACCGCGTTTTCAT |
| M[0][5] | CCACCCTCAGACCTTTAGCGTCAGACTGTAGCCTCAGAACCG |
| M[1][0] | CTTATCATTCCGCATTAGACGGGAGAATTAACGCTGTCTTTC |
| M[1][1] | GAACAAAGTCAGTAGAAACCAATCAATAATCGTGAACACCCT |
| M[1][2] | TTTACGAGCATGAGGGTAATTGAGCGCTAATACCCATCCTAA |
| M[1][3] | AACCCACAAGACCTGAACAAGAAAAATAATATTCAGAGAGAT |
| M[1][4] | AATAGATAAGTATTGAGTTAAGCCCAATAATAGTTTATCAAC |
| M[1][5] | ACAATGAAATACAGCTAATGCAGAACGCGCCTAGAGCAAGAA |
| M[2][0] | TAAGAGCAACAAGTAACAGTACCTTTTACATCCGAGGCATAG |
| M[2][1] | ATAACGGATTCACATAACGCCAAAAGGAATTAGGGAGAAACA |
| M[2][2] | CTAATGCAGATGCCTGATTGCTTTGAATACCACCACATTCAA |
| M[2][3] | TCGCGCAGAGGATCAGTTGAGATTTAGGAATAAGTTACAAAA |
| M[2][4] | TAGAAAGATTCCGAATTATTCATTTCAATTACTTATTACAGG |
| M[2][5] | GAAGATGATGAAAACGAACTAACGGAACAACACTGAGCAAAA |
| M[3][0] | GGGTGGTTTTTGAAGGGAAGAAAGCGAAAGGATTGGGCGCCA |
| M[3][1] | GGGCGCTGGCACGGGGAGAGGCGGTTTGCGTAGCGGGCGCTA |
| M[3][2] | CGGCCAACGCGAGTGTAGCGGTCACGCTGCGCATTAATGAAT |
| M[3][3] | CACCCGCCGCGGAAACCTGTCGTGCCAGCTGCGTAACCACCA |
| M[3][4] | TTTCCAGTCGGCTTAATGCGCCGCTACAGGGCCACTGCCCGC |
| M[3][5] | GTTGCTTTGACTCACATTAATTGCGTTGCGCTGCGTACTATG |
| M[4][0] | GCCTTTATTTCTAACCAATAGGAACGCCATCATGCGGGAGAA |
| M[4][1] | GCGTCTGGCCTATTATGACCCTGTAATACTTTAAAATAATTC |
| M[4][2] | GTACCAAAAACTCCTGTAGCCAGCTTTCATCATAAATCGGTT |
| M[4][3] | TGAGCGAGTAATAAAGCCTCAGAGCATAAAGCACATTAAATG |
| M[4][4] | AAATTAAGCAACAACCCGTCGGATTCTCCGTGAGAATTAGCA |
| M[4][5] | GCGGATTGACCTAAATCATACAGGCAAGGCAAGGAACAAACG |
| Mb[5][0] | CGAACGAGTAGTCTACTAATAGTAGTAGCATTCCCAATTCTG |
| Mb[5][1] | GTGGCATCAATATTTAGTTTGACCATTAGATAAGCTGAAAAG |
| Mb[5][2] | TAAAATACGTAGCAACGGCTACAGAGGCTTTGATTAAACGGG |
| Mb[5][3] | GAACGAGGGTAATGCCACTACGAAGGCACCAAGACAGCATCG |
| Mm[5][0] | TAACCTGTTCATGAGGAAGTTTCCAGGACTAA |

| | |
|---|---|
| Mm[5][1] | AGACTTTTTAGCTATATTTCATATGGTCAA |
| Ms[5] | TTGGGGCGCGCATTTCGCAA |
| Mb[6][0] | AATTGAGAATCCTGTCCAGACGACGACAATAAAGTAGGGCTT |
| Mb[6][1] | GTAAAGTAATTGCCATATTTAACAACGCCAACCCGACAAAAG |
| Mb[6][2] | CGCCTGCAACATAAAAATACCGAACGAACCACGTATTAACAC |
| Mb[6][3] | AACATCGCCATGTGCCACGCTGAGAGCCAGCAATAGCCCTAA |
| Mm[6][0] | CATTTTCGAGAGGTGAGGCGGTCACAGCAGAA |
| Mm[6][1] | GATAAAACAGCCAGTAATAAGAGAGGCAGAGG |
| Ms[6] | ATATAAAGTAATGTAATTTA |
| L[0][0] | CAGACGACGATAAAAAGATTAAGAGGAAGCCCCCTCGTTTAC |
| L[0][1] | CGGATTGCATCAAAAACCAAAATAGCGAGAGGAGAAGCAAAG |
| L[0][2] | AGAAGTTTTGCTTTACCCTGACTATTATAGTCCTTTTGCAAA |
| L[0][3] | AAAATCAGGTCCAGAGGGGGTAATAGTAAAATCCATAAATCA |
| L[0][4] | GATAGCGTCCACAGTTCAGAAAACGAGAATGAGTTTAGACTG |
| L[0][5] | AGAATACACTAGGCGCATAGGCTGGCTGACCTAAGAGGCAAA |
| L[0][6] | TGTACAGACCAAAACACTCATCTTTGACCCCCAGATGAACGG |
| L[0][7] | CCAAGCGCGAATGACCAACTTTGAAAGAGGACAGCGATTATA |
| L[0][8] | GGGAACCGAACACAAAGTACAACGGAGATTTGTCAATCATAA |
| L[0][9] | TGATAAATTGTGCCGGAACGAGGCGCAGACGGTATCATCGCC |
| Lm[0][0] | AAATATTCTGCTCCATGTTACTTAGTCGAAAT |
| Lm[0][1] | CCGCGACCATTGAATCCCCCTCAAATCGTCAT |
| Ls[0] | ATGCTTTAAAATACTGCGGA |
| L[1][0] | AAACAAATAAAAAGGATTAGGATTAGCGGGGTTGATATTCAC |
| L[1][1] | CTCCTCAAGAGTCCTCATTAAAGCCAGAATGGGAGGCTGAGA |
| L[1][2] | CTCTGAATTTATCTGAAACATGAAAGTATTAAAAAGCGCAGT |
| L[1][3] | GAACCTATTATCCGTTCCAGTAAGCGTCATACGCCTATTTCG |
| L[1][4] | ATGATACAGGATATAAACAGTTAATGCCCCCTATGGCTTTTG |
| L[1][5] | AGGAATTGCGATTTTGCGGGATCGTCACCCTCGAACAACTAA |
| L[1][6] | TTAAAGGCCGCATAATAATTTTTTCACGTTGATTGCAGGGAG |
| L[1][7] | AAAAAAGGCTCGATATATTCGGTCGCTGAGGCAAATCTCCAA |
| L[1][8] | CACGCATAACCCAAAAGGAGCCTTTAATTGTAAACCATCGCC |
| L[1][9] | AGCTTGCTTTCAGTTGCGCCGACAATGACAACTCGGTTTATC |
| Lm[1][0] | TTAACGGGACAGCTTGATACCGATGAGGTGAA |
| Lm[1][1] | TTTCTTAAGTCAGTGCCTTGAGTAAATAAGTT |
| Ls[1] | ACAGTGCCCGGTGTACTGGT |
| L[2][0] | ACGCGAGGCGTTACAGAGAGAATAACATAAAATATTCTAAGA |
| L[2][1] | ATAGCAGCCTTTTTAGCGAACCTCCCGACTTGAAAAATGAAA |
| L[2][2] | TGAAGCCTTAAAACGATTTTTTGTTTAACGTCCGGGAGGTTT |

| | |
|---|---|
| L[2][3] | TCCAAATAAGAATCAAGATTAGTTGCTATTTTTTATCCCAA |
| L[2][4] | ACAATTTTATCACAAAATAAACAGCCATATTAGCACCCAGCT |
| L[2][5] | ATTCATATGGTACCGTAATCAGTAGCGACAGATCAATAGAAA |
| L[2][6] | TCGATAGCAGCTTACCAGCGCCAAAGACAAAAAATGAAACCA |
| L[2][7] | CAACCGATTGAAGCAAGGCCGGAAACGTCACCGGGCGACATT |
| L[2][8] | CCATTACCATTGGGAGGGAAGGTAAATATTGAACCAGTAGCA |
| L[2][9] | TCATTAAAGGTGGAATTAGAGCCAGCAAAATCCGGAAATTAT |
| Lm[2][0] | AACGAGCGCGACTTGAGCCATTTGGAATTATC |
| Lm[2][1] | ACCGTCACTCTTTCCAGAGCCTAACCAACGCT |
| Ls[2] | TTTGCCAGTTCTGAATCTTA |
| L[3][0] | AATTAATTACAGGTTGGGTTATATAACTATATAAGAAAACAA |
| L[3][1] | CCTCCGGCTTATTTAACAATTTCATTTGAATTACCTTTTTAA |
| L[3][2] | ATGGAAACAGTAAATCATAGGTCTGAGAGACTACCTTTTTA |
| L[3][3] | TGAATTTATCAACATAAATCAATATATGTGAGGAGTCAATAG |
| L[3][4] | TGCTTCTGTAACTTAGATTAAGACGCTGAGAATGAATAACCT |
| L[3][5] | ATCACCTTGCTCGAACGTTATTAATTTTAAAAAATCTAAAGC |
| L[3][6] | AATCCTTTGCCGAACCTCAAATATCAAACCCTACTCGTATTA |
| L[3][7] | CTGGTCAGTTGGACTTTACAAACAATTCGACACAATCAATAT |
| L[3][8] | TAGAAGTATTAGCAAATCAACAGTTGAAAGGATTTGAGGATT |
| L[3][9] | GTTATCTAAAAGAGCCGTCAATAGATAATACAATTGAGGAAG |
| Lm[3][0] | TTTTCCCTAACAACTAATAGATTATATCTTTA |
| Lm[3][1] | GGAGCACTTAGAATCCTTGAAAACTTAATTAA |
| Ls[3] | ATAGCGATAGATCGTCGCTA |
| L[4][0] | CAAAATCCCTTACGGGGAAAGCCGGCGAACGTCCGAAATCGG |
| L[4][1] | TTTAGAGCTTGATAAATCAAAAGAATAGCCCGGAGCCCCCGA |
| L[4][2] | GAGTGTTGTTCACTAAATCGGAACCCTAAAGGAGATAGGGTT |
| L[4][3] | TGCCGTAAAGCCAGTTTGGAACAAGAGTCCACGGGGTCGAGG |
| L[4][4] | CGTGGACTCCAATCACCCAAATCAAGTTTTTTATTAAAGAA |
| L[4][5] | AATATCCAGAAGAATGGCTATTAGTCTTTAATCCTTGCTGGT |
| L[4][6] | ACAATATTTTTCAATATTACCGCCAGCCATTGCGTGGCACAG |
| L[4][7] | AACGCTCATGGTGACCTGAAAGCGTAAGAATACAACAGGAAA |
| L[4][8] | TAGAACCCTTCAAATACCTACATTTTGACGCTCCAACAGAGA |
| L[4][9] | AAATGGATTATGTAATAAAAGGGACATTCTGGCAATCGTCTG |
| Lm[4][0] | CCGTCTATACCAGTCACACGACCATTACATTG |
| Lm[4][1] | GCAGATTCCAGGGCGATGGCCCACGCGAAAAA |
| Ls[4] | TACGTGAACCACGTCAAAGG |
| L[5][0] | TTCTAGCTGATCATTAAATTTTTGTTAAATCATATTCAACCG |
| L[5][1] | GTTAAAATTCGAAATTAATGCCGGAGAGGGTATTAATATTTT |

| | |
|---|---|
| L[5][2] | AGAGATCTACACAAATATTTAAATTGTAAACGGCTATTTTTG |
| L[5][3] | GATTGTATAAGAAGGCTATCAGGTCATTGCCTAAAACAGGAA |
| L[5][4] | AGCAAACAAGATTGATAATCAGAAAAGCCCCAGAGAGTCTGG |
| L[5][5] | GCAAGGCGATTAAAGCCTGGGGTGCCTAATGAGGGATGTGCT |
| L[5][6] | GCATAAAGTGTAAGTTGGGTAACGCCAGGGTTCGAGCCGGAA |
| L[5][7] | CGACGTTGTAACTCACAATTCCACACAACATATTCCCAGTCA |
| L[5][8] | ATTGTTATCCGAACGACGGCCAGTGCCAAGCTCCTGTGTGAA |
| L[5][9] | CAGGTCGACTCTAATCATGGTCATAGCTGTTTTGCATGCCTG |
| Lm[5][0] | CGTAAAACACCGAGCTCGAATTCGTAGAGGAT |
| Lm[5][1] | CCCCGGGTTAGCATGTCAATCATAACGGTAAT |
| Ls[5] | TGTACCCCGGGAATCGATGA |
| Ef[0] | GGCAGGTCAGACCTCAGAACCGCCACCCTCAGAGGAGGTTGA |
| Ef[1] | AGATATAGAAGATTAAACCAAGTACCGCACTCCAAGCAAATC |
| Ef[2] | AAATCCTGTTTGTGAGACGGGCAACAGCTGATCAGCAGGCGA |
| Ef[3] | GTCAAATCACCTAAAAATTTTTAGAACCCTCAGCCGGAGACA |
| X[0][0] | CATTCCATATAAGGTCACGTTGGTGTAGATGGCTGGAAGTTT |
| X[0][1] | ACCGTGCATCTTATGCAACTAAAGTACGGTGTGCGCATCGTA |
| fX[0][0] | CATGTTTTAAAGCCAGTTTGAGGGGACGACGATGTAGCTCAA |
| fX[0][1] | CTCAGGAAGATCTTAATTGCTGAATATAATGCCAGTATCGGC |
| fX[0][2] | ATGGCTTAGAGCGCACTCCAGCCAGCTTTCCGATTTTTGCGG |
| fX[0][3] | TGGTGCCGGAATGCTCCTTTTGATAAGAGGTCGCACCGCTTC |
| fX[0][4] | GTACCTTTAATACCAGGCAAAGCGCCATTCGCGGATTAGAGA |
| fX[0][5] | GCGCAACTGTTGAAGCAAACTCCAACAGGTCACATTCAGGCT |
| X[0][8] | GAACCAGACCGGGGAAGGGCGATCGGTGCGGGGCTTCAAAGC |
| X[0][9] | ATTACGCCAGCAAATATCGCGTTTTAATTCGACCTCTTCGCT |
| X[1][0] | GAAGAAAAATCTCATTTTCAGGGATAGCAAGCAGGACGTTGG |
| X[1][1] | CCCATGTACCGAACTGGCTCATTATACCAGTCCCAATAGGAA |
| fX[1][0] | GCGATTTTAAGTAACACTGAGTTTCGTCACCAATTACCTTAT |
| fX[1][1] | CAACGCCTGTATTTCAACTTTAATCATTGTGAGTACAAACTA |
| fX[1][2] | AGATGGTTTAAGCATTCCACAGACAGCCCTCAATTGGGCTTG |
| fX[1][3] | AACGATCTAAAAAACACCAGAACGAGTAGTAATAGTTAGCGT |
| fX[1][4] | GCCCTGACGAGGTTTTGTCGTCTTTCCAGACGAATAAGGCTT |
| fX[1][5] | AATTTTCTGTAAACAAAGCTGCTCATTCAGTGTTAGTAAATG |
| X[1][8] | CAAATCAACGTTGGGATTTTGCTAAACAACTTATTCATTACC |
| X[1][9] | CAGCGGAGTGATAATCTTGACAAGAACCGGATTCAACAGTTT |
| X[2][0] | AGTATAAAGCCTCTTACCGAAGCCCTTTTTAAAATTCTTACC |
| X[2][1] | CAGATAGCCGATAGTATCATATGCGTTATACAGAAAAGTAAG |
| fX[2][0] | AAAAGCCTGTTACAAAGTTACCAGAAGGAAACAATTACTAGA |

| | |
|---|---|
| fX[2][1] | CAATAATAACGAAGAATAAACACCGGAATCATCGAGGAAACG |
| fX[2][2] | AGGCGTTAAATGAATACCCAAAAGAACTGGCAGTGATAAATA |
| fX[2][3] | TCCTTATTACGATGGTTTGAAATACCGACCGTTGATTAAGAC |
| fX[2][4] | ACCTAAATTTACAGTATGTTAGCAAACGTAGATCATCTTCTG |
| fX[2][5] | ATAAAGGTGGCTCAAATATATTTTAGTTAATTAAATACATAC |
| X[2][8] | AGAAAACTTTTAACATATAAAAGAAACGCAAAAAAGAACGCG |
| X[2][9] | AATAAGTTTATATGCAAATCCAATCGCAAGACGACACCACGG |
| X[3][0] | TTTAACGTCAGAACGTGCTTTCCTCGTTAGAAGATTTTCAGG |
| X[3][1] | AGCTAAACAGGCAGAAATAAAGAAATTGCGTATCAGAGCGGG |
| fX[3][0] | TGCACGTAAAAAGGCCGATTAAAGGGATTTTACAAAATTATT |
| fX[3][1] | GTACGCCAGAAAAGGGTTAGAACCTACCATATGACAGGAACG |
| fX[3][2] | CTGAATAATGGTCCTGAGAAGTGTTTTTATAAGATTATACTT |
| fX[3][3] | CACCGAGTAAACAATATAATCCTGATTGTTTGTCAGTGAGGC |
| fX[3][4] | TGGCAATTCATAGAGTCTGTCCATCACGCAAATATCAGATGA |
| fX[3][5] | TAGCAATACTTAATTATCATCATATTCCTGATTTAACCGTTG |
| X[3][8] | AGAAGGAGCGGCTTTGATTAGTAATAACATCAAGAAACCACC |
| X[3][9] | TAGAAGAACTCCATTATCATTTTGCGGAACAACTTGCCTGAG |
| mX[0][0] | CATGTTTTAAAGCCAGTTTGACCAGCTTTCCGTGTAGCTCAA |
| mX[0][1] | TGGTGCCGGAACTTAATTGCTGAATATAATGCGCACCGCTTC |
| mX[0][2] | ATGGCTTAGAGACCAGGCAAAGCGCCATTCGCATTTTTGCGG |
| mX[0][3] | GCGCAACTGTTGAAGCAAACTGATAAGAGGTCCATTCAGGCT |
| mX[1][0] | GCGATTTTAAGTAACACTGAGGACAGCCCTCAATTACCTTAT |
| mX[1][1] | AACGATCTAAATTTCAACTTTAATCATTGTGATAGTTAGCGT |
| mX[1][2] | AGATGGTTTAAGTTTTGTCGTCTTTCCAGACGATTGGGCTTG |
| mX[1][3] | AATTTTCTGTAAACAAAGCTGACGAGTAGTAATTAGTAAATG |
| mX[2][0] | AAAAGCCTGTTACAAAGTTACAAGAACTGGCAAATTACTAGA |
| mX[2][1] | TCCTTATTACGAAGAATAAACACCGGAATCATTGATTAAGAC |
| mX[2][2] | AGGCGTTAAATCAGTATGTTAGCAAACGTAGAGTGATAAATA |
| mX[2][3] | ATAAAGGTGGCTCAAATATATATACCGACCGTAAATACATAC |
| mX[3][0] | TGCACGTAAAAAGGCCGATTATGTTTTTATAACAAAATTATT |
| mX[3][1] | CACCGAGTAAAAAGGGTTAGAACCTACCATATTCAGTGAGGC |
| mX[3][2] | CTGAATAATGGAGAGTCTGTCCATCACGCAAAGATTATACTT |
| mX[3][3] | TAGCAATACTTAATTATCATCCTGATTGTTTGTTAACCGTTG |
